# Expanded ME-model analysis capabilities enable genetic recoding, proteostasis, RNA decay, and cofactor essentiality in bacterial models

**DOI:** 10.64898/2026.11.13.765983

**Authors:** Rodrigo Santibáñez-Palominos, Juan D. Tibocha-Bonilla, Diego Tec-Campos, Maxwell J. Neal, Karsten Zengler

## Abstract

Genome-scale metabolic models (M-models) are widely used to understand and manipulate microbial metabolism *in silico*, and are characterized by an extensive availability of automated reconstruction, analysis tools, and simulation software. The scope and accuracy of M-models was greatly improved by the development of metabolic and gene expression models (ME-models) aimed to infer the optimal allocation of individual macromolecules such as proteins, RNA, and enzymatic complexes which constrain metabolic fluxes to their availability and abundance. Although ME-models reconstruction and simulation is currently aided by open-source software such as coralME, COBRAme, ECOLIme, and solveME, they lag behind their M-model counterparts and lack formal analysis tools such as gene essentiality analysis, that would facilitate a broader adoption of ME-models in research. Moreover, the rigid and incomplete formulation of ME-models limit advanced analyses they are theoretically capable of, including the simulation of genomically recoded organisms, protein homeostasis, enzyme cofactor essentiality, and RNA decay. Here, we present coralME v1.3, a vastly improved software version aimed to expand the applicability of ME-models by adding support for translation of non-canonical amino acids, protein and RNA degradation reactions, use of arbitrary coupling coefficients and parameter values, and formalized gene and cofactor essentiality analyses. This work expands, formalizes, and streamlines ME-model reconstruction and analyses for applied research.

## Introduction

Genome-scale metabolic and gene expression models (ME-models hereafter) have been proposed to increase the simulation accuracy of complex bacterial phenotypes over prior methods ^1–4^. While mathematically similar to genome-scale metabolic models (M-models hereafter) as both type of models are constraint-based optimization problems with a stoichiometric matrix (S-matrix) and reaction bounds that encode biochemical reactions and reversibilities, ME-models have demonstrated broader utility than M-models. The first reconstruction of an ME-model totalled 13,694 reactions and established the structure of the gene expression stoichiometric matrix (E-matrix) as a subset of the S-matrix ^1^, but it simplified the metabolic reactions (M-matrix) to only model the transport of extracellular components. By iteratively applying an increasing number of constraints such as rRNA transcription initiation and mRNA decay rates in addition to metabolic uptake rates, the model developed by Thiele et al., 2009, was capable of determining ribosome production rates in close agreement to experimental data. Consequently, ‘reaction coupling’ and ‘coupling coefficients’ were introduced in order to enforce the synthesis of enzymes and tRNAs in proportion to the fluxes through gene expression reactions ^2^, a strategy that was also applied successfully to a *Thermotoga maritima* model resulting in an E-matrix of 206 genes, 18,438 metabolites, and 17,535 reactions ^5^. The first fully integrated ME-model represented 1,937 genes, 62,212 metabolites, and 76,589 reactions ^3^, and since then, multiple ME-models have been reconstructed for *Bacillus subtilis* ^6,7^, *Clostridium ljungdahlii* ^8^, *Escherichia coli* ^9–12^, *Pseudomonas aeruginosa* ^13^, *Pseudomonas putida* ^14^, and *Rhodopseudomonas palustris* ^15^.

Thus far, ME-models have enabled a deeper understanding of the necessary macromolecule expression for optimal bacterial growth, emergence of overflow metabolism, and responses to stressors, including low nutrients, high temperature, low pH, and oxidative damage from reactive oxygen species ^5–18^. Historically, reconstruction and simulation of ME-models have been methodologically time-consuming ^14,16^, and simplified alternatives such as enzyme-constrained M-models (ecGEMs), Resource Balance Analysis, and Protein Allocation Models ^19–21^ have partially remedied reconstruction and simulation time at the expense of model scope and accuracy. These alternative modeling frameworks focus mainly on determining proteome allocation, while ME-models directly model many more components of cellular biomass, such as intracellular metabolic pools, DNA, mRNA, rRNA, tRNA, ncRNA, proteins, enzymatic cofactors, membrane lipids, and peptidoglycan ^17^, making ME-models a more thorough method to understand bacterial gene expression and enzyme activity. To address reconstruction and simulation in a cost-effective way, we have previously developed coralME as an automated pipeline for a reference-based ME-model reconstruction and troubleshooting algorithm ^18^. After validation of coralME v1.0 using published *B. subtilis*, *C. lungdahlii*, *E. coli*, and *T. maritima* ME-models ^5,7,8,16^, the software was capable of reconstructing, correcting, and simulating 17 new, fully functional ME-models for bacteria belonging to eight different phyla (see Supplementary Fig. S4 in ^18^), and 495 new, fully functional ME-models for a diverse set of human gut-associated bacteria belonging to ten different phyla (see Fig. 3 in ^18^).

While reconstruction of ME-models has been automated and readily produces hundreds of models in a few hours ^18^, it still lacks the analysis tools that are common to M-models ^22^ and other metabolic modeling frameworks that complement optimization of wild-type and modified strain phenotypes. Therefore, the simulation of them has mainly focused on standard or modified medium conditions by applying Flux Balance Analysis ^23–25^. For instance, the *Thermotoga maritima* ME-model was simulated in a defined minimal medium with either arabinose or cellobiose as sole carbon sources, determining 142 essential genes out of 206 as a consequence of the carbon source and the necessary upcycling of RNA degradation products and RNA modification byproducts ^5^. Similarly, an *E. coli* ME-model was tested in 170 growth conditions, and a single-gene knockout analysis ^3^ revealed that six non-essential genes in the M-model *i*AF1260 ^26^ were false negatives, which could not have been determined without incorporating macromolecular constraints. The ME-model for *C. ljungdahlii* also was reconstructed and simulated in defined medium (CO, CO_2_+H_2_, fructose, glucose, pyruvate, and arginine as sole carbon sources) determining the maximum growth rates and overflow of acetate, ethanol, and glycerol ^8^. In addition, Liu et al., 2019, determined the effect of nickel availability on the activity of the Wood-Ljungdahl pathway enzymes.

Metabolic modeling using M-models has long benefited from a wide suite of analysis tools and software ^22^. For instance, COBRApy ^27^ includes functions for the structural analysis of M-models ^28^, including the determination of essential genes. COBRAme ^16^ can only approximate this by using the ‘remove_genes_from_model’ function, which simply removes user provided genes, and requires further user scripting to then test the feasibility of the knockout and repeat systematically. Multiple studies using ME-models have relied on the prediction of essential genes and subsequent comparison to experimental data ^1,3,5,7,14^. This has been approached in other ME-models by setting translation reaction bounds to zero ^1,3^, which is also absent from COBRAme’s functions. Moreover, ME-models lack modeling flexibility for three main reasons: they disregard rRNA, tRNA, and protein degradation that are established to be important during normal and stress growth conditions ^29,30^, they employ coupling coefficients whose formulas are evaluated only once during model reconstruction ^17^ and are hard to modify after model reconstruction, and they lack methods to set new linear and non-linear constraints.

Here, we present a widely improved version of the open-source coralME software that expands the ME-model framework to model rRNA, tRNA, protein degradation, facilitates the addition of linear and non-linear flux constraints, and automates the systematic analysis of essential genes and enzymatic cofactors (‘single_gene_essentiality_analysis’ and ‘single_cofactor_essentiality_analysis’ functions). Finally, the ME-model reconstruction algorithm was extended to consider recoded internal stop codons ^31,32^ and it includes convenient reconstruct options such as the possibility to use the output of tRNAscan ^33^ to complete missing tRNA annotations. These additions and modifications significantly improve the robustness of coralME as a software for the reconstruction, simulation, and analysis of ME-models for wild-type and engineered bacteria.

## Results

### coralME v1.3 extends the ME-model framework

The formalization of the ME-model structure was originally defined in two submatrices, the M- and E-matrices ^17^. In the M-matrix, the M-model metabolic network is modified to include complexes in the left-hand side of the reactions associated with a ‘dilution coefficient’ α (Fig. 1, top) proportional to the growth rate and inversely proportional to the enzyme complex catalytic efficiency (*keff*). The E-matrix contains reactions modeling transcription, charging of tRNA, translation, and formation of complexes, whose metabolites can be associated with either ‘dilution coefficients’, α, or ‘degradation coefficients’, β (Fig. 1, middle-left panel). Although ME-models consider the major mechanisms in gene expression, the current ME-model framework ^16,18^ disregards protein degradation and most forms of RNA degradation. To incorporate these mechanisms, we extended and modified the computational structure of ME-models to accommodate rRNA, tRNA, ncRNA, and protein degradation reactions (Fig. 1, middle-right and bottom panels). For rRNA degradation, we created the reaction class *rRNAFormationReaction* and extended its stoichiometry to consume rRNA biomass and produce the correct amount of nucleotides (Fig. 1, bottom-left panel). Because rRNA modifications can be optionally added in transcription reactions ^16^, rRNA degradation was implemented for unmodified sequences for sake of simplicity. To represent tRNA degradation, tRNA charging reactions were modified by adding a corrected stoichiometry on the right-hand side of the reactions, and a similar assumption that tRNAs are degraded before being modified (including aminoacylation) is integrated to simplify the stoichiometric calculation. In addition to the main three RNA classes, ME-models include two ncRNA components, the ffs component of the signal recognition particle and the rnpB component of the RNase P, which are requirements for translocation and the processing of pre-tRNAs in transcription reactions, respectively. In these two cases, the RNA degradation reactions were added directly to their respective transcription reactions. The degradation of rRNAs and tRNAs can be similarly incorporated into their specific, existing transcription reactions.

**Figure 1.**
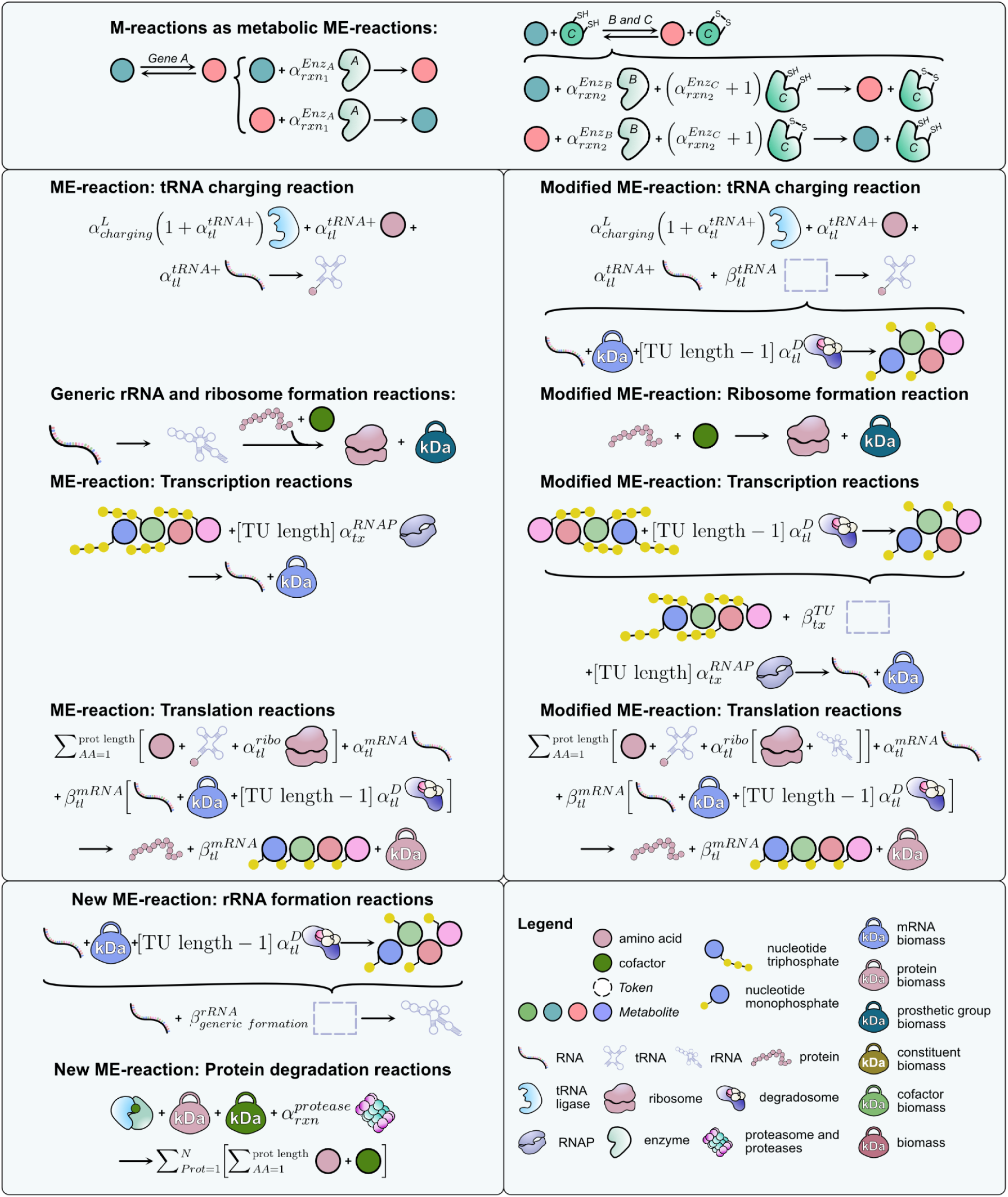
Overview of changes implemented in the updated ME-model framework. **Top panel,** metabolic reactions are implemented without changes, where enzymes are modeled in the left-hand side with proper dilution coefficients. **Middle-left and middle-right panels,** schematic description of the implemented changes to transcription, tRNA charging, the ribosome formation, and translation reactions. Degradation of RNA is extended from previous description affecting only mRNA in translation reaction to currently include stable RNA (rRNA and tRNA) and transcriptional units. **Bottom-left panel,** new reactions added for the formation of generic rRNA coupled to its degradation and protein degradation (including macromolecular complexes). Degradation of proteinic components can be stoichiometrically coupled to metabolic and other reactions. Degradation of rRNA and tRNA are assumed to be carried out by the cellular degradosome complex and coupled to their corresponding formation and charging reactions, respectively. Note that the ribosome in translation reactions is a protein-only complex, requiring a reformulation of the ribosome formation reaction. Additional components such as ATP, GTP, and water were omitted. Abbreviations: α’s define dilution coefficients and β’s define degradation coefficients. D, degradosome; L, tRNA ligase; tRNA+, charged aminoacyl-tRNA; ribo, ribosome; RNAP, RNA polymerase; rxn, reaction; tx, transcription; tl, translation; kDa: biomass constraints. AA, amino acid, N, stoichiometric coefficient of a protein in a complex.

Notably, the user now has full control of the parameter values that determine the flux through any degradation reaction (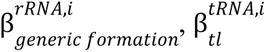, and 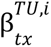, Fig. 1 and Supplementary File S1). Similarly, we modified the ME-model structure so that each mRNA decay is controlled by an independent, tunable rate 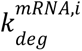. To keep retrocompatibility to the original ME-model framework ^16^, the dimensionless degradation coefficients 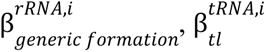, and 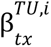 are instantiated as zero and the degradation rate for mRNAs in translation reactions 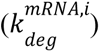 are instantiated at the *E. coli* default value ^16,17^.

Similarly, coralME now creates independent degradation reactions for proteins and protein complexes catalyzed by the bacterial proteasome and proteases, including careful consideration of covalent modifications (e.g., biotin, lipoate, acetate, and dipyrromethane) that when removed produce modified amino acids (e.g., biocytin, N(6)-lipoyl-L-lysine, N-acetyl-L-alanine, and dipyrrolylmethanemethyl-L-cysteine), informing the user of the need for further reactions to keep the stationary state. Two parameters control the energy consumption, defaulting to 9 ATP per proteasome cut every 15 amino acids ^34^. These protein degradation reactions cover most of the complexes in an ME-model, but the process is complicated by the ability of coralME to create generic components. As these components represent several structural versions of a complex, their degradation products must also vary. This problem is notable in lipoproteins, where multiple forms using different lipids are not individualized, complicating the determination of which free lipids will be produced from the complete degradation of such complexes. Generics are essential in ME-models to reduce combinatorial complexity, but they confound the calculation of complex composition. To demonstrate this, a toy ME-model is presented (Supplementary File S1) containing 263 degradation reactions for its 264 complexes. The single missing reaction is for the holo acyl-carrier protein that is degraded by a phosphodiesterase reaction, present in M-models as ACPpds and in ME-models as a metabolic reaction.

To improve the ability of ME-models to consider any new biochemical reactions, we replaced set parameter values throughout the model with fully symbolic expressions as ‘coupling coefficients’. Previously, COBRAme ^16^ encoded dilution coefficients as fixed values determined from other, fixed parameter values and the growth rate (Fig. 2, top-left panel). Thus, few ‘coupling coefficients’ in the ME-models could be easily modified after reconstruction. For instance, enzyme *keffs* needed to be modified not only by setting a new value, but also requiring the user to update the associated metabolic reaction stoichiometry. On the other hand, modification of the ‘coupling coefficients’ for RNAP, ribosomes, tRNAs, tRNA ligases, and mRNAs required an iterative, direct, and error-prone manipulation of the reaction stoichiometries. To overcome this methodological problem and enable arbitrary parameter values as set by the user, dilution and degradation coefficients are now symbolic functions that are evaluated at the time of simulation (Fig. 2, top-left panel). The new parameter dictionary (*model.default_parameters*) declares variables and their values to use during optimization of the growth rate, allowing the user to redefine values that diverge from the *E. coli* defaults ^16^. As a consequence, parameter sensitivity analysis can be performed by setting new values in the parameter dictionary, immediately followed by optimization and assessing impact on the growth rate. The new parameters module contains functions allowing the user full control and creation of ME-model parameters, new symbolic expressions (*model.symbols*) and ultimately, new ‘coupling coefficients’ from these symbolic expressions that can be set model-wide or into specific reactions. To reduce the probability of using incorrect coupling coefficients, symbols can be associated with physical units, allowing them to self-check for dimension agreement. The manipulation of reactions stoichiometries has been further improved through the use of *MCounter* and *ScalableKeyDict*. Previously, metabolites can be added or removed from reactions by using the ‘add_metabolites’ method and carefully setting new stoichiometric values. However, python dictionaries do not allow simple addition or scalar multiplication, limiting the ways the user can manipulate reaction stoichiometries. Now, reactions can be added to other reactions as if they were algebraic equations, and reactions can be scaled up or down by a scalar, including the use of symbolic expressions (Supplementary File S1).

**Figure 2.**
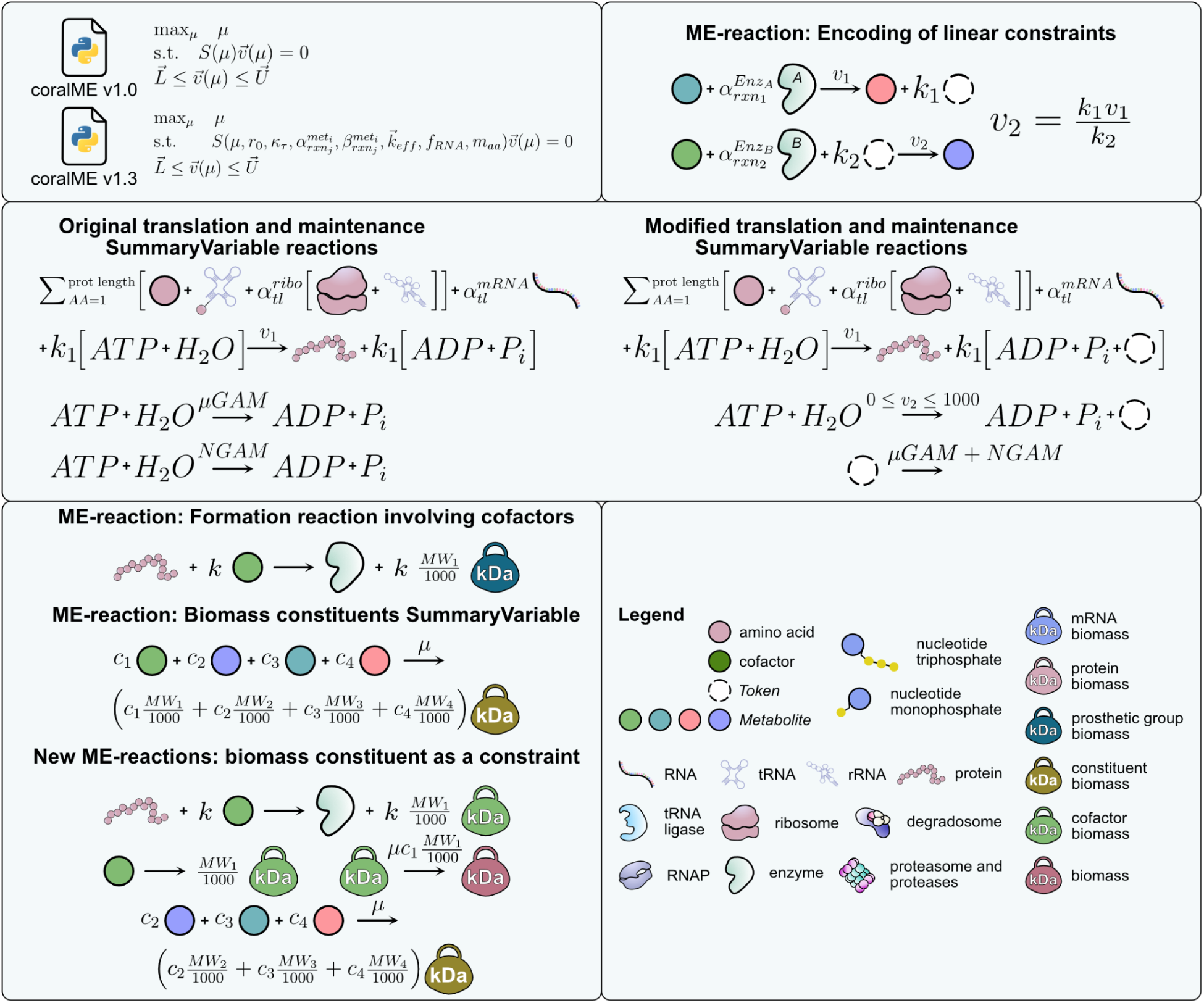
Encoding of linear and non-linear constraints using virtual metabolites. **Top-left panel,** parametric representation of the ME-model structure before simulation. **Top-right panel,** creation of a virtual metabolite (white circle) and its pairing in opposite sides of multiple equations permits the addition of flux constraints of the form *v_1_ = kv_2_*, with *k* being a proportionality linear or non-linear constraint into ME-models. **Middle panel, determination of GAM and NGAM correction values in a modified ME-model.** Left, simplified translation reaction showing explicitly ATP hydrolysis where the stoichiometry coefficient *k_1_* is equal to three times the protein length. GTP and other elements are omitted for simplicity (see Fig. 1). The GAM reaction was modified to transfer the GAM value from the stoichiometry to the bounds of the reaction. Right, modifications imposed into the ME-model for the calculation of an explicit GAM+NGAM value (e.g., right-hand side of a translation reaction), where a *token* metabolite quantifies the amount of ATP and GTP hydrolysis. The GAM reaction is modified to produce the implicit GAM+NGAM value, while the NGAM reaction is inactivated or removed from the ME-model. A new constraint in the ME-model is imposed by modifying the bounds of a demand reaction to match the total GAM+NGAM value during optimization. **Bottom-left panel,** addition of non-linear constraints based on biomass components. The example shows the formation of a complex from a protein and a cofactor (*k* is a positive integer), cofactor that is also in the biomass constituent reaction (*c_1_* is in mmol per gDW), leading to overestimation if the enzyme is essential for growth. To encode *c_1_* as a constraint, three modifications should be implemented: the formation reaction should produce a new biomass component (green kettlebell) instead of the generic *prosthetic group* biomass (blue kettlebell), the addition of a reaction converting the free cofactor into its biomass, and the addition of a reaction to convert this biomass component into biomass (light red kettlebell). The latter reaction encodes the *c_1_* constraint in its lower bound. Abbreviations: µ, growth rate; tl, translation; ribo, ribosome.

### coralME v1.3 facilitates the addition of flux constraints

The simulation of ME-models requires tuning the ‘coupling coefficients’ that scale the interdependencies between metabolism and expression and specialized linear programming (LP) solvers to calculate feasible flux values, which span many orders of magnitude ^1,16–18^. To solve ME-models, the COBRAme and coralME software interface with the quad MINOS solver ^35,36^ as a default solver. Additional constraints are not allowed except for the accumulation or depletion of specified metabolites by adding sink or demand reactions. To add additional complex constraints to an M-model, COBRApy ^27^ currently relies on optlang ^37^, but this dependency was removed to accelerate ME-model reconstruction ^18^. Here, we extended coralME to partially recover the capacity to encode linear and non-linear constraints (Fig. 2, top-right panel), and expand the utility of ME-models to scenarios where fluxes are functions of others. To do so, a virtual metabolite or *Token* is created and added to the left- and right-hand sides of the reactions to be stoichiometrically coupled. If the stoichiometric coefficients are identical, the flux ratio is 1, and modifying them the flux ratio is modified as well (Fig. 2, top-right panel). It is noteworthy to mention that the stoichiometric coefficient for the *Token* metabolite can be any symbolic expression, including constant values, as long as it is dimensionless.

The use of *Tokens* also simplifies the representation and calculation of coupled fluxes present in earlier formulations. For instance, *Tokens* allow us to bypass the combination and deletion of reactions previously deployed for mRNA degradation in translation reactions (Fig. 1). This is especially expedient in the recalculation of the GAM and the NGAM values (Fig. 2, middle panel). Previously, M-models use experimental data to determine the amount of energy associated to growth and non-growth biological processes ^38^, values that ME-models can reproduce partially from ATP and GTP hydrolysis through explicitly modeled energy consumption in translation, transcription, tRNA charging, and posttranslation reactions. The manipulation of the ME-model to add a *Token* in the right-hand side of translation reactions allows the LP solver to quantify the consumed ATP at each growth rate (explicit GAM+NGAM), while adding the same *Token* in the right-hand side of a non-enzymatic ATP hydrolysis reaction and using a new constraint derived from GAM and NGAM values automatically calculates the remainder not accounted for (implicit GAM+NGAM). The complete computational procedure is described in Supplementary File S1.

Another utility of *Token* metabolites for the encoding of complex constraints is the use of biomass components as limitations on enzyme activities. In M-models, enzyme cofactor usage is represented by fixed values in the biomass composition, opening the possibility for their underestimation or overestimation when they are also used as enzyme cofactors in ME-models. When this biomass composition is imported to ME-models, this leads to overestimation of cofactor usage if ME-models also predict the use of magnesium, zinc, iron, and other metals that are essential for metabolism in formation reactions (e.g., ribosome in Fig. 1), in addition to the amounts mandated by the biomass reaction. As a rule of thumb, enzyme cofactors are omitted from the biomass reaction ^17^, but this overcorrects and leads to the underestimation of non-essential metals such as calcium and sodium as a consequence. Here, we propose a methodology to incorporate such constraints into ME-models (Fig. 2, bottom-left panel). In the current ME-model framework, the consumption of enzyme cofactors (e.g., magnesium) in formation reactions contributes to a virtual metabolite called *prostethic_group_biomass*, in grams by using a conversion factor equal to the corresponding molecular weight (MW) divided by 1000. Similarly, the same enzyme cofactor in the ME-model biomass reaction contributes to a virtual metabolite called *constituent_biomass*. To use an enzyme cofactor as a constraint, a new *Token* metabolite is added to the ME-model that quantifies how much biomass is used in formation reactions for that specific cofactor (i.e., Σ *k_i_* MW*_cofactor_*/1000) separate from the *prostethic_group_biomass*, where *k_i_* is the stoichiometric coefficient for the cofactor in the formation reaction *i*. Then, two new reactions are added, the first to freely convert the cofactor into biomass (e.g., *magnesium_c_biomass*) in grams and a second reaction to dilute into the overall biomass calculation with a lower bound equal to the proportion of the biomass that the cofactor represents (i.e., *μc*_1_MW_1_/1000). For convenience, the user can use the new *flux_of_biomass_constraints* reconstruction dictionary instead of *flux_of_biomass_constituents*, letting coralME v1.3 automate the creation of these metabolites, reactions, and bounds. The complete computational procedure is described in Supplementary File S1.

### Efficient modeling of genomically recoded organisms using coralME v1.3

The reconstruction of ME-models was originally formalized for *E. coli* where selenocysteine (Sec) can be added to internal stop codons (TGA), a hallmark of the bacterial genetic code. To detect and implement this modification in translation reaction stoichiometries, coralME uses protein homology to detect selA and selB genes in the input genome and creates a translation subreaction that uses SelA and SelB complexes as enzymes. The reaction is completed if the selC gene (tRNA^(Sec)^) is detected in the genome by its name. For all the other canonical amino acids, coralME creates tRNA charging reactions pairing an amino acid to the corresponding tRNA and tRNA ligase, assigned automatically or from a user input file. To model recoded organisms ^31,32,39–41^, we exploited this feature to instruct coralME v1.3 to create tRNA charging reactions for arbitrary tRNA-amino acid pairs, and redesigned coralME to detect internal stop codons (Fig. 3, top panel). To identify genes with internal stop codons, *TranslationData* objects contain information in the notes property.

**Figure 3.**
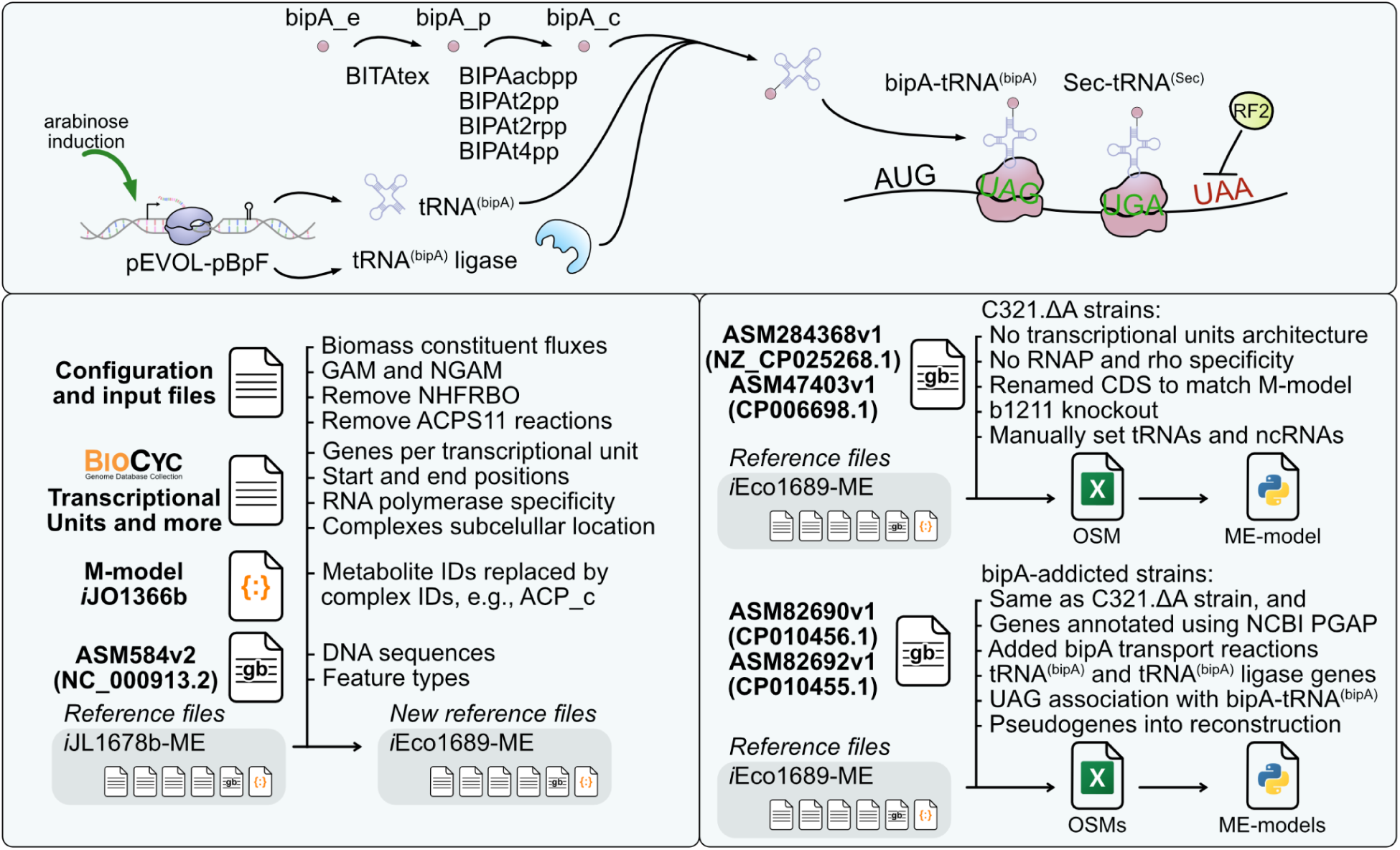
Schematic reconstruction of reference *i*Eco1689-ME and GRO models. **Top panel,** schematic representation of the manual curation to incorporate a new bipA-tRNA^(bipA)^ synthesis (charging tRNA reaction) and translation of UAG codons in pseudogenes. Transport reactions associated with alanine are assumed to transport bipA from the periplasm using ATP hydrolysis (BIPAabcpp), proton symport (BIPAt2pp and BIPAt2rpp), or sodium symport (BIPAt4pp). Induction of arabinose genes is not modeled. The BIPAtex reaction transports bipA from the extracellular medium by diffusion. **Bottom-left panel,** description of input files used for the reconstruction of *i*Eco1689-ME model from reference files included in coralME. The reference files to reconstruct *i*Eco1689-ME were previously manually curated ^18^, and troubleshooting was performed to ensure correctness. **Bottom-right panel,** reconstruction of a ME-model for the engineered strain C321.ΔA ^31^ from reference files with modifications, and reconstruction of two ME-models for the bipA-addicted strains that use bipA to rescue the expression of the tyrS and serS genes encoding the essential L-tyrosine:tRNA(Tyr) ligase (AMP-forming) and L-serine:tRNASer ligase (AMP-forming) enzymes, respectively.

The general procedure for the reconstruction of ME-models for recoded organisms (hereafter GRO-models) consists of four, manual curation steps (Fig. 3, top panel). The first step is the addition of transport reactions for a non-canonical amino acid (e.g., bisphenylalanine, hereafter bipA), that can be done to the M-model before generation of input files or to the *reaction_corrections.txt* file before ME-model reconstruction. If using *reaction_corrections.txt*, new metabolites in new reactions can be added in the *metabolite_corrections.txt* input file to complete formula and charges. The second step is the pairwise declaration of an amino acid ID and its tRNA ligase in the *amino_acid_trna_synthetase.txt* input file. If there are multiple tRNA ligases, a generic complex should be used and added to the *generic_dict.txt* input file to denote complexes equivalent in function (e.g., *generic_lys_ligase*). In the third step, the *genetic_recoding* reconstruction flag indicates the TAG codon is decoded by the product of an amino acid and a tRNA (e.g., “{ ‘TAG’ : { ‘bipa L_c’ : ‘trna-bipA’ }”). Failure to do this will instruct coralME v1.3 to stop translation and the amino acid sequence will be truncated. Finally as a fourth step, the GenBank file (see Fig. 1 in ^18^) is modified to include locus tags for the tRNA ligase(s) and the tRNA gene(s) (e.g., ‘trna-bipA’), and locus tags are modified to match the M-model GPRs or vice versa. Modifying the GenBank locus tags is preferred in order to agree with the other reference files, avoiding additional, extensive manual curation.

To test the reconstruction of GRO-models and replicability of the methodology, we downloaded five public assemblies for different *E. coli* strains ^31,32,40,42^ and derived a reconstruction protocol from the reference ME-model included in coralME (Fig. 3, bottom-left and bottom-right panels). The reconstruction was initiated using a modified *i*JO1366 M-model ^16^ with non-metabolite IDs replaced by the corresponding complex ID [e.g., *actACP_c* is replaced by *EG50003-MONOMER_mod_pan4p(1)_mod_act(1)*] and GenBank locus tags modified to be identical to the wild-type assembly (ASM584v2), decided on a best bidirectional hit approach. At this point, coralME v1.3 generates input files tailored to the M-model, and missing genes in the GenBank will produce metabolic reactions associated with a dummy complex. After the generation of input files for the first time, information from a reference ME-model should replace these newly created files: *enzyme_reaction_association.txt*, *excision_machinery.txt*, *generic_dict.txt*, *lipid_modifications.txt*, *metabolite_corrections.txt*, *peptide_release_factors.txt*, *protein_complexes.txt*, *protein_modification.txt*, *reaction_corrections.txt*, *ribosomal_proteins.txt*, *ribosome_subreactions.txt*, *rna_degradosome.txt*, *special_modifications.txt*, *special_trna_subreactions.txt*, *transcription_subreactions.txt*, and *translocation_multipliers.txt*. The use of all of these reference files ensures that the reconstruction of GRO-models follows the same namespace for protein complexes and avoids the creation of functional complexes derived from non-functional pseudogenes. Models for the genomic recoded organisms can be reproduced using Supplementary File S2, File S3, and File S4.

The reconstruction of GROs from five genomic sequences resulted in five fully functional GRO-models with distinct numbers of genes, reactions, metabolites, and maximum growth rates (Table 1). The total number of genes is maximal for the wild-type assembly and minimal for the *rE.coli-57* genome. It is worth noting that the wild-type ME-model in this work contains less number of genes than the reference ME-model (1,663 vs 1,689 genes) as we deliberately omitted tRNA modifications to simplify ME-model reconstruction, and as a consequence, 37 associated enzymes were removed from the ME-model during reaction pruning. Moreover, as a consequence of the omission of the TU input file, the reconstruction also removes six of the seven sigma factors and the rho transcription factor from the ME-model, and the gene b2092 (galactitol-specific PTS enzyme IIC component) that is annotated as a pseudogene in ASM584v2. On the other hand, multiple ribosomal accessory proteins were detected and added to the ribosome complex, as well as Lon protease and ClpXP, FtsH/HflKC, and HslUV protease complexes totalling 19 new genes. In the case of the chemically synthesized *rE.coli-57* genome, it was initiated from the MDS42 strain where 15% of the genome was previously removed ^43^. As expected, its GRO-model contained the smallest number of total genes (1,505 vs 1,528-1,571 genes in all the other GRO assemblies). The reduced number of genes is also a product of the higher number of pseudogenes in the metabolic and gene expression networks (84 vs 23-30 pseudogenes in all the other GRO assemblies).

**Table 1.**
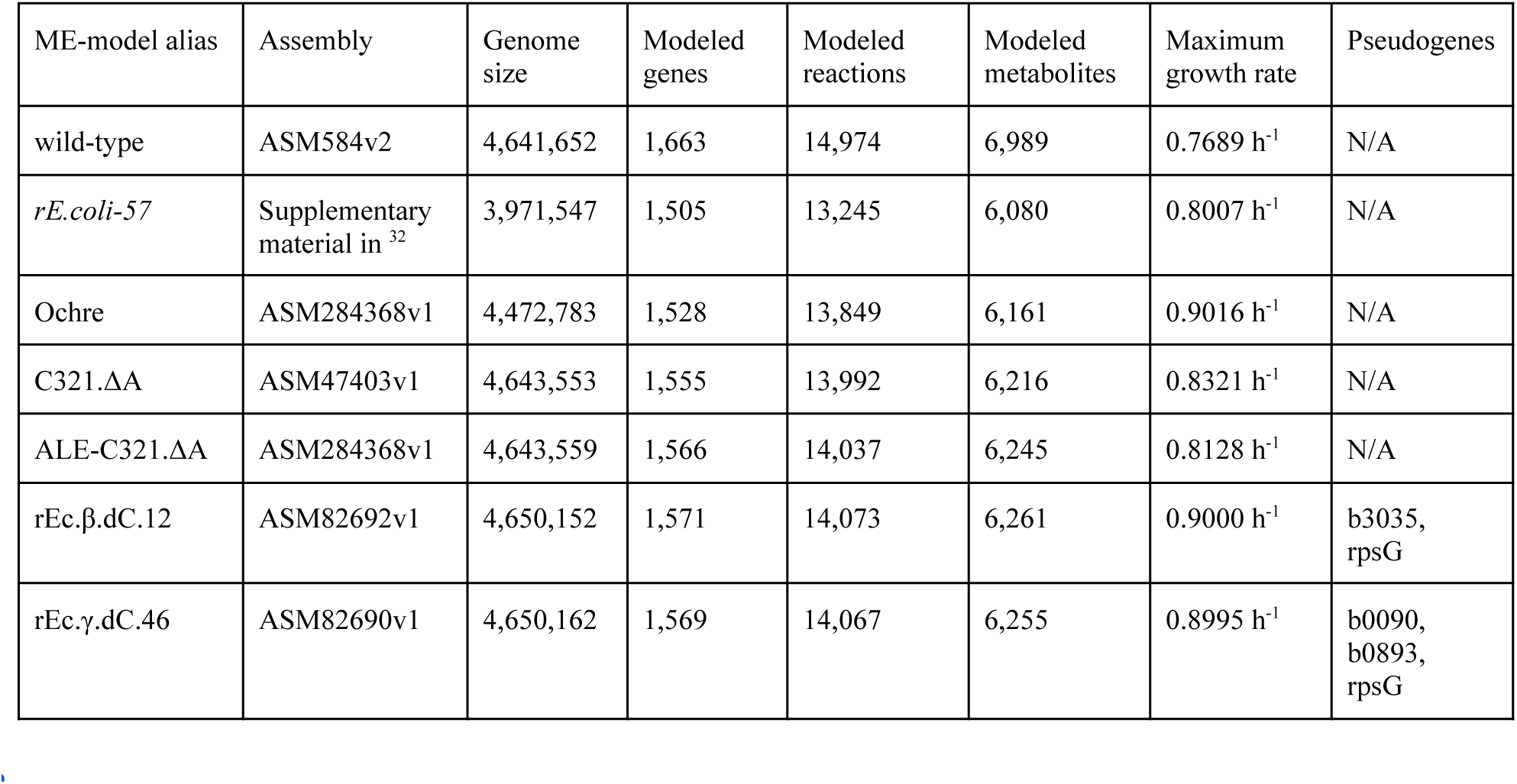
Properties for five GRO-models and their reference wild-type ME-model.

| ME-model alias | Assembly | Genome size | Modeled genes | Modeled reactions | Modeled metabolites | Maximum growth rate | Pseudogenes |
|---|---|---|---|---|---|---|---|
| wild-type | ASM584v2 | 4,641,652 | 1,663 | 14,974 | 6,989 | 0.7689 h<sup>-1</sup> | N/A |
| *rE.coli-57* | Supplementary material in<sup>32</sup> | 3,971,547 | 1,505 | 13,245 | 6,080 | 0.8007 h<sup>-1</sup> | N/A |
| Ochre | ASM284368v1 | 4,472,783 | 1,528 | 13,849 | 6,161 | 0.9016 h<sup>-1</sup> | N/A |
| C321.ΔA | ASM47403v1 | 4,643,553 | 1,555 | 13,992 | 6,216 | 0.8321 h<sup>-1</sup> | N/A |
| ALE-C321.ΔA | ASM284368v1 | 4,643,559 | 1,566 | 14,037 | 6,245 | 0.8128 h<sup>-1</sup> | N/A |
| rEc.β.dC.12 | ASM82692v1 | 4,650,152 | 1,571 | 14,073 | 6,261 | 0.9000 h<sup>-1</sup> | b3035,<br>rpsG |
| rEc.γ.dC.46 | ASM82690v1 | 4,650,162 | 1,569 | 14,067 | 6,255 | 0.8995 h<sup>-1</sup> | b0090,<br>b0893,<br>rpsG |

To control for the effects of variable gene sequences on the catalytic efficiency of enzymes (*keff*’s) by using the SASA method ^16,18^, the same *keff* values were used for all the six ME-models. However, differences in maximum growth rates were still observed (Table 1), indicating that sequence variability can have a significant impact. For comparison, five M-models were derived from the same model used in COBRAme, *i*JO1366 ^44^ by knocking out pseudogenes, resulting in all of them predicting a maximum growth rate of 0.98 h^-1^, except rEc.γ.dC.46 (Table 2). The assembly for rEc.γ.dC.46 shows an internal stop codon in b0090 (N-acetylglucosaminyl transferase), impeding flux through the essential reaction undecaprenyldiphospho-muramoylpentapeptide beta-N-acetylglucosaminyltransferase (EC 2.4.1.227). The mutation in b0090 is rescued by bipA in the corresponding ME-model by consuming 2.78e-6 mmol gDCW^-1^ h^-1^ (0.672 µg of bipA gDCW^-1^ h^-1^).

**Table 2.** Properties for derived M-models using genomic assemblies.

| M-model alias | Assembly | Genome size | Modeled genes | Modeled reactions | Modeled metabolites | Maximum growth rate | Total pseudogenes |
|---|---|---|---|---|---|---|---|
| wild-type | ASM584v2 | 4,641,652 | 1,367 | 2,583 | 1,805 | 0.9824 h<sup>-1</sup> | N/A |
| *rE.coli-57* | Supplementary material in<sup>32</sup> | 3,971,547 | 1,288 | 2,518 | 1,805 | 0.9823 h<sup>-1</sup> | 84 |
| Ochre | ASM284368v1 | 4,472,783 | 1,344 | 2,551 | 1,805 | 0.9720 h<sup>-1</sup> | 58 |
| C321.ΔA | ASM47403v1 | 4,643,553 | 1,338 | 2,537 | 1,805 | 0.9824 h<sup>-1</sup> | 30 |
| ALE-C321.ΔA | ASM284368v1 | 4,643,559 | 1,347 | 2,549 | 1,805 | 0.9824 h<sup>-1</sup> | 23 |
| rEc.β.dC.12 | ASM82692v1 | 4,650,152 | 1,347 | 2,549 | 1,805 | 0.9824 h<sup>-1</sup> | 25 |
| rEc.γ.dC.46 | ASM82690v1 | 4,650,162 | 1,344 | 2,551 | 1,805 | 0.0000 h<sup>-1</sup> | 26 |

### coralME v1.3 allows for out-of-the-box structural analyses of ME-models

A toy ME-model was developed for the purposes of analyzing model behavior when adding new constraints (Fig. 2), and performing single cofactor and single gene knockouts (Fig. 4). The *i*Mmy281-ME model for *Mycoplasma mycoides* has 1,241 metabolites, 2,168 reactions, and 281 genes (6 rRNAs, 29 tRNAs, 244 mRNAs, 2 ncRNAs), an increment of 302, 575, and 89% respectively from a curated M-model ^34^. The reconstruction and structural analysis of the ME-model can be reproduced from Supplementary File S1 and File S5, respectively. The unmodified ME-model predicts a growth rate of 0.1140 h^-1^, 31% of the maximum growth of the constrained M-model ^34^ (See Methods). While this indicates that inferred enzyme efficiencies (*keffs*) using the SASA method are incompatible with experimental growth rate values, we proceeded with single cofactor and single gene essentiality analysis as they are independent of the *keff* values at the tested growth rate of 0.01 h^-1^ (approximately a 1% of the maximum growth rate for the constrained ME-model).

**Figure 4.**
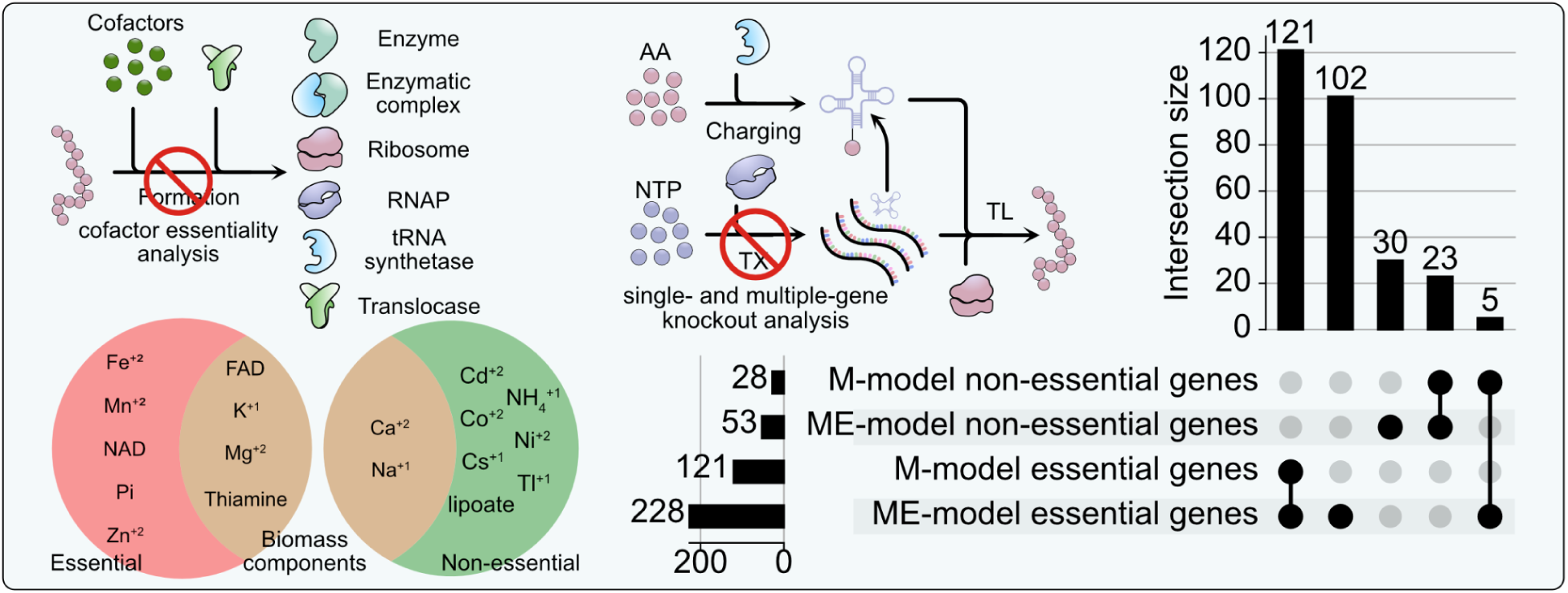
Single cofactor and single gene essentiality analyses of a toy ME-model. **Left panel,** schematic representation of cofactor essentiality analysis that targets formation reactions in a ME-model rather than the synthesis or uptake of the cofactor. For a toy ME-model representing the metabolic network of *Mycoplasma mycoides*, detecting 18 out of 19 cofactors that were classified as essential (9) or non-essential (9). Only 6 are also biomass components (4 essential/2 non-essential), highlighting the robustness of the approach. **Right panel,** schematic representation of gene essentiality analysis that targets transcription reactions. Analysis of M- and ME-models showed high agreement (121 essential and 23 non-essential, 144 out of 149 genes). New genes in the ME-model (102 essential and 30 non-essential), as well as disagreements (5 genes) were detected in a single-gene genome-scale essentiality analysis.

For the analysis of essential cofactors, a total of 19 metabolites were identified (Fig. 4, left panel). Of them, metals (Fe^+2^, K^+1^, Mg^+2^, Mn^+2^, and Zn^+2^) and vitamins (FAD, NAD, and thiamine) were identified as essential cofactors in enzymatic reactions. Phosphate also appears as an essential cofactor for the ribose-phosphate diphosphokinase ^16^. Non-essential metals (Ca^+2^, Cd^+2^, Co^+2^, Cs^+1,^ Na^+1^, Ni^+2^, and Tl^+1^) are mostly associated with the methionine adenosyltransferase enzyme. The model allows any annotated cofactor combination to be used, and the optimal solution was obtained by using ammonia and zinc as cofactors for the essential methionine adenosyltransferase reaction. Moreover, the gene was experimentally determined to be essential, but the M-model failed to predict essentiality (see Table 2 in Breuer et al., 2019). The product of the reaction, S-adenosyl-L-methionine, participates as an rRNA modifier in methylation reactions, which M-models do not capture. The last non-essential identified cofactor, lipoate, is a prosthetic group for the pyruvate dehydrogenase (PDH) enzyme. The PDH reaction was determined to be non-essential on both M- and ME-models. Finally, acetylation is an essential modification of the 50S ribosomal subunit protein L7 ^45^, but the single-cofactor essentiality analysis failed to detect acetate as a cofactor because the modification uses acetyl-CoA as an acetyl donor.

Next, for a single-gene essentiality analysis, the ME-model for *Mycoplasma mycoides* reported a high agreement to its M-model counterpart, despite the latter having only 53% of the total genes of the ME-model. Together, both models agree on 121 essential and 23 non-essential genes (Fig. 4, right panel). The additional reactions in the ME-model predict 102 essential and 30 non-essential genes. Some of these genes were introduced as metabolites into the M-model to represent tRNA and other complexes in reactions (e.g., GLNTRAT reaction converts Glu-tRNA^(Gln)^ into Gln-tRNA^(Gln)^), explaining why they are essential in the ME-model. These genes failed to be detected as essential in the M-model because they were not included in gene-protein-reaction (GPR) rules, and include rRNAs, tRNAs, ribosomal proteins, and RNAP subunits. For instance, ferredoxin that normally appears as a metabolite in M-model reactions should be also included in the GPR rule. Unique ME-model genes also include rRNA modification enzymes and tetM, a unique ribosomal factor added during synthesis of JCVI-syn1.0 for selection purposes ^46^. Initially, the gene JCVISYN3A_0447 was falsely identified as non-essential in the ME-model. It is a component of the biomass reaction in the M-model, but the associated reaction dUTP diphosphatase is not essential. Once JCVISYN3A_0447 was corrected, five genes (JCVISYN3A_0381, JCVISYN3A_0432, JCVISYN3A_0886, JCVISYN3A_0732, and JCVISYN3A_0747) are reported as false non-essential genes if compared to the M-model prediction (Fig. 4, right panel), encompassing the necessity of structural analysis for ME-model quality assurance. The case for essentiality of JCVISYN3A_0381 (5’-methylthioadenosine nucleosidase) and JCVISYN3A_0432 (methionine adenosyltransferase) can be explained for the new requirement of rRNA methylation and upcycling of the byproducts in the ME-model. However, transposon mutagenesis determined that JCVISYN3A_0381 is not an essential gene, indicating an unidentified redundancy in the genome ^34^. JCVISYN3A_0886 (proton-glutamate symporter) was also essential in the ME-model but not in the M-model, seemingly due to a modeling error in the latter, where the *Protein_transl* and *Protein_degrad* reactions are able to cycle to falsely produce amino acids consumed in the *BIOMASS* reaction. This gene was determined as quasi-essential in a transposon mutagenesis experiment ^34^. Similarly, the essentiality of JCVISYN3A_0732 (deoxyribose-phosphate aldolase) and JCVISYN3A_0747 (purine-nucleoside phosphorylase) only in the ME-model could result from a similar modeling error in the M-model where the *RNA_degrad* reaction produces nucleoside monophosphates as intermediates for other reactions, but this pseudoreaction added in the M-model for convenience was removed from the ME-model. The transposon mutagenesis experiments determined JCVISYN3A_0747 to be quasi-essential while JCVISYN3A_0732 was non-essential ^34^.

The ME-model reports 84% accuracy if quasi-essential (QE) genes ^34^ are considered essential, ∼75% if QE genes are considered non-essential, and ∼86% if QE genes are excluded from the analysis (Table 3). The reduction of specificity (TN/(TN+FP)) resulted from the increase from zero to eight genes falsely predicted to be essential (FP), which was not compensated by the increase from 12 to 15 truly non-essential genes (TN). A similar reduction of sensitivity (TP/(TP+FN)) resulted from the increase from 18 to 38 genes falsely predicted to be non-essential (FN) not compensated by the increase from 123 to 219 genes correctly predicted to be essential (TP). The eight false essential genes are two ribosome subunits (rpmG and tetM), rRNA modification enzymes (ksgA, rluD, rlmB, and rsmG), and genes implicated in upcycling of byproducts (mtnN and deoC). The ME-model predictive capability is less impacted if only genes in the M-model are considered (Table 3). Here, only mtnN (JCVISYN3A_0381) and deoC (JCVISYN3A_0732) are false essential genes, implying that accumulation of the byproducts they clear might be tolerated *in vivo* and/or rRNA modifications are not necessary for full ribosomal activity. On the other hand, a reduction from 18 to 14 false non-essential genes and from 12 to 9 true non-essential genes was observed, while only one gene (JCVISYN3A_0432, methionine adenosyltransferase) is correctly predicted as a true essential gene in addition to 123 true essential genes as in the ME-model, totalling 124 out of 149 analyzed genes. Interestingly, the two rRNA operons (JCVISYN3A_0067-0069 and JCVISYN3A_0532-0534) are essential *in vivo*, but the ME-model represents them to be redundant in function. Similarly, tRNA^(Leu)^ genes (JCVISYN3A_0070 and JCVISYN3A_0423) and tRNA^(Lys)^ genes (JCVISYN3A_0071 and JCVISYN3A_0507) were also essential *in vivo* ^34^, but only one copy of each is required for complete ME-model functionality. The full list of predictions are in Supplementary Table S1.

**Table 3.** ME-model performance in comparison to published experimental data.

|  |  | Accuracy | Sensitivity | Specificity | Matthews Correlation Coefficient |
|---|---|---|---|---|---|
| **All 281 modeled genes** | **Quasi-essential as essential genes** | 83.63% (-4.61%) | 85.27% (-1.96%) | 65.22% (-34.78%) | 0.35 (-0.24) |
|  | **Quasi-essential as non-essential genes** | 75.09% (-7.92%) | 87.98% (-8.21%) | 38.36% (-15.81%) | 0.29 (-0.29) |
|  | **No quasi-essential genes** | 85.71% (-10.87%) | 87.98% (-8.21%) | 65.22% (-34.78%) | 0.41 (-0.43) |
| **Common 149 genes** | **Quasi-essential as essential genes** | 89.26% (+1.03%) | 89.86% (+2.62%) | 81.82% (-18.18%) | 0.52 (-0.07) |
|  | **Quasi-essential as non-essential genes** | 80.54% (-2.47%) | 96.19% (+0.00%) | 43.18% (-10.98%) | 0.50 (-0.09) |
|  | **No quasi-essential genes** | 94.83% (-1.75%) | 96.19% (+0.00%) | 81.82% (-18.18%) | 0.72 (-0.12) |

## Discussion

Since the first proposal of a computational, constraint-based genome-scale models designed to describe the molecular steps in the synthesis of macromolecules and gene expression ^1^, degradation of RNA molecules has been circumscribed to only consider messenger RNAs due the assumption that degradosome complexes compete with ribosomes for free mRNAs ^17^. Similarly, protein degradation was completely disregarded except for targeted protease processing of polypeptides in translocation reactions, and even when considered for a subset of the proteome, the computational formulation disregarded the recovery of amino acids as experimentally determined ^47^. In *E. coli*, the membrane protease HflB is essential ^48^, while protein homeostasis is an essential process across bacteria ^49^. Later, protein degradation was considered in ME-models specifically in the context of re-folding proteins after a heat shock mediated by the Lon metalloprotease ^9,12^. The degradation of rRNA, tRNA, and proteins are commonly associated with stress conditions ^6,50^, and therefore, it is highly desirable to reconstruct ME-models containing both types of reactions to better understand and predict the effects of stress and the expense of stress recovery in natural and engineered environments. Here, we proposed an extension to the original ME-model framework ^16^ that greatly expands the number of reactions concerning macromolecule degradation. The approach taken in this coralME update is similar to the protein degradation reactions in the FoldME and StressME models ^9,12^, with degradation reactions added for all proteins in the model with energy expense and recovery of biomass. In contrast to FoldME and StressME models, protein degradation is modeled as the concomitant activities of an ATP-dependent proteasome complex that produces oligopeptides and an ATP-independent protease finalizes the hydrolysis to individual amino acids. This approach was used in *i*MMSYN for the degradation of a single representative protein ^34^. To further expand past the innovations of the FoldME and StressME models, coralME v1.3 can now create degradation reactions for complexes and for modified proteins with weakly-bound cofactors and covalently-bound prosthetic groups. Similar to *i*MMSYN and the mRNA degradation in prior ME-models, we extended the framework to model the degradation of rRNAs, tRNAs, and ncRNAs via the degradosome, producing nucleotide monophosphates. The user can identify and associate other ribonucleases such as the endoribonuclease MazF ^50^ to the degradation process or couple the degradation flux to the availability of ribonucleases, enabling the modeling and study of a wider range of cellular strategies. These features are further enhanced by our update changing coralME’s coupling coefficients to flexible symbolic expressions that are evaluated before simulation, our addition of virtual metabolites that cleanly enforce stoichiometric coupling between reactions, and a greatly expanded system for manipulating reaction stoichiometries as if they were algebraic expressions.

The recalculation of the GAM and NGAM constraints provides a well-established challenge in ME-model reconstruction. These two values represent the amount of energy used for growth and non-growth associated processes, respectively ^38^. Because ME-models explicitly represent translation and other non-metabolic energy-consuming processes, some of the GAM and NGAM energy is already accounted for, and a failure to adjust will force the model to overestimate energy, therefore, underestimating growth rate. For instance, O’Brien et al., 2013 corrected NGAM from 8.39 to 1.00 mmol ATP gDCW^-1^ h^-1^ and GAM from 59.81 to 35.00 mmol ATP gDCW^-1^ ^17,26^, assuming the difference is already represented in the model. To correct GAM and NGAM values, an iterative minimization procedure can be employed where explicit ATP and GTP hydrolysis fluxes are summed up then subtracted from the total GAM value. However, this process is time consuming and requires an arbitrary tolerance error to be set prior to optimization of the difference. This would only create a single, static correction, and would need to be repeated in each condition. Here, we devised an alteration to the ME-formulation that corrects GAM and NGAM values automatically during simulation. The procedure is equivalent to setting a constraint in the LP problem *∑c_i,ATP_v_i_ + ∑c_i,GTP_v_i_ - μ*GAM *-* NGAM *=* 0, where *v_i_* are the fluxes of all the non-metabolic reactions, *μ* is the growth rate, and *c_i,ATP_*and *c_i,GTP_* are the stoichiometric coefficients for ATP and GTP in these reactions, respectively. This forces the model to hydrolyze only the amount of ATP needed to account for the difference between the current, directly modeled energy expenditure and the experimental value. This approach also factors in the dependence of *v_i_* on the growth rate, further demonstrating its value over an independent, iterative minimization problem. By doing this procedure, a toy ME-model shows that its GAM+NGAM value should be 44.46% lower than in the corresponding M-model. In other words, translation and other modeled processes consume 55.54% of the produced energy.

Currently, ME-models cannot simulate exactly zero growth rate due to the mRNA degradation coupling coefficient ^16,17^, but minimizing the ME-model could estimate the explicit NGAM value isolated from the GAM. However, a number of technical challenges must be overcome. For instance, protein synthesis fluxes are linearly proportional to the growth rate ^3,17^, therefore a hypothetical simulation of zero growth rate would have zero protein production and be unable to produce the ATP synthesis machinery needed for non-growth maintenance. Moreover, the synthesis rates for ribosomes, RNAP, mRNAs, tRNAs, and tRNA ligases depend as well on the growth rate, hypothetically collapsing these aspects of the ME-model into an M-model at zero growth rate. While interesting from a theoretical point of view, non-growth maintenance is biologically relevant for bacterial physiology and survival ^6,50^, and so deriving a ME-model suitable for its study would improve understanding and modification of bacterial behavior.

Another identified problem is the lack of systematized tools for the structural analysis of ME-models. Multiple previous works have relied on single gene deletion for the determination of true positives and false negatives to compare to M-models and experimental data ^3,5,7,14,17^. The implementation of such analysis is initiated with the knockout of a translation reaction (i.e., setting the reaction bounds to zero) and the optimization of the model. This single gene deletion analysis was accelerated by replacing a full optimization by a single feasibility check at a predetermined growth rate ^18^, and then by targeting the stoichiometry of transcription reactions rather than the bounds of translation reactions as it was previously done ^14^. However, the algorithm was not systematized into a convenient function either in COBRAme or coralME v1.0, while M-model shows a plethora of such methods, some conveniently incorporated into COBRApy ^22^, supporting the predominant choice of using M-models for research ^51^. Here, we extended coralME to contain analysis methods under the coralme.util.essentiality module, contributing to closing the gap with its M-model counterpart. The ‘perform_gene_knockouts’ is the basic function that performs gene knockouts (single and multiple), while optionally adding sink reactions following user input for the computational analysis of *rescued* phenotypes, while the ‘single_gene_deletion’ function will additionally perform a feasibility check of the growth rate. Moreover, ME-models offer the possibility to determine if enzyme cofactors are essential as well as genes. To do so, all the formation reactions using a certain cofactor can be knocked out before checking if a predetermined growth rate is feasible. This procedure contrasts with M-models as they typically include enzyme cofactors in the biomass reaction and the effect of their elimination is less clearly represented ^52^. Now, coralME can perform this analysis by calling the ‘single_cofactor_essentiality_analysis’ function, returning if an enzyme cofactor is essential or not without modifying the biomass reaction of an ME-model. For a toy ME-model, the methodology determined that metals such as Fe^+2^, K^+1^, Mg^+2^, Mn^+2^, and Zn^+2^ and vitamins such as FAD, NAD, and thiamine are essential for growth in *Mycoplasma mycoides*, while metals such as Ca^+2^, Cd^+2^, Co^+2^, Na^+1^, and Ni^+2^ are individually dispensable. Using the M-model biomass reaction, Ca^+2^ and Na^+1^ would have been identified as essential metabolites. Results of this analysis depend on the correct annotation of reference genes. For instance, the essentiality of Fe^+2^ in the analyzed ME-model is linked to the activities of the peptide deformylase and methionyl aminopeptidase enzymes, essentiality that is controversial in Mycoplasmas ^53^. It is believed the enzymes evolved to not use iron, further supported by the lack of specific transporters for iron. We also demonstrated that this capability of ME-models can be transferred to M-models by extending the GPR to consider metabolites that act as enzymatic cofactors. The coralME software v1.3 extends GPRs to enzymatic cofactors, and the user can determine if they are essential or not by using the ‘single_cofactor_essentiality_analysis’ function. As an example, we updated the *i*ML1515 M-model for *E. coli* ^54^ with cofactor usage information from *i*JL1678b-ME ^16^, resulting in 14 essential cofactors (Supplementary File S6).

In addition to new analysis methods, we extended coralME’s ability to capture differences in gene expression across wild-type and engineered bacteria. Reconstruction of ME-models using coralME v1.0 is restricted to natural bacterial genetic codes #4 (encoding tryptophan in TGA codons) and #11 (encoding selenocysteine in internal TGA stop codons), while COBRAme is only compatible with genetic code #11 (more information https://www.ncbi.nlm.nih.gov/Taxonomy/taxonomyhome.html/index.cgi?chapter=cgencodes). Previously, coralME assumed that a single stop codon is present in the coding DNA sequence, at the end of it. The amino acid stoichiometry is then determined directly from the protein translation using BioPython ^55^. The addition of unnatural, internal stop codons in genomically recoded organisms ^31,32,39–41^ necessitated changes to the architecture of the reconstruction framework itself. Now, coralME v1.3 determines if an internal stop codons have associated tRNAs (TGA with tRNA^(Trp)^ in #4, TGA with tRNA^(Sec)^ in #11), and only if not, translation is terminated. To ensure truncated proteins are flagged appropriately as non-functional, the new *knockouts* reconstruction flag should be used (see Supplementary File S3). For the reconstruction of GRO models, this implies a tripartite linking of codon to amino acid to tRNA from user input. We demonstrated these changes by reconstructing five GRO models. One, rEc.γ.dC.46, resulted in the dependency of a non-canonical amino acid to rescue the essential expression of b0090 (undecaprenyldiphospho-muramoylpentapeptide beta-N-acetylglucosaminyltransferase) and b0893 (L-serine:tRNASer ligase (AMP-forming)) genes, confirming the integration of the atypical genetic code into the model.

## Conclusion

ME-models have revolutionized our capability to understand and predict bacterial physiology under normal growth conditions. However, many aspects of bacterial physiology that are relevant under stress conditions have not been addressed, and computational tools such as COBRAme and coralME have focused on accelerating reconstruction, leaving structural analyses to ad-hoc methods not present in their codebases. Here, we systematized gene and cofactor analyses, expanded the ME-model framework to model RNA and protein degradation, and extended the capability of coralME to model unnatural genomes with protein sequences requiring non-canonical amino acids. In doing so, we expect to strengthen the available software for ME-models, contribute to a broader adoption of ME-model in applied research, and close the gap with M-models and protein allocation modeling frameworks by providing an initial set of structural analysis methods.

## Methods

### Development of new core features

The coralME software version v1.0 ^18^ was updated and modified to accommodate new reaction, process data, and component types (Fig. 1). New reaction types include *BoundaryReaction* for easier identification of exchange, demand, and sink reactions from other *MEReaction* types during queries; *ComplexDegradation* to hold specific methods for the determination of a mass-balanced reaction to fully degrade a macromolecular complex into its constitutive components (i.e., amino acids, modified amino acids, cofactors, ncRNAs, and generics); and *rRNAFormationReaction* that allows the optional degradation of rRNAs if its corresponding coupling coefficient 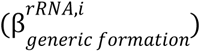 is positive. Similarly, the original *tRNAChargingReaction* was modified to allow the optional degradation of tRNAs 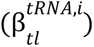. Degradation of mRNA components is now controlled by gene-specific parameters rather than a single, global parameter value 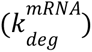 as it was in COBRAme ^16^ and coralME v1.0 ^18^. Regarding process data, a new *rRNAData* class derived from the original *tRNAData*, and both now define the degradation coupling coefficients (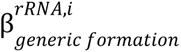 and 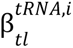, where *i* is the specific gene). A new component type called *Token* allows the creation of pseudo-metabolites and their use as linear constraints such as *v_2_ = kv_1_* where *k* is the coupling coefficient (numeric or symbolic) in the right-hand side of reaction *r_1_*. The addition of the same *Token* in opposite sides of two reactions forces the rate of reaction *r_2_* to be proportional to *r_1_*(Fig. 2). Finally, the user can create new *SummaryVariable* reactions to apply arbitrary constraints from biomass components, and coralME v1.3 will automatically redistribute biomass in *ComplexFormation* reactions during reconstruction (Fig. 2).

The coralME software was also restructured, with new methods added for convenience. The methods related to optimization procedures are now in their own module (*optimization.py*) and attached to a coralME model object upon creation or loading. This reorganization allows better readability, direct access to simulation methods, and maintenance of the code. Three other new modules are included. First, the *extended_classes.py* module declares improved classes such as *ScalableKeyDict* and *MCounter*, dictionary-like objects that unlike dictionaries, can be manipulated as mathematical vectors (sum and scalar multiplication) to update reaction stoichiometries. Next is *diet.py* that includes functions to read and set exchange reaction bounds to simulate different medium compositions. Last, *parameters.py* defines symbolic coupling coefficients, default parameter values, and a physical unit system, as well as declaring methods to modify (parametric) coupling coefficients globally and per reaction. The parameter values are defined in their own object called *DefaultParameters* that creates keys from strings as positive sympy *Symbols*, and retrieves values using string notation, simplifying the modification and reevaluation of parametric coupling coefficients using the *Symbol*’s *eval* method (Supplementary File S1). Therefore, evaluation of parameter sensitivity can be done without reconstructing ME-models with new numerical values or modification of the reaction stoichiometries.

Finally, new utility functions are included to extend and systematize the application of analyses. For instance, the manual curation of M-models can be performed using the output from the *ToExcel* method. The spreadsheet file can be read back as a coralME M-model using the *FromExcel* method, or used directly as input for the MEBuilder method (Supplementary File S1). A coralME M-model differs from a COBRApy M-model as the first adds ‘cofactors’ as a reaction property, setting it as a requirement for reaction functionality upon optimization (Supplementary File S6 and File S7). The new *essentiality.py* module contains methods for the determination of gene essentiality, cofactor essentiality, and metabolite complementation of gene knockouts. And the *agora.py* module contains methods to download and reconstruct ME-models from the (AGORA) resource ^56^. Other changes include the update of reference files for *Escherichia coli* ^16^ and the inclusion of a new reference model based on a ME-model for *Bacillus subtilis* ^7^, and accessed before reconstructions using the *dev_reference* reconstruction flag set as ‘iJL1678b’ (gram negative) or ‘iJT964’ (gram positive), respectively. Examples are described in Supplementary File S1.

The coralME software has been tested under Ubuntu 26.04, gfortran 5, and gurobipy 11, fully compatible with python 3.11 to 3.15, and it can be obtained from GitHub ^57^, Zenodo, or the Python Package Index (pip install coralme --upgrade). The MINOS and quad MINOS LP solvers are now compatible with RHEL (Rocky Linux 8.6) along with debian-based Linux distributions. The source code for coralME v1.3 is available as Supplementary File S8.

### Changes to the ME-model reconstruction pipeline

The reconstruction pipeline from coralME v1.0 ^18^ was refined and performs additional checks. The M-model metabolites are checked for formula and charge completion, and M-model reactions are checked for mass and charge balance. If a metabolite formula and/or charge is missing, the reactions consuming or producing that metabolite are flagged and reported to the user, and if the reaction can be corrected by adding or subtracting protons, the ‘check_and_correct_stoichiometries’ function will proceed to perform and report to the user such modifications. Consequently, the formulas and charges of electron carriers such as ferredoxins are also computed with user-input information regarding component IDs. The new ‘electron_transfers’ flag helps to determine which complex ID requires formula corrections: cytochromes, ferredoxins, glutaredoxins, flavodoxins, and peroxiredoxins change the oxidation state only, while thioredoxins transfer two protons per oxidation state. The ‘add_remaining_complex_formulas’ function can be called after reconstruction to manually and iteratively correct metabolite formulas from user input, and consequently, correct reaction mass and charge balances. Moreover, the new ‘get_me_mass_balance’ reaction method will attempt to determine the mass and charge balance or report a reason why it is not possible. Boundary reactions such as demand, exchange, and sink reactions are mass and charge imbalanced by definition, while other reactions involving generic, constraint, and lipoprotein components that have no formula by definition are skipped. The function reports the output of ‘check_me_mass_balance’ at a hypothetical zero growth rate.

Finally, to help in the reconstruction of ME-models, the processing of homology search results was extended to keep information from reference genes, complexes, and modified complexes. The consequence is the ability to include reference M-reactions without modifying GPRs and to include ME-reactions without changing reference IDs into target IDs. For instance, reconstruction of ME-models that require protoheme for functionality also require a new ME-reaction to produce the modified enzyme hydroxymethylbilane synthase with a dipyrromethane prosthetic group, a reaction readily available from the reconstruction files of the reference model *i*Eco1689-ME ^18^. While this capability is only available for protein components, a new ‘tRNAscan_output’ reconstruction flag helps to determine unannotated tRNA loci in the GenBank input file using the output of tRNAscan-SE ^33^. Currently, the reconstruction will halt if any tRNA is missing, and the user can set the ‘add_missing_tRNAs’ reconstruction flag to assign a random tRNA locus to missing tRNA(s), avoiding an expensive troubleshooting procedure. Similarly, if a 5S, 16S, and/or 23S rRNA locus is missing, coralME will identify upon reconstruction of the ribosome reaction and report the missing component(s). The troubleshooting of the missing rRNA(s) will happen before any other metabolite if it is requested, while previously the missing component was simply not added to the formation of the ribosome (i.e., a sink reaction was not added by the troubleshooter function).

### Development of Genomically Recoded Organism ME-models of *Escherichia coli* (*i*EcoGRO)

To perform the reconstruction of GRO models for *E. coli* strains, the input files to reconstruct *i*Eco1689-ME ^18^ were used without modification except for the GenBank and the transcriptional units (TU) files (see Fig. 1c in ^18^). The TU file was omitted because it encodes TU start and end coordinates tailored to the reference genome ASM584v2. As a consequence, RNA polymerase specificity and Rho termination dependency also encoded in the TU file are missing from GRO models, and we defined the homolog of RpoD as the default sigma factor. We selected six public genomic assemblies, reconstructing ME-models for ASM47403v1 (C321.ΔA) ^31,39,58^, ASM284368v1, an adaptive laboratory evolved strain (ALE-C321.ΔA) ^40^, *rE.coli-57* ^32^, ASM284368v1 (Ochre) ^42^, and both ASM82690v1 (rEc.γ.dC.46) and ASM82692v1 (rEc.β.dC.12) from the same source ^39^ using coralME v1.4.

To reconstruct the E-matrix, coralME relies on genomic annotation for ribosomal proteins, RNA polymerase proteins, tRNA ligases, tRNAs, and rRNAs rather than strict homology. Consequently, for genomic assemblies *rE.coli-57*, ASM82690v1, and ASM82692v1, the NCBI Prokaryotic Genome Annotation Pipeline ^59^ version 2024-07-18.build7555 was used to perform gene annotation under the same namespace as ASM584v2 and the other assemblies. Next, locus tags were renamed to match the M-model *i*JO1366b ^16^ based on strict protein homology to ASM584v2 using the output of Orthofinder 3.0.1b1 ^60,61^. Alternatively, renaming of M-model genes can be performed using COBRApy ‘manipulation.rename_genes’ ^27^ and coralME ‘model.rename_genes’ functions. For identified pseudogenes as recoded genes (i.e., presenting internal TAG codons and lacking of internal TAA codons), the locus tag IDs for lspA, dnaX, tolC, secY, murG, serS, and dnaA, were renamed manually to b0027, b0470, b3035, b3300, b0090, b0893, and b3702 respectively, and added into the ME-model by using the ‘include_pseudo_genes’ option. Similarly, the four tRNA-initiator Met(CAU) genes (two in the case of Ochre) were manually identified from equivalent genomic positions to ASM584v2. To make gene expression of recoded DNA sequences functional, transport reactions of biphenylalanine (bipA) across membranes (BIPAabcpp, BIPAt2pp, BIPAt2rpp, BIPAt4pp, and BIPAtex reactions) were added to the M-model associated with genes homologous to subunits of L-alanine transporters. To reconstruct ΔPrfA strains, the locus tag b1211 was added to the ‘knockouts’ configuration key. Then, to reconstruct GRO models for bipA-dependent strains (ASM82690v1 and ASM82692v1), the sequences for the tRNA and the tRNA ligase genes from the pEVOL-pBpF plasmid ^62,63^ were annotated using tRNAscan-SE v1.4 ^33^ and Prokka ^64^, and appended to the corresponding assembly (i.e., obtaining a two-locus GenBank file for each strain). Finally, the ‘genetic_recoding’ configuration flag assigns the TAG (stop) codon to the metabolite ‘bipa L_c’ and the ‘trna-bipA’ locus tag, while the ‘amino_acid_synthetase’ dictionary maps the metabolite ‘bipa L_c’ to the bipA_tRNA_synth_copy1-MONOMER complex ID. Simulation was performed using the default medium (M9-glucose) defined in *i*JO1366b ^16^ supplemented with biotin and bipA, and the quad MINOS solver ^35,36^ with default options (growth rate error 1e-6, quadratic-precision float-point arithmetic). The GRO models are available from https://github.com/ZenglerLab/iEco-GRO and they can be reconstructed using Supplementary File S2 (generation of input files), Supplementary File S3 (building of ME-models), and Supplementary File S4 (optimization of ME-models). Manual curation data was obtained from the reference ME-model and copied manually after generation of input files.

### Protein and RNA degradation reactions and development of a toy ME-model

Protein degradation reactions can be added to an ME-model during reconstruction using the ‘add_prot_deg_reactions’ reconstruction flag (default value is False), or after reconstruction using the ‘create_complex_degradation’ method (see Supplementary File S1). The method will determine a stoichiometrically complete and mass-balanced reaction that will recover protein biomass and other biomass constraints if they participate in the corresponding formation reaction (Fig. 1). The degradation of prosthetic groups produces modified amino acids, such as biocytin from biotinylated proteins, except for the case of the holo-acyl carrier protein for which the ACP phosphodiesterase can be already incorporated in M-models (e.g., ACPpds). Current supported modified amino acids are N(6)-lipoyl-L-lysine, biocytin, dipyrrolylmethanemethyl-L-cysteine, and N-acetyl-L-alanine that require sink or metabolic reactions to avoid model infeasibility. In the case of RNA degradation reactions (Fig. 1), the update method of *TranscriptionReaction*, *rRNAFormationReaction*, and *tRNAChargingReaction* reactions will include a stoichiometrically complete and mass-balanced subreaction for degradation of transcription units, rRNAs, and tRNAs, respectively. Similar to protein degradation reactions, degradation of rRNAs and tRNAs recover the respective biomass constraint.

A toy ME-model was developed to assess mass-balance and correctness of the newly created degradation reactions. The model derives from the *Mycoplasma mycoides i*MMSYN model ^34^ and the *M. mycoides* ME-model ^18^ with additional manual curation. The respective M-model was curated to complete metabolite formulas and charges, add sink reactions for iron(II), manganese(II), and zinc(II) for functionality of the peptide deformylase and methionyl aminopeptidase and other complexes, while the ME-model was curated to add reactions such as the glutamyl-tRNA(Gln) amidotransferase and new protein complex stoichiometries such as the pyruvate dehydrogenase. Protein degradation is assigned to JCVISYN3A_0394 (Endopeptidase La) homologous to the *Escherichia coli* Lon metalloprotease and enzymatically equivalent to ClpXP ^34^, and peptidases JCVISYN3A_0133, JCVISYN3A_0305, JCVISYN3A_0444, and JCVISYN3A_0479 (peptidase-1, 2, 3, and 4, respectively). Biomass components and stoichiometric coefficients (*i*MMSYN’s *BIOMASS* reaction) were transferred to four *SummaryVariable* reactions (cofactors, lipids, biomass, and DNA), with the exception of amino acids, ribonucleotides, ATP, water, protons, and phosphate. The growth-associated ATP maintenance (GAM) was set to 25.0 mmol of ATP gDCW^-1^, while NGAM (non-GAM) was set to 1.0 mmol of ATP gDCW^-1^ h^-1^. Input files to reconstruct the ME-model can be found https://github.com/ZenglerLab/supp-coralme-v1.3. Simulations were performed using the default medium (SP4) defined in *i*MMSYN ^34^ and the quad MINOS solver with default options (growth rate error 1e-6, quadratic-precision float-point arithmetic).

### Correction of NGAM and GAM values

The reconstruction of M-models requires the parametrization of both NGAM and GAM values to represent the energy consumed under non-growth cellular and non-metabolic processes, respectively ^26^. For the reconstruction of ME-models, the coralME software uses NGAM and GAM derived from the M-model without correction, leading to an overestimation of the energy. Previously, corrections have been applied [NGAM was reduced from 8.39 to 1.00 mmol ATP gDCW^-1^ h^-1^, and GAM from 59.81 to 35.00 mmol ATP gDCW^-1^ ^16^], but no reproducible method was presented. A modified ME-model was derived to determine the explicit and implicit amounts of energy from simulation in the following way:

1. A *Token* metabolite is created (e.g., ‘gam_token’) and added to the ME-model.
2. The token is added in the right-hand side of translation reactions and any other reaction in the E-matrix that consumes ATP and/or GTP, associated with a stoichiometric coefficient from the sum of both coefficients for ATP and GTP.
3. The GAM reaction bounds are set to (0, 1000) and the NGAM reaction bounds to (0, 0).
4. A demand reaction is added to the ME-model with bounds equal to *µGAM+NGAM* that consumes the token and keeps the pseudo-stationary state.

### Gene and cofactor essentiality analyses of a toy ME-model

To perform gene essentiality analysis, each transcription reaction where a gene participates was modified to remove the gene product and updated to recalculate reaction stoichiometry, rendering subsequent reactions (i.e., translation) that use the gene product as a substrate unable to carry flux. Simulations were performed using the default medium in *i*Mmy281-ME fixing the growth rate to 0.01 h^-1^ and the quad MINOS solver with default options (quadratic-precision float-point arithmetic). The simulation of the ME-model was carried out using the default bounds for glucose uptake (7.4 mmol gDCW^-1^ h^-1^), glycerol (0.193 mmol gDCW^-1^ h^-1^), and a maximum acetate secretion rate (6.9 mmol gDCW^-1^ h^-1^).

To perform cofactor essentiality analysis, all upper bounds of formation reactions associated with one enzymatic cofactor were set to 0.0 mmol gDCW^-1^ h^-1^, rendering subsequent reactions (i.e., post-translation and metabolic reactions) unable to carry flux if the cofactor uptake or the *de novo* biosynthesis is essential. Manipulation of the formation reactions avoids false positives due to components in the biomass reaction (Fig. 2). Simulations were performed using the default medium in *i*Mmy281-ME fixing the growth rate to 0.01 h^-1^ and the quad MINOS solver with default options (quadratic-precision float-point arithmetic). Analysis of gene and cofactor essentiality can be reproduced by using Supplementary File S5.

## Resource availability

### Lead contact

Further information and requests for resources and reagents should be directed to and will be fulfilled by the lead contact, Karsten Zengler.

### Materials availability

This study did not generate new materials.

### Data and code availability

No experimental data were generated in this work. The reconstructed ME-models and all related files are available at GitHub repositories https://github.com/ZenglerLab/supp-coralme-v1.3 and https://github.com/ZenglerLab/iEco-GRO and Zenodo repositories.

The coralME software is open-source and freely available under an MIT license from GitHub^94^/Zenodo and the Python Package Index.

Any additional information required to reanalyze the data reported in this paper is available from the lead contact upon request.

### Funding

This material is based upon work supported by the U.S. Department of Energy, Office of Science, Biological and Environmental Research under the ‘Accelerate Innovations in Emerging Technologies’ program, as well as by the Genomic Science Program under Secure Biosystems Design Science Focus Area (SFA) contract number DE-AC36-08GO28308 and the National Science Foundation under award numbers EFMA-2223669 and DMS-2325172.

### Author contributions

Conceptualization, R.S.-P., J.D.T.-B., D.T.-C., M.J.N., and K.Z.; methodology, R.S.-P., J.D.T.-B., and K.Z.; investigation, R.S.-P., J.D.T.-B., and K.Z.; visualization, R.S.-P.; supervision, K.Z.; writing – original draft, R.S.-P., J.D.T.-B., D.T.-C., M.J.N., and K.Z.; and writing – review and editing, R.S.-P. and K.Z.

