## Supplementary File S1 for "Expanded ME-model analysis capabilities enable genetic recoding, proteostasis, RNA decay, and cofactor essentiality in bacterial models"

November 4, 2026

[1]:
```
# # Install dependencies
# !{sys.executable} -m pip install itables
```

[2]:
```
import itables
itables.init_notebook_mode(all_interactive=True)
```

```
<IPython.core.display.HTML object>
```

```
<IPython.core.display.HTML object>
```

[3]:
```
import os
os.environ["TQDM_DISABLE"] = "1"
from tqdm import tqdm
```

[4]:
```
# # Install dependencies
# !{sys.executable} -m pip install -r Supplementary-File-S8/requirements.txt
```

[5]:
```
import sys
sys.path.insert(0, './Supplementary-File-S8')
import coralme
coralme.__file__.split('Supplementary-File-S8')[1]
```

[5]:
```
'/coralme/__init__.py'
```

### 0.0.1 Reconstruction of a toy ME-model: iMmy281-ME

[6]:
```
cofactors_in_biomass = {
    "ca2_c": -0.004658,
    "cl_c": -0.055853,
    "coa_c": -0.000158,
    "fad_c": -0.00021,
    "k_c": -0.840160,
    "mg2_c": -0.007763,
    "methfglu3_c": -0.00021,
    "na1_c": -0.057153,
    "nadp_c": -0.000105,
    "pydx5p_c": -0.00021,
    "sprm_c": -0.006344,
    "thmpp_c": -0.000210
```

```
    }

import coralme
builder = coralme.builder.main.MEBuilder(
    config_files=['iMmy281-ME-inputs/config-input.json', 'iMmy281-ME-inputs/
↪config-organism.json'],
    out_directory='iMmy281-ME-cofactors-as-biomass-constituents',
    log_directory='iMmy281-ME-cofactors-as-biomass-constituents')

builder.configuration['flux_of_biomass_constituents'].
↪update(cofactors_in_biomass)

builder.generate_files()
builder.build_me_model()

builder.me_model.metabolites.fe2_c.add_boundary(type = 'sink')
builder.me_model.metabolites.mn2_c.add_boundary(type = 'sink')
builder.me_model.metabolites.zn2_c.add_boundary(type = 'sink')

builder.troubleshoot()
```

```
Initiating file processing…
~ Processing files for mycoplasmaME…
Reading mycoplasmaME done.
~ Processing files for iJL1678b…
Reading iJL1678b done.
~ Running BLAST with 15 threads…
BLAST done.
Writing the Organism-Specific Matrix to iMmy281-ME-inputs/automated-org-with-
refs.xlsx…
Organism-Specific Matrix saved to iMmy281-ME-inputs/automated-org-with-refs.xlsx
file.
File processing done.
Initiating ME-model reconstruction…
ME-model was saved in the iMmy281-ME-cofactors-as-biomass-constituents directory
as MEModel-step1-mycoplasmaME.pkl
ME-model was saved in the iMmy281-ME-cofactors-as-biomass-constituents directory
as MEModel-step2-mycoplasmaME.pkl
ME-model reconstruction is done.
Number of metabolites in the ME-model is 1228 (+297.41%, from 309)
Number of reactions in the ME-model is 1884 (+486.92%, from 321)
Number of genes in the ME-model is 281 (+88.59%, from 149)
Number of missing genes from reconstruction with homology, but no function is 9.
Check the curation notes for more details.
The MINOS and quad MINOS solvers are a courtesy of Prof Michael A. Saunders.
Please cite Ma, D., Yang, L., Fleming, R. et al. Reliable and efficient solution
of genome-scale models of Metabolism and macromolecular Expression. Sci Rep 7,
```

```
40863 (2017). https://doi.org/10.1038/srep40863

~ Troubleshooting started…
  Checking if the coralME model can simulate growth without gapfilling
reactions…
  Original coralME model is feasible with a tested growth rate of 0.001000 1/h
~ Final step. Fully optimizing with precision 1e-6 and save solution into the
coralME model…
  Gapfilled coralME model is feasible with growth rate 0.310233 (M-model:
0.367676).
coralME model was saved in the iMmy281-ME-cofactors-as-biomass-constituents
directory as MEModel-step3-mycoplasmaME-TS.pkl
```

#### 0.0.2 Reconstruction of a toy ME-model: iMmy281-ME (Reference B. subtilis)

[7]:
```
import coralme
builder = coralme.builder.main.MEBuilder(
    reference = 'iJT964',
    config_files=['iMmy281-ME-inputs/config-input.json', 'iMmy281-ME-inputs/
 ↪config-organism.json'],
  ␣
 ↪out_directory='iMmy281-ME-cofactors-as-biomass-constituents-Bsubtilis-as-reference',
  ␣
 ↪log_directory='iMmy281-ME-cofactors-as-biomass-constituents-Bsubtilis-as-reference',**{'df_
 ↪: ''})

builder.configuration['flux_of_biomass_constituents'].
 ↪update(cofactors_in_biomass)

builder.generate_files()
builder.build_me_model()

# builder.me_model.metabolites.fe2_c.add_boundary(type = 'sink')
builder.me_model.metabolites.mn2_c.add_boundary(type = 'sink')
builder.me_model.metabolites.zn2_c.add_boundary(type = 'sink')

builder.troubleshoot()
```

```
Initiating file processing…
~ Processing files for mycoplasmaME…
Reading mycoplasmaME done.
~ Processing files for iJT964…
Reading iJT964 done.
~ Running BLAST with 15 threads…
BLAST done.
Writing the Organism-Specific Matrix to iMmy281-ME-inputs/automated-org-with-
refs.xlsx…
Organism-Specific Matrix saved to iMmy281-ME-inputs/automated-org-with-refs.xlsx
```

```
file.
File processing done.
Initiating ME-model reconstruction…
ME-model was saved in the iMmy281-ME-cofactors-as-biomass-constituents-
Bsubtilis-as-reference directory as MEModel-step1-mycoplasmaME.pkl
ME-model was saved in the iMmy281-ME-cofactors-as-biomass-constituents-
Bsubtilis-as-reference directory as MEModel-step2-mycoplasmaME.pkl
ME-model reconstruction is done.
Number of metabolites in the ME-model is 1213 (+292.56%, from 309)
Number of reactions in the ME-model is 1865 (+481.00%, from 321)
Number of genes in the ME-model is 276 (+85.23%, from 149)
Number of missing genes from reconstruction with homology, but no function is 4.
Check the curation notes for more details.
The MINOS and quad MINOS solvers are a courtesy of Prof Michael A. Saunders.
Please cite Ma, D., Yang, L., Fleming, R. et al. Reliable and efficient solution
of genome-scale models of Metabolism and macromolecular Expression. Sci Rep 7,
40863 (2017). https://doi.org/10.1038/srep40863

~ Troubleshooting started…
  Checking if the coralME model can simulate growth without gapfilling
reactions…
  Original coralME model is feasible with a tested growth rate of 0.001000 1/h
~ Final step. Fully optimizing with precision 1e-6 and save solution into the
coralME model…
  Gapfilled coralME model is feasible with growth rate 0.309748 (M-model:
0.367676).
coralME model was saved in the iMmy281-ME-cofactors-as-biomass-constituents-
Bsubtilis-as-reference directory as MEModel-step3-mycoplasmaME-TS.pkl
```

### 0.0.3 Reconstruction of a toy ME-model: iMmy281-ME (cofactors as constraints)

[8]:
```python
import coralme
builder = coralme.builder.main.MEBuilder(
    config_files=['iMmy281-ME-inputs/config-input.json', 'iMmy281-ME-inputs/
 ↪config-organism.json'],
    out_directory='iMmy281-ME-cofactors-as-biomass-constraints',
    log_directory='iMmy281-ME-cofactors-as-biomass-constraints')

builder.configuration['flux_of_biomass_constraints'] = cofactors_in_biomass
builder.configuration['add_prot_deg_reactions'] = False

builder.generate_files()
builder.build_me_model()

builder.me_model.metabolites.fe2_c.add_boundary(type = 'sink')
builder.me_model.metabolites.mn2_c.add_boundary(type = 'sink')
builder.me_model.metabolites.zn2_c.add_boundary(type = 'sink')
```

```
builder.troubleshoot()
```

```
Initiating file processing…
~ Processing files for mycoplasmaME…
Reading mycoplasmaME done.
~ Processing files for iJL1678b…
Reading iJL1678b done.
~ Running BLAST with 15 threads…
BLAST done.
Writing the Organism-Specific Matrix to iMmy281-ME-inputs/automated-org-with-
refs.xlsx…
Organism-Specific Matrix saved to iMmy281-ME-inputs/automated-org-with-refs.xlsx
file.
File processing done.
Initiating ME-model reconstruction…
ME-model was saved in the iMmy281-ME-cofactors-as-biomass-constraints directory
as MEModel-step1-mycoplasmaME.pkl
ME-model was saved in the iMmy281-ME-cofactors-as-biomass-constraints directory
as MEModel-step2-mycoplasmaME.pkl
ME-model reconstruction is done.
Number of metabolites in the ME-model is 1240 (+301.29%, from 309)
Number of reactions in the ME-model is 1908 (+494.39%, from 321)
Number of genes in the ME-model is 281 (+88.59%, from 149)
Number of missing genes from reconstruction with homology, but no function is 9.
Check the curation notes for more details.
The MINOS and quad MINOS solvers are a courtesy of Prof Michael A. Saunders.
Please cite Ma, D., Yang, L., Fleming, R. et al. Reliable and efficient solution
of genome-scale models of Metabolism and macromolecular Expression. Sci Rep 7,
40863 (2017). https://doi.org/10.1038/srep40863

~ Troubleshooting started…
  Checking if the coralME model can simulate growth without gapfilling
reactions…
  Original coralME model is feasible with a tested growth rate of 0.001000 1/h
~ Final step. Fully optimizing with precision 1e-6 and save solution into the
coralME model…
  Gapfilled coralME model is feasible with growth rate 0.114033 (M-model:
0.367676).
coralME model was saved in the iMmy281-ME-cofactors-as-biomass-constraints
directory as MEModel-step3-mycoplasmaME-TS.pkl
```

### 0.0.4 Reconstruct ME-model adding protein degradation reactions

[9]:
```
import coralme
builder = coralme.builder.main.MEBuilder(
```

```
    config_files=['iMmy281-ME-inputs/config-input.json', 'iMmy281-ME-inputs/
↪config-organism.json'],
    out_directory='iMmy281-ME-cofactors-as-biomass-constraints',
    log_directory='iMmy281-ME-cofactors-as-biomass-constraints')

builder.configuration['flux_of_biomass_constituents'].
↪update(cofactors_in_biomass)
builder.configuration['add_prot_deg_reactions'] = True

builder.generate_files()
builder.build_me_model()

builder.me_model.metabolites.fe2_c.add_boundary(type = 'sink')
builder.me_model.metabolites.mn2_c.add_boundary(type = 'sink')
builder.me_model.metabolites.zn2_c.add_boundary(type = 'sink')

builder.me_model.metabolites.acala__L_c.add_boundary(type = 'sink')

builder.troubleshoot()
```

```
Initiating file processing…
~ Processing files for mycoplasmaME…
Reading mycoplasmaME done.
~ Processing files for iJL1678b…
Reading iJL1678b done.
~ Running BLAST with 15 threads…
BLAST done.
Writing the Organism-Specific Matrix to iMmy281-ME-inputs/automated-org-with-
refs.xlsx…
Organism-Specific Matrix saved to iMmy281-ME-inputs/automated-org-with-refs.xlsx
file.
File processing done.
Initiating ME-model reconstruction…
ME-model was saved in the iMmy281-ME-cofactors-as-biomass-constraints directory
as MEModel-step1-mycoplasmaME.pkl
ME-model was saved in the iMmy281-ME-cofactors-as-biomass-constraints directory
as MEModel-step2-mycoplasmaME.pkl
ME-model reconstruction is done.
Number of metabolites in the ME-model is 1229 (+297.73%, from 309)
Number of reactions in the ME-model is 2147 (+568.85%, from 321)
Number of genes in the ME-model is 281 (+88.59%, from 149)
Number of missing genes from reconstruction with homology, but no function is 9.
Check the curation notes for more details.
The MINOS and quad MINOS solvers are a courtesy of Prof Michael A. Saunders.
Please cite Ma, D., Yang, L., Fleming, R. et al. Reliable and efficient solution
of genome-scale models of Metabolism and macromolecular Expression. Sci Rep 7,
40863 (2017). https://doi.org/10.1038/srep40863
```

```
~ Troubleshooting started…
  Checking if the coralME model can simulate growth without gapfilling
reactions…
  Original coralME model is feasible with a tested growth rate of 0.001000 1/h
~ Final step. Fully optimizing with precision 1e-6 and save solution into the
coralME model…
  Gapfilled coralME model is feasible with growth rate 0.310233 (M-model:
0.367676).
coralME model was saved in the iMmy281-ME-cofactors-as-biomass-constraints
directory as MEModel-step3-mycoplasmaME-TS.pkl
```

[10]:
```
sol = builder.me_model.gem.optimize()
builder.me_model.growth_rate / sol.objective_value # sol.objective_value␣
↪(COBRApy) has no unit associated
```

[10]: $0.8437681057380937 \frac{1}{\text{hour}}$

[11]:
```
# WARNING; two sets of complex data produce the same complex from different␣
↪precursors
len(builder.me_model.complex_data)
```

[11]: 266

[12]:
```
for cplx in builder.me_model.complex_data:
    if builder.me_model.reactions.has_id('degradation_'+cplx.id) or␣
↪'_lipoyl_scavenging' in cplx.id:
        continue
    else:
        print(cplx.id)
```

```
JCVISYN3A_0621-MONOMER_mod_pan4p(1)
```

[13]:
```
builder.me_model.remove_troubleshooted_reactions()
for rxn in tqdm(builder.me_model.query(types = [coralme.core.reaction.
↪ComplexDegradation ])):
    rxn.lower_bound = 0.00035/244
    if not builder.me_model.feasibility():
        print(rxn.id)
    rxn.lower_bound = 0.
builder.me_model.growth_rate
```

[13]: $0.01 \frac{1}{\text{hour}}$

### 0.0.5 Save and use M-model from a spreadsheet file as input file for ME-model reconstruction

[14]:

```
coralme.util.model2excel.ToExcel(builder.me_model.gem, 'iMmy149.xlsx')

import coralme
builder = coralme.builder.main.MEBuilder(
    m_model_path='iMmy149.xlsx',
    genbank_path='iMmy281-ME-inputs/genome.gb',
    df_gene_cplxs_mods_rxns='iMmy281-ME-inputs/automated-org-with-refs.xlsx',
    df_matrix_stoichiometry='iMmy281-ME-inputs/reaction_matrix.txt',
    df_matrix_subrxn_stoich='iMmy281-ME-inputs/subreaction_matrix.txt',
    df_metadata_metabolites='iMmy281-ME-inputs/me_metabolites.txt',
    config_files=['iMmy281-ME-inputs/config-organism.json'],
    out_directory='iMmy281-ME-cofactors-as-biomass-constraints',
    log_directory='iMmy281-ME-cofactors-as-biomass-constraints')

builder.configuration['flux_of_biomass_constraints'] = cofactors_in_biomass
builder.configuration['add_prot_deg_reactions'] = True

builder.generate_files()
builder.build_me_model()

builder.me_model.metabolites.fe2_c.add_boundary(type = 'sink')
builder.me_model.metabolites.mn2_c.add_boundary(type = 'sink')
builder.me_model.metabolites.zn2_c.add_boundary(type = 'sink')

builder.troubleshoot()
```

```
Initiating file processing…
~ Processing files for coralME…
INFO: genes are added from the 'reactions' spreadsheet, and gene annotations
from the 'genes' spreadsheet.
Reading coralME done.
~ Processing files for iJL1678b…
Reading iJL1678b done.
~ Running BLAST with 15 threads…
BLAST done.
Writing the Organism-Specific Matrix to iMmy281-ME-inputs/automated-org-with-
refs.xlsx…
Organism-Specific Matrix saved to iMmy281-ME-inputs/automated-org-with-refs.xlsx
file.
File processing done.
Initiating ME-model reconstruction…
ME-model was saved in the iMmy281-ME-cofactors-as-biomass-constraints directory
as MEModel-step1-coralME.pkl
ME-model was saved in the iMmy281-ME-cofactors-as-biomass-constraints directory
as MEModel-step2-coralME.pkl
```

```
ME-model reconstruction is done.
Number of metabolites in the ME-model is 1241 (+301.62%, from 309)
Number of reactions in the ME-model is 2171 (+576.32%, from 321)
Number of genes in the ME-model is 281 (+88.59%, from 149)
Number of missing genes from reconstruction with homology, but no function is 9.
Check the curation notes for more details.
The MINOS and quad MINOS solvers are a courtesy of Prof Michael A. Saunders.
Please cite Ma, D., Yang, L., Fleming, R. et al. Reliable and efficient solution
of genome-scale models of Metabolism and macromolecular Expression. Sci Rep 7,
40863 (2017). https://doi.org/10.1038/srep40863

~ Troubleshooting started…
  Checking if the coralME model can simulate growth without gapfilling
reactions…
  Original coralME model is feasible with a tested growth rate of 0.001000 1/h
~ Final step. Fully optimizing with precision 1e-6 and save solution into the
coralME model…
  Gapfilled coralME model is feasible with growth rate 0.114033 (M-model:
0.367676).
coralME model was saved in the iMmy281-ME-cofactors-as-biomass-constraints
directory as MEModel-step3-coralME-TS.pkl
```

[15]:
```
m_model = coralme.util.excel2model.FromExcel('iMmy149.xlsx')

import coralme
builder = coralme.builder.main.MEBuilder(
    m_model_path=m_model,
    genbank_path='iMmy281-ME-inputs/genome.gb',
    df_gene_cplxs_mods_rxns='iMmy281-ME-inputs/automated-org-with-refs.xlsx',
    df_matrix_stoichiometry='iMmy281-ME-inputs/reaction_matrix.txt',
    df_matrix_subrxn_stoich='iMmy281-ME-inputs/subreaction_matrix.txt',
    df_metadata_metabolites='iMmy281-ME-inputs/me_metabolites.txt',
    config_files=['iMmy281-ME-inputs/config-organism.json'],
    out_directory='iMmy281-ME-cofactors-as-biomass-constraints',
    log_directory='iMmy281-ME-cofactors-as-biomass-constraints')

builder.configuration['flux_of_biomass_constraints'] = cofactors_in_biomass
builder.configuration['add_prot_deg_reactions'] = True

builder.generate_files()
builder.build_me_model()

builder.me_model.metabolites.fe2_c.add_boundary(type = 'sink')
builder.me_model.metabolites.mn2_c.add_boundary(type = 'sink')
builder.me_model.metabolites.zn2_c.add_boundary(type = 'sink')

builder.troubleshoot()
```

```
INFO: genes are added from the 'reactions' spreadsheet, and gene annotations
from the 'genes' spreadsheet.
Initiating file processing…
~ Processing files for coralME…
Reading coralME done.
~ Processing files for iJL1678b…
Reading iJL1678b done.
~ Running BLAST with 15 threads…
BLAST done.
Writing the Organism-Specific Matrix to iMmy281-ME-inputs/automated-org-with-
refs.xlsx…
Organism-Specific Matrix saved to iMmy281-ME-inputs/automated-org-with-refs.xlsx
file.
File processing done.
Initiating ME-model reconstruction…
ME-model was saved in the iMmy281-ME-cofactors-as-biomass-constraints directory
as MEModel-step1-coralME.pkl
ME-model was saved in the iMmy281-ME-cofactors-as-biomass-constraints directory
as MEModel-step2-coralME.pkl
ME-model reconstruction is done.
Number of metabolites in the ME-model is 1241 (+301.62%, from 309)
Number of reactions in the ME-model is 2171 (+576.32%, from 321)
Number of genes in the ME-model is 281 (+88.59%, from 149)
Number of missing genes from reconstruction with homology, but no function is 9.
Check the curation notes for more details.
The MINOS and quad MINOS solvers are a courtesy of Prof Michael A. Saunders.
Please cite Ma, D., Yang, L., Fleming, R. et al. Reliable and efficient solution
of genome-scale models of Metabolism and macromolecular Expression. Sci Rep 7,
40863 (2017). https://doi.org/10.1038/srep40863

~ Troubleshooting started…
  Checking if the coralME model can simulate growth without gapfilling
reactions…
  Original coralME model is feasible with a tested growth rate of 0.001000 1/h
~ Final step. Fully optimizing with precision 1e-6 and save solution into the
coralME model…
  Gapfilled coralME model is feasible with growth rate 0.114033 (M-model:
0.367676).
coralME model was saved in the iMmy281-ME-cofactors-as-biomass-constraints
directory as MEModel-step3-coralME-TS.pkl
```

### 0.0.6 New RNA degradation reactions

[16]:
```
import coralme
model = coralme.io.pickle.load_pickle_me_model('iMmy281-ME.pkl')
# model.to_json('iMmy281-ME.json')
# model = coralme.io.json.load_json_me_model('iMmy281-ME.json')
```

[17]:
```python
# new rRNAFormationReaction
model.reactions.RNA_JCVISYN3A_0067_to_generic_5s_rRNAs.reaction
```

[17]: '1.0 RNA_JCVISYN3A_0067 --> 1.0 generic_5s_rRNAs'

[18]:
```python
# new rRNAFormationReaction
model.reactions.RNA_JCVISYN3A_0068_to_generic_23s_rRNAs.reaction
```

[18]: '1.0 RNA_JCVISYN3A_0068 --> 1.0 generic_23s_rRNAs'

[19]:
```python
# new rRNAFormationReaction
model.reactions.RNA_JCVISYN3A_0069_to_generic_16s_rRNAs.symbolic_stoichiometry
```

[19]:
```
{'RNA_JCVISYN3A_0069': -beta_{generic\ formation}^{rRNA_JCVISYN3A_0069} - 1.0,
 'RNA_degradosome': <Quantity(-beta_{generic\
formation}^{rRNA_JCVISYN3A_0069}*mu/(3600*k^default_cat), 'dimensionless')>,
 'amp_c': 434*beta_{generic\ formation}^{rRNA_JCVISYN3A_0069},
 'cmp_c': 304*beta_{generic\ formation}^{rRNA_JCVISYN3A_0069},
 'generic_16s_rRNAs': 1.0,
 'gmp_c': 434*beta_{generic\ formation}^{rRNA_JCVISYN3A_0069},
 'h2o_c': -1534*beta_{generic\ formation}^{rRNA_JCVISYN3A_0069},
 'rRNA_biomass': -495.060664115*beta_{generic\ formation}^{rRNA_JCVISYN3A_0069},
 'ump_c': 363*beta_{generic\ formation}^{rRNA_JCVISYN3A_0069}}
```

[20]:
```python
# extended tRNAChargingReaction
model.reactions.charging_tRNA_JCVISYN3A_0070_CUA.symbolic_stoichiometry
```

[20]:
```
{'JCVISYN3A_0634-MONOMER': <Quantity(-mu*(f_tRNA*m_aa*(r_0 + mu/k_t)/m_tRNA +
1)/(3600*k^default_cat), 'dimensionless')>,
 'RNA_JCVISYN3A_0070': <Quantity(-beta_{tl}^{tRNA_JCVISYN3A_0070} -
f_tRNA*m_aa*(r_0 + mu/k_t)/m_tRNA, 'dimensionless')>,
 'RNA_degradosome':
<Quantity(-beta_{tl}^{tRNA_JCVISYN3A_0070}*mu/(3600*k^default_cat),
'dimensionless')>,
 'amp_c': 20*beta_{tl}^{tRNA_JCVISYN3A_0070},
 'cmp_c': 21*beta_{tl}^{tRNA_JCVISYN3A_0070},
 'generic_tRNA_CUA_leu__L_c': 1.0,
 'gmp_c': 23*beta_{tl}^{tRNA_JCVISYN3A_0070},
 'h2o_c': -83*beta_{tl}^{tRNA_JCVISYN3A_0070},
 'leu__L_c': <Quantity(-f_tRNA*m_aa*(r_0 + mu/k_t)/m_tRNA, 'dimensionless')>,
 'tRNA_biomass': -26.971300784*beta_{tl}^{tRNA_JCVISYN3A_0070},
 'ump_c': 20*beta_{tl}^{tRNA_JCVISYN3A_0070}}
```

[21]:
```python
# extended TranscriptionReaction
model.reactions.transcription_TU_JCVISYN3A_0004.symbolic_stoichiometry
```

[21]:
```
{'GreA_mono': <Quantity(-mu/(3600*k^default_cat), 'dimensionless')>,
 'NusA_mono': <Quantity(-mu/(3600*k^default_cat), 'dimensionless')>,
```

```
 'NusG_mono': <Quantity(-mu/(3600*k^default_cat), 'dimensionless')>,
 'RNAP_JCVISYN3A0407MONOMER': <Quantity(-267.0*f_rRNA*m_aa*(r_0 + mu/k_t)/m_rr,
'dimensionless')>,
 'RNA_JCVISYN3A_0004': 1.0,
 'amp_c': 347*beta_{tx}^{TU_JCVISYN3A_0004},
 'atp_c': -347*beta_{tx}^{TU_JCVISYN3A_0004} - 347,
 'cmp_c': 67*beta_{tx}^{TU_JCVISYN3A_0004},
 'ctp_c': -67*beta_{tx}^{TU_JCVISYN3A_0004} - 67,
 'gmp_c': 91*beta_{tx}^{TU_JCVISYN3A_0004},
 'gtp_c': -91*beta_{tx}^{TU_JCVISYN3A_0004} - 91,
 'h2o_c': -800*beta_{tx}^{TU_JCVISYN3A_0004},
 'mRNA_biomass': 255.91306706100002,
 'ppi_c': 801,
 'ump_c': 296*beta_{tx}^{TU_JCVISYN3A_0004},
 'utp_c': -296*beta_{tx}^{TU_JCVISYN3A_0004} - 296}
```

[22]:
```
# Stoichiometric coefficient for mRNA in translation reactions is␣
↪beta^{mRNA,i}_{tl} + alpha^{mRNA,i}_{tl}
# The coefficient is negative indicating it is a substrate of the reaction
model.reactions.translation_JCVISYN3A_0004.
↪symbolic_stoichiometry['RNA_JCVISYN3A_0004'].magnitude
```

[22]:

$$-\frac{0.3333333333333333 f_{mRNA} k_{deg}^{mRNA_{J}CVISYN3A_{0}004} m_{aa}\left(r_0+\frac{\mu}{k_t}\right)}{m_{nt}\mu}-\frac{0.3333333333333333 f_{mRNA} m_{aa}\left(r_0+\frac{\mu}{k_t}\right)}{m_{nt}}$$

[23]:
```
# and modified ribosome formation reaction
# notice the prosthetic_group_biomass coefficient is he contribution of 1 GTP␣
↪in gram per millimole (see next cell)
model.reactions.formation_ribosome.reaction
```

[23]:
```
'[4.27350427350427e-6*mu] Era_dim + 1.0 JCVISYN3A_0025-MONOMER + 1.0
JCVISYN3A_0027-MONOMER + 1.0 JCVISYN3A_0082-MONOMER + 1.0 JCVISYN3A_0137-MONOMER
+ 1.0 JCVISYN3A_0148-MONOMER + 1.0 JCVISYN3A_0149-MONOMER + 1.0
JCVISYN3A_0198-MONOMER + 1.0 JCVISYN3A_0199-MONOMER + 1.0 JCVISYN3A_0238-MONOMER
+ 1.0 JCVISYN3A_0294-MONOMER + 1.0 JCVISYN3A_0362-MONOMER + 1.0
JCVISYN3A_0365-MONOMER + 1.0 JCVISYN3A_0422-MONOMER + 1.0 JCVISYN3A_0482-MONOMER
+ 1.0 JCVISYN3A_0499-MONOMER + 1.0 JCVISYN3A_0500-MONOMER + 1.0
JCVISYN3A_0501-MONOMER + 1.0 JCVISYN3A_0526-MONOMER + 1.0 JCVISYN3A_0540-MONOMER
+ 1.0 JCVISYN3A_0637-MONOMER + 1.0 JCVISYN3A_0638-MONOMER + 1.0
JCVISYN3A_0644-MONOMER + 1.0 JCVISYN3A_0646-MONOMER + 1.0 JCVISYN3A_0647-MONOMER
+ 1.0 JCVISYN3A_0648-MONOMER + 1.0 JCVISYN3A_0653-MONOMER + 1.0
JCVISYN3A_0654-MONOMER + 1.0 JCVISYN3A_0655-MONOMER + 1.0 JCVISYN3A_0656-MONOMER
+ 1.0 JCVISYN3A_0657-MONOMER + 1.0 JCVISYN3A_0658-MONOMER + 1.0
JCVISYN3A_0659-MONOMER + 1.0 JCVISYN3A_0660-MONOMER + 1.0 JCVISYN3A_0661-MONOMER
+ 1.0 JCVISYN3A_0662-MONOMER + 1.0 JCVISYN3A_0663-MONOMER + 1.0
JCVISYN3A_0664-MONOMER + 1.0 JCVISYN3A_0665-MONOMER + 1.0 JCVISYN3A_0666-MONOMER
+ 1.0 JCVISYN3A_0667-MONOMER + 1.0 JCVISYN3A_0668-MONOMER + 1.0
```

```
JCVISYN3A_0669-MONOMER + 1.0 JCVISYN3A_0670-MONOMER + 1.0 JCVISYN3A_0671-MONOMER
+ 1.0 JCVISYN3A_0672-MONOMER + 1.0 JCVISYN3A_0806-MONOMER + 1.0
JCVISYN3A_0806-MONOMER_mod_acetyl(2) + 1.0 JCVISYN3A_0807-MONOMER + 1.0
JCVISYN3A_0809-MONOMER + 1.0 JCVISYN3A_0810-MONOMER + 1.0 JCVISYN3A_0833-MONOMER
+ 1.0 JCVISYN3A_0910-MONOMER + 1.0 JCVISYN3A_0913-MONOMER + 1.0
JCVISYN3A_0930-MONOMER + 1.0 JCVISYN3A_0932-MONOMER + 3.0 gtp_c + 2.0 h2o_c +
171.0 mg2_c --> 2.0 gdp_c + 2.0 h_c + 2.0 pi_c + 4.675303663
prosthetic_group_biomass + 1.0 ribosome'
```

[24]:
```
model.metabolites.gtp_c.molecular_weight(numeric_only=False), model.metabolites.
↪gtp_c.formula_weight/1000.
```

[24]:
```
(<Quantity(0.519148663, 'gram / millimole')>, 0.5191486630000001)
```

### 0.0.7 Coupling coefficients are full symbolic expressions

[25]:
```
exp = model.process_data.JCVISYN3A_0004.coupling_coefficient_rna_degradation
exp
```

[25]:

$$\frac{0.333333333333333 f_{mRNA} k_{deg}^{mRNA_JCVISYN3A_0004} m_{aa} \left(r_0 + \frac{\mu}{k_t}\right)}{m_{nt}\mu}$$

[26]:
```
# evaluation using default_parameters results in a function of the growth rate␣
↪only
# this expression is set as a coupling coefficient in COBRAme and coralME v1.0
exp.subs(model.default_parameters)
```

[26]:

$$\frac{0.0269135802469136 \cdot (0.222222222222222\mu + 0.087)}{\mu}$$

[27]:
```
# evaluation using default_parameters and a growth rate
exp.subs(model.default_parameters).subs({ model.mu.magnitude : 0. })
```

[27]:

$$\tilde{\infty}$$

[28]:
```
# easy access to current model growth rate
model.growth_rate, exp.subs(model.default_parameters).subs({ model.mu.magnitude␣
↪: model.growth_rate.magnitude })
```

[28]:
```
(<Quantity(0.310233439, '1 / hour')>, 0.0135282782025462)
```

[29]:
```
# reaction bounds have units
model.reactions[0].bounds
```

[29]:
```
(<Quantity(mu, '1 / hour')>, <Quantity(mu, '1 / hour')>)
```

[30]:
```
# and symbolic coupling coefficients have units
model.reactions.transcription_TU_JCVISYN3A_0004.
↪symbolic_stoichiometry['GreA_mono'].units
```

[30]: dimensionless

### 0.0.8 Extended classes reduce user coding

[31]:
```python
import warnings
from IPython.core.formatters import FormatterWarning
warnings.filterwarnings('ignore', category = FormatterWarning)

# New mappable list class allows for-loops for many properties using the get␣
↪method
# query method returns a mappable list; get method returns a list that is␣
↪easily converted into a DataFrame
# This shows the fluxes of 545 reactions
pandas.DataFrame(model.query(types = [coralme.core.reaction.MetabolicReaction]).
↪get('id', 'flux.magnitude'))
```

[31]:
```
                                      0             1
0             5FTHFabc_REV_CPLX_5FTHFabc-0  0.000000e+00
1             5FTHFabc_FWD_CPLX_5FTHFabc-0  6.514902e-05
2     5FTHFPGS_REV_JCVISYN3A_0823-MONOMER  0.000000e+00
3     5FTHFPGS_FWD_JCVISYN3A_0823-MONOMER  6.514902e-05
4                     ACALDt_REV_CPLX_dummy  0.000000e+00
..                                      …             …
540                   GLNTRATa_REV_GlnS_tri  0.000000e+00
541                   GLNTRATa_FWD_GlnS_tri  7.105781e-02
542                   GLNTRATb_REV_GlnS_tri  0.000000e+00
543                   GLNTRATb_FWD_GlnS_tri  2.615805e-03
544                dummy_reaction_FWD_SPONT  8.820969e-07

[545 rows x 2 columns]
```

[32]:
```python
import warnings
from IPython.core.formatters import FormatterWarning
warnings.filterwarnings('ignore', category = FormatterWarning)

# New mappable list class allows for-loops for many properties using the get␣
↪method
pandas.DataFrame(model.query(types = [coralme.core.component.Metabolite]).
↪get('id', 'formula', 'charge'))
```

[32]:
```
               0             1    2
0    10fthfglu3_c   C30H33N9O13 -4.0
1         12dgr_c      C35H68O5  0.0
2         13dpg_c     C3H4O10P2 -4.0
3         1ag3p_c     C19H37O7P -2.0
4         2dr1p_c       C5H9O7P -2.0
..              …             …    …
```

```
613          tl_c             Tl   NaN
614         cd2_c             Cd   NaN
615       hemed_c  C34H32N4O6Fe   NaN
616      lipoyl_c       C8H12OS2   NaN
617      acetyl_c          C2H2O   NaN

[618 rows x 3 columns]
```

### 0.0.9 New coupling coefficients are instantiated as zero

[33]:
```
parameter_name = model.process_data.tRNA_JCVISYN3A_0070_CUA.
  ↪coupling_coefficient_rna_degradation
model.default_parameters[parameter_name]
```

[33]: 0.0

[34]:
```
parameter_name = model.process_data.rRNA_JCVISYN3A_0067.
  ↪coupling_coefficient_rna_degradation
model.default_parameters[parameter_name]
```

[34]: 0.0

[35]:
```
parameter_name = model.process_data.TU_JCVISYN3A_0004.
  ↪coupling_coefficient_rna_degradation
model.default_parameters[parameter_name]
```

[35]: 0.0

[36]:
```
# default parameters has a query method
# this returns mRNA constant degradation, and TU and protein degradation␣
  ↪coupling coefficients
model.default_parameters.query('JCVISYN3A_0932')
```

[36]:
```
{k_{deg}^{mRNA_JCVISYN3A_0932}: 12.0,
 beta_{tx}^{TU_JCVISYN3A_0932}: 0.0,
 beta_{deg}^{JCVISYN3A_0932-MONOMER}: 0.0}
```

### 0.0.10 Use of token metabolites: GAM+NGAM correction

[37]:
```
# in-solver correction to GAM and NGAM
print('Original bounds*:', model.global_info['gam'], model.global_info['ngam'],␣
  ↪end = '\n\n')

# Step 1
# create a virtual metabolite
tid = 'gam_ngam_token'
token = coralme.core.component.Token(tid)
```

```python
# add it to the model
model.add_metabolites([ token ])

# Step 2
# add it in the RHS of E-reactions consuming ATP and/or GTP
for rxn in model.query(types = [ coralme.core.reaction.TranslationReaction ]):
    # coeff = float(rxn.symbolic_stoichiometry['atp_c'].magnitude.
 ↪as_ordered_terms()[-1]) # This is energy for translation
    coeff = rxn.symbolic_stoichiometry.get('atp_c', 0) # total ATP = tRNA␣
 ↪charging + mRNA degradation
    coeff += rxn.symbolic_stoichiometry.get('gtp_c', 0) # total GTP is addition␣
 ↪of amino acids
    rxn.add_metabolites({ token : -1*coeff }, combine = False) # 3␣
 ↪energy-equivalents per amino acid

for rxn in model.query(types = [ coralme.core.reaction.ComplexDegradation ]):
    coeff = rxn.symbolic_stoichiometry.get('atp_c', 0)
    coeff += rxn.symbolic_stoichiometry.get('gtp_c', 0)
    rxn.add_metabolites({ token : -1*coeff }, combine = False)

for rxn in model.query(types = [ coralme.core.reaction.ComplexFormation ]):
    coeff = rxn.symbolic_stoichiometry.get('atp_c', 0)
    coeff += rxn.symbolic_stoichiometry.get('gtp_c', 0)
    rxn.add_metabolites({ token : -1*coeff }, combine = False)

for rxn in model.query(types = [ coralme.core.reaction.PostTranslationReaction␣
 ↪]):
    coeff = rxn.symbolic_stoichiometry.get('atp_c', 0)
    coeff += rxn.symbolic_stoichiometry.get('gtp_c', 0)
    rxn.add_metabolites({ token : -1*coeff }, combine = False)

for rxn in model.query(types = [ coralme.core.reaction.TranscriptionReaction ]):
    for subrxn in rxn._transcription_data.subreactions: # avoid ATP and GTP␣
 ↪used for mRNA synthesis and degradation
        coeff = model.subreaction_data.get_by_id(subrxn).stoichiometry.
 ↪get('atp_c', 0)
        coeff += model.subreaction_data.get_by_id(subrxn).stoichiometry.
 ↪get('gtp_c', 0)
    rxn.add_metabolites({ token : -1*coeff }, combine = False)

# Step 3
# correct multiplicity of the GAM reaction
model.gam = 1.
# and add the token to the GAM reaction
model.reactions.GAM.add_metabolites({ token : +1. }, combine = False)
# open the reaction bounds
```

```python
model.reactions.GAM.bounds = (0., 1000.)
print('New GAM stoichiometry:', model.reactions.GAM.reaction, end = '\n\n')
# finally, close (or delete) the NGAM reaction
model.reactions.ATPM.bounds = (0., 0.)

# Step 4
# Add a demand reaction for the token (bounds are 0, 1000)
model.metabolites.get_by_id(tid).add_boundary(type = 'demand')
# and set the constraint (here, I'm moving the constraint from the original GAM and NGAM reactions)
lb = model.global_info['gam'] * model.mu #
lb += model.global_info['ngam'] * model.unit_registry.parse_units('1/h')
print('Lower bound:', lb, type(lb), end = '\n\n')
model.reactions.get_by_id('DM_' + tid).bounds = (lb, lb)

# Step 5, optimize
model.optimize()
```

```
Original bounds*: 34.98 1.0

New GAM stoichiometry: 1.0 atp_c + 1.0 h2o_c --> 1.0 adp_c + 1.0 gam_ngam_token
+ 1.0 h_c + 1.0 pi_c

Lower bound: 34.98*mu + 1.0 / hour <class 'pint.Quantity'>

Iteration       Solution to check       Solver Status
---------       ------------------      -------------
        1       1.4050280687025918      Not feasible
        2       0.7025140343512959      Not feasible
        3       0.3512570171756479      Optimal
        4       0.5268855257634719      Not feasible
        5       0.4390712714695599      Not feasible
        6       0.3951641443226039      Optimal
        7       0.4171177078960819      Optimal
        8       0.4280944896828209      Not feasible
        9       0.4226060987894514      Optimal
       10       0.4253502942361362      Optimal
       11       0.4267223919594786      Not feasible
       12       0.4260363430978074      Optimal
       13       0.4263793675286430      Not feasible
       14       0.4262078553132251      Not feasible
       15       0.4261220992055162      Not feasible
       16       0.4260792211516618      Optimal
       17       0.4261006601785891      Optimal
       18       0.4261113796920526      Optimal
       19       0.4261167394487845      Optimal
       20       0.4261194193271504      Optimal
       21       0.4261207592663333      Optimal
```

```
22        0.4261214292359248       Not feasible
```

[37]: True

[38]:
```python
# model explains 55.54% of the GAM+NGAM value at mu 0.1154 per h
model.get('GAM').flux.magnitude / model.get('DM_gam_ngam_token').flux.magnitude␣
↪* 100
```

[38]: 53.34979079870303

### 0.0.11 Protein degradation reactions can be added after model reconstruction

[39]:
```python
# add protein degradation reactions to ME-model
model.remove_reactions(list(model.query(types = [ coralme.core.reaction.
↪ComplexDegradation ])))
for cplx in model.complex_data:
    cplx.create_complex_degradation()
```

```
WARNING: Missing reactions will block flux from
'degradation_PDH_1_CPLX_mod_thmpp(1)_mod_lipoyl(1)_mod_fad(1)'. Check if
degradation or sink of N(6)-lipoyl-L-lysine ('lplys__L_c') is present.
WARNING: Missing reactions will block flux from
'degradation_PDH_2_CPLX_mod_thmpp(1)_mod_lipoyl(1)_mod_fad(1)'. Check if
degradation or sink of N(6)-lipoyl-L-lysine ('lplys__L_c') is present.
INFO: The degradation of 'JCVISYN3A_0621-MONOMER_mod_pan4p(1)' should be carried
out by the acyl carrier protein phosphodiesterase reaction.
WARNING: Missing reactions will block flux from
'degradation_JCVISYN3A_0806-MONOMER_mod_acetyl(2)'. Check if degradation or sink
of N-acetyl-L-alanine ('acala__L_c') is present.
WARNING: Missing reactions will block flux from 'degradation_ribosome'. Check if
degradation or sink of N-acetyl-L-alanine ('acala__L_c') is present.
ERROR: error getting modifications for 'RNA_JCVISYN3A_0356' in reaction
'degradation_rRNA_containing_excision_machinery'.
ERROR: RNA_JCVISYN3A_0356 in reaction
'degradation_rRNA_containing_excision_machinery'
ERROR: error getting modifications for 'RNA_JCVISYN3A_0356' in reaction
'degradation_monocistronic_excision_machinery'.
ERROR: RNA_JCVISYN3A_0356 in reaction
'degradation_monocistronic_excision_machinery'
```

[40]:
```python
len(model.complex_data), len(model.query(types = [ coralme.core.reaction.
↪ComplexDegradation ]))
```

[40]: (266, 265)

### 0.0.12 Add transcription, translation, formation reactions to an ME-model

[41]:
```
from Bio import SeqIO
contigs = []
for contig in SeqIO.parse('genes.fna', 'fasta'):
    contigs.append(contig)
contigs[0].seq
```

[41]:
```
Seq('atgAAAACGAAAATCCCTGATGCGGTATTGGCTGCTGAGGTGAGTCGCCGTGGT…taa')
```

[42]:
```
# running this without a transcrition reaction will create the RNA_<locus tag>␣
↪metabolite
if model.process_data.has_id('b0894'):
    model.process_data.remove('b0894')
if model.reactions.has_id('translation_b0894'):
    model.reactions.remove('translation_b0894')

coralme.util.building.add_translation_reaction(
    model, 'b0894', contigs[0].seq, prot_sequence = contigs[0].translate(11).
↪seq, transl_table=11, update=True)

if model.process_data.has_id('TU_b0894'):
    model.process_data.remove('TU_b0894')
if model.reactions.has_id('transcription_TU_b0894'):
    model.reactions.remove('transcription_TU_b0894')

coralme.util.building.add_transcription_reaction(model, 'TU_b0894', {'b0894',},␣
↪contigs[0].seq, update = True)
```

```
INFO: The 'RNA_b0894' component was not present in ME-model and it was created.
INFO: A TranslationReaction with ID 'translation_b0894' was added to the ME-
model.
INFO: A TranscriptionReaction with ID 'transcription_TU_b0894' was added to the
ME-model.
WARNING: The 'TU_b0894' component has no RNA Polymerase associated to it. It
might be added later during the reconstruction.
```

[43]:
```
if model.process_data.has_id('b0894-MONOMER'):
    model.process_data.remove('b0894-MONOMER')
coralme.util.building.add_complex_to_model(model, 'b0894-MONOMER',␣
↪{'protein_b0894' : 1})
```

[44]:
```
model.query('b0894-MONOMER', types = [coralme.core.reaction.MEReaction])

model.process_data.get_by_id('b0894-MONOMER').create_complex_formation()
model.process_data.get_by_id('b0894-MONOMER').
↪create_complex_degradation(inplace = True)
```

### 0.0.13 Scalar multiplication and addition of reactions stoichiometries allow reaction coupling

[45]:
```
# folding reactions in FoldME and StressME are associated to the Lon␣
↪metalloprotease and normalized to use 2 ATP
scale_factor = 0.8024691358024691 / 65

# normalization to 2 ATP per flux unit
model.global_info['atp_per_proteasome_cut'] = 2 / scale_factor
model.process_data.get_by_id('b0894').aa_per_proteasome_cut = 814. # b0894␣
↪length is 814 aa
model.reactions.get_by_id('degradation_b0894-MONOMER').update()

# this demonstrates water stoichiometry is incorrect in StressME model
stoich = model.reactions.get_by_id('degradation_b0894-MONOMER').metabolites *␣
↪scale_factor

# remove protease and proteasome, and assign Lon metalloprotease
stoich.pop(model.get('generic_protease'))
stoich.pop(model.get('generic_proteasome'))

# catalytic efficiency is 196,000 cuts per second, and the stoichometric␣
↪coefficient must be dimensionless
stoich.update({ model.get('Lon') : 1/196000*3600 * model.unit_registry.
↪parse_units('h') * model.mu * scale_factor })
# direct assignment; equivalent to reaction.add_metabolites(combine=False)
model.reactions.get_by_id('degradation_b0894-MONOMER')._metabolites = stoich
model.reactions.get_by_id('degradation_b0894-MONOMER').metabolites
```

[45]:
```
{<Complex b0894-MONOMER at 0x7bb1a7fe00e0>: -0.012345679012345678,
 <Metabolite atp_c at 0x7bb1a4c56f30>: -2.0,
 <Metabolite h2o_c at 0x7bb1a6fa9310>: -12.037037037037036,
 <Metabolite adp_c at 0x7bb1abe8aea0>: 2.0,
 <Metabolite h_c at 0x7bb19fe977a0>: 2.0,
 <Metabolite pi_c at 0x7bb1a0dd9760>: 2.0,
 <Metabolite ala__L_c at 0x7bb1a118fce0>: 0.8024691358024691,
 <Metabolite arg__L_c at 0x7bb1a118c740>: 0.6049382716049383,
 <Metabolite asn__L_c at 0x7bb1a118d550>: 0.48148148148148145,
 <Metabolite asp__L_c at 0x7bb1a118d3a0>: 0.5802469135802468,
 <Metabolite cys__L_c at 0x7bb1a6d603b0>: 0.18518518518518517,
 <Metabolite gln__L_c at 0x7bb1ab269ca0>: 0.35802469135802467,
 <Metabolite glu__L_c at 0x7bb1af145190>: 0.6296296296296295,
 <Metabolite gly_c at 0x7bb1a118df70>: 0.8888888888888888,
 <Metabolite his__L_c at 0x7bb1a118f1a0>: 0.20987654320987653,
 <Metabolite ile__L_c at 0x7bb1a118f500>: 0.6790123456790123,
 <Metabolite leu__L_c at 0x7bb1a118f7d0>: 0.6790123456790123,
 <Metabolite lys__L_c at 0x7bb1a118f8c0>: 0.5432098765432098,
```

```
<Metabolite met__L_c at 0x7bb1a118fa40>: 0.30864197530864196,
<Metabolite phe__L_c at 0x7bb1a118fd10>: 0.2222222222222222,
<Metabolite pro__L_c at 0x7bb1a118fda0>: 0.5679012345679012,
<Metabolite ser__L_c at 0x7bb1a118c050>: 0.5432098765432098,
<Metabolite thr__L_c at 0x7bb1a118d9a0>: 0.6049382716049383,
<Metabolite trp__L_c at 0x7bb1a8aafec0>: 0.16049382716049382,
<Metabolite tyr__L_c at 0x7bb1ac3dc9e0>: 0.43209876543209874,
<Metabolite val__L_c at 0x7bb1ac3df2f0>: 0.5679012345679012,
<Constraint protein_biomass at 0x7bb1a0eb0b60>: -1.1159577491358024,
<Complex Lon at 0x7bb19f074260>: 0.000226757369614512*mu}
```

### 0.0.14 New reaction, metabolites, and process data properties

[46]:
```
# determine simulation artifact(s): reactions that convert a metabolite-limited␣
↪into a proteome-limited ME-model
pandas.DataFrame(coralme.core.extended_classes.MappableList(model.
↪stoichiometric_data).get('id', 'artifact'))
```

[46]:
```
                   0     1
0          5FTHFabc  None
1          5FTHFPGS  None
2            ACALDt  None
3              ACKr  None
4            ACPPAT  None
..              ...   ...
198            NH3c  None
199             PDH  None
200        GLNTRATa  None
201        GLNTRATb  None
202  dummy_reaction  None

[203 rows x 2 columns]
```

[47]:
```
pandas.DataFrame(model.query('MAN6PI|UDPGALM', types = [ coralme.core.reaction.
↪MetabolicReaction ]).get('reaction', 'flux'))
```

[47]:
```
                                                    0  \
0  [5.21851832759033e-6*mu] JCVISYN3A_0435-MONOME…
1  [5.21851832759033e-6*mu] JCVISYN3A_0435-MONOME…
2  [4.24293808771141e-6*mu] JCVISYN3A_0814-MONOME…
3  [4.24293808771141e-6*mu] JCVISYN3A_0814-MONOME…

                                         1
0                  0.0 millimole / gram / hour
1                  0.0 millimole / gram / hour
2  817.0375540722313 millimole / gram / hour
3  817.2057196470305 millimole / gram / hour
```

### 0.0.15 model.get defaults to model.query if multiple results match the query (case insensitive)

[48]:
```
model.get('Generic', mappable = False) == model.get('generic', mappable =␣
↪False) # get method is case insensitive
```

[48]: True

### 0.0.16 Working with the unit system

[49]:
```
import coralme, sympy
model = coralme.io.pickle.load_pickle_me_model('iMmy281-ME.pkl')

# define custom units if needed and their dimensions
for unit in ['gene', 'cell', 'initiations']:
    model.unit_registry.define(f'[{unit}]')
    model.unit_registry.define(f'{unit} = [{unit}]')

# create new symbols, parameters, and set them as bounds or coupling␣
↪coefficients
# Thiele et al., 2009, used maximum transcription initiation rates v_stable␣
↪RNA,max,i

# copy number of the stable RNA gene i per cell due to gene dosage
model.symbols['16S'] = sympy.Symbol('16S', positive = True) * model.
↪unit_registry.parse_units('gene per cell')

# rRNA transcription initiation rate
model.symbols['i_rrn'] = sympy.Symbol('i_rrn', positive = True) * model.
↪unit_registry.parse_units('initiations per minute per gene')

# doubling time
model.symbols['T_D'] = sympy.Symbol('T_D', positive = True) * model.
↪unit_registry.parse_units('minute')

model.symbols['v_{stable RNA,max,i}'] = model.symbols['16S'] * model.
↪symbols['i_rrn'] * model.symbols['T_D']

model.default_parameters['16S'] = 4.49 # Table 3
model.default_parameters['i_rrn'] = 39 # initiations/min/gene at T_D = 30
model.default_parameters['T_D'] = 30 # min

model.symbols['v_{stable RNA,max,i}'], model.symbols['v_{stable RNA,max,i}'].
↪subs(model.default_parameters)
```

[49]:
```
(<Quantity(16S*T_D*i_rrn, 'initiations / cell')>, 5253.30000000000)
```

[50]:
```python
import coralme, sympy
model = coralme.io.pickle.load_pickle_me_model('iMmy281-ME.pkl')

# check if unit registry already contains custom units
for name in ['gene', 'cell', 'initiations']:
    print(name, model.unit_registry._units.get(name))
for name in ['[gene]', '[cell]', '[initiations]']:
    print(name, model.unit_registry._dimensions.get(name))

# if they are defined, we should remove them
for unit in ['gene', 'cell', 'initiations']:
    model.unit_registry._units.pop(unit, None)
for dimension in ['[gene]', '[cell]', '[initiations]']:
    model.unit_registry._dimensions.pop(dimension, None)

# define custom units if needed and their dimensions
for unit in ['gene', 'cell', 'initiations']:
    model.unit_registry.define(f'[{unit}]')
    model.unit_registry.define(f'{unit} = [{unit}]')

# create new symbols, parameters, and set them as bounds or coupling␣
↪coefficients
# Thiele et al., 2009, used maximum transcription initiation rates v_stable␣
↪RNA,max,i

# copy number of the stable RNA gene i per cell due to gene dosage
model.symbols['16S'] = sympy.Symbol('16S', positive = True) * model.
↪unit_registry.parse_units('gene per cell')

# rRNA transcription initiation rate
model.symbols['i_rrn'] = sympy.Symbol('i_rrn', positive = True) * model.
↪unit_registry.parse_units('initiations per minute per gene')

# doubling time
model.symbols['T_D'] = sympy.Symbol('T_D', positive = True) * model.
↪unit_registry.parse_units('minute')

model.symbols['v_{stable RNA,max,i}'] = model.symbols['16S'] * model.
↪symbols['i_rrn'] * model.symbols['T_D']

model.default_parameters['16S'] = 4.49 # Table 3
model.default_parameters['i_rrn'] = 39 # initiations/min/gene at T_D = 30
model.default_parameters['T_D'] = 30 # min

model.symbols['v_{stable RNA,max,i}'], model.symbols['v_{stable RNA,max,i}'].
↪subs(model.default_parameters)
```

```
gene UnitDefinition(name='gene', defined_symbol=None, aliases=(),
converter=ScaleConverter(scale=1), reference=<UnitsContainer({'[gene]': 1})>,
start_line=1, start_col=0, end_line=1, end_col=13, raw='gene = [gene]')
cell UnitDefinition(name='cell', defined_symbol=None, aliases=(),
converter=ScaleConverter(scale=1), reference=<UnitsContainer({'[cell]': 1})>,
start_line=1, start_col=0, end_line=1, end_col=13, raw='cell = [cell]')
initiations UnitDefinition(name='initiations', defined_symbol=None, aliases=(),
converter=ScaleConverter(scale=1), reference=<UnitsContainer({'[initiations]':
1})>, start_line=1, start_col=0, end_line=1, end_col=27, raw='initiations =
[initiations]')
[gene] DimensionDefinition(name='[gene]', start_line=1, start_col=0, end_line=1,
end_col=6, raw='[gene]')
[cell] DimensionDefinition(name='[cell]', start_line=1, start_col=0, end_line=1,
end_col=6, raw='[cell]')
[initiations] DimensionDefinition(name='[initiations]', start_line=1,
start_col=0, end_line=1, end_col=13, raw='[initiations]')
```

[50]:
```
(<Quantity(16S*T_D*i_rrn, 'initiations / cell')>, 5253.30000000000)
```

[51]:
```
model.reactions.transcription_TU_JCVISYN3A_0067.bounds # A mmol of biomass is␣
↪equivalent to 1 gram of biomass
```

[51]:
```
(<Quantity(0.0, 'millimole / gram / hour')>,
 <Quantity(1000.0, 'millimole / gram / hour')>)
```

[52]:
```
# although 'v_{stable RNA,max,i}' has wrong units (it should be mmol per gDCW␣
↪per hour), we can use it as reaction bounds
model.reactions.transcription_TU_JCVISYN3A_0067.bounds = (model.
↪symbols['v_{stable RNA,max,i}'], model.symbols['v_{stable RNA,max,i}'])
```

[53]:
```
model.reactions.transcription_TU_JCVISYN3A_0067.bounds
```

[53]:
```
(<Quantity(16S*T_D*i_rrn, 'initiations / cell')>,
 <Quantity(16S*T_D*i_rrn, 'initiations / cell')>)
```

### 0.0.17 New MEModelWrapper for convenient access to model components

[54]:
```
test = coralme.core.extended_classes.MEModelWrapper(model)
```

[55]:
```
test.transcription_TU_JCVISYN3A_0067 == test.reactions.
↪transcription_TU_JCVISYN3A_0067
```

[55]:
```
True
```

### 0.0.18 Download ME-model input files and reconstruct for selected organisms

[56]:
```
from coralme.util.agora import list_available_models, get_model_from_manifest,␣
↪get_manifest, download_file
list_available_models()[0]
```

[56]: 'AGORA-Abiotrophia_defectiva_ATCC_49176-ME'

[57]:
```
agora = get_model_from_manifest(
    model_key='AGORA-Abiotrophia_defectiva_ATCC_49176-ME', troubleshoot = False,
    extra_configuration = {
        'relax_rrna_modifications' : True,
        'allow_massless_biomass_precursors' : True
    })
```

```
Initiating file processing…
~ Processing files for Abiotrophia_defectiva_ATCC_49176-ME…
Reading Abiotrophia_defectiva_ATCC_49176-ME done.
File processing done.
Initiating ME-model reconstruction…
ME-model was saved in the models/AGORA-Abiotrophia_defectiva_ATCC_49176-ME
directory as MEModel-step1-Abiotrophia_defectiva_ATCC_49176-ME.pkl
ME-model was saved in the models/AGORA-Abiotrophia_defectiva_ATCC_49176-ME
directory as MEModel-step2-Abiotrophia_defectiva_ATCC_49176-ME.pkl
ME-model reconstruction is done.
Number of metabolites in the ME-model is 2975 (+208.29%, from 965)
Number of reactions in the ME-model is 6234 (+494.85%, from 1048)
Number of genes in the ME-model is 617 (+-3.44%, from 639)
Number of missing genes from reconstruction cannot be determined.
```

[58]:
```
agora.me_model.remove_reactions(list(agora.me_model.query('sink')))
```

[59]:
```
df = pandas.read_csv('AGORADiets.txt', index_col = 1, sep = ' ')
diet_dct = coralme.core.diet.parse_diet(df, column = 'WesternDiet')
coralme.core.diet.constrain_diet(agora.me_model, diet_dct)

biodf = pandas.read_csv('all_biomass.csv', index_col = 0)
coralme.util.agora.correct_biomass(agora.me_model, biodf.
↪loc['Abiotrophia_defectiva_ATCC_49176'])

agora.troubleshoot()
```

```
The following biomass constituents are not in the ME-model or have no formula:
colipa_c, glyc45tca_c, glyc45tcaglc_c, tcam_c, glyc45tcaala__D_c.
The MINOS and quad MINOS solvers are a courtesy of Prof Michael A. Saunders.
Please cite Ma, D., Yang, L., Fleming, R. et al. Reliable and efficient solution
of genome-scale models of Metabolism and macromolecular Expression. Sci Rep 7,
40863 (2017). https://doi.org/10.1038/srep40863
```

```
~ Troubleshooting started…
  Checking if the coralME model can simulate growth without gapfilling
reactions…
  Original coralME model is not feasible with a tested growth rate of 0.001000
1/h
  Step 1. Gapfill reactions to provide components of type 'ME-Deadends' using
brute force.
          Finding gaps in the ME-model…
          Finding gaps from the M-model only…
          9 metabolites were identified as deadends.
            PGP_c: Peptidoglycan polymer (n subunits)
            cs_c: Missing metabolite in the M-model.
            dmbzid_c: 5,6-Dimethylbenzimidazole
            dnarep_c: DNA replication
            glycyl_c: Missing metabolite in the M-model.
            ni2_c: Missing metabolite in the M-model.
            proteinsynth_c: Protein biosynthesis
            rnatrans_c: RNA transcription
            tl_c: Missing metabolite in the M-model.
          Adding sink reactions for 9 components…
          Provided metabolites through sink reactions cannot recover growth.
Proceeding to next set of metabolites.
  Step 2. Gapfill reactions to provide components of type 'Biomass' using brute
force.
          Adding sink reactions for 44 components…
            Sink reactions shortlisted to 37 metabolites.
             Processed:  1/37, Gaps: 0. The ME-model is feasible if TS_2dmmq8_c
is closed.
             Processed:  2/37, Gaps: 0. The ME-model is feasible if
TS_i17tcaala__D_c is closed.
             Processed:  3/37, Gaps: 0. The ME-model is feasible if
TS_i17tcaglc_c is closed.
             Processed:  4/37, Gaps: 0. The ME-model is feasible if TS_ribflv_c
is closed.
             Processed:  5/37, Gaps: 0. The ME-model is feasible if TS_pgai17_c
is closed.
             Processed:  6/37, Gaps: 0. The ME-model is feasible if TS_gthrd_c
is closed.
             Processed:  7/37, Gaps: 0. The ME-model is feasible if TS_pheme_c
is closed.
             Processed:  8/37, Gaps: 0. The ME-model is feasible if TS_pe180_c
is closed.
             Processed:  9/37, Gaps: 0. The ME-model is feasible if TS_pydx5p_c
is closed.
             Processed: 10/37, Gaps: 0. The ME-model is feasible if TS_peai17_c
is closed.
             Processed: 11/37, Gaps: 0. The ME-model is feasible if
TS_i17tcaacgam_c is closed.
```

```
            Processed: 12/37, Gaps: 0. The ME-model is feasible if TS_pei17_c
is closed.
            Processed: 13/37, Gaps: 0. The ME-model is feasible if TS_clpn180_c
is closed.
            Processed: 14/37, Gaps: 0. The ME-model is feasible if
TS_sttcaacgam_c is closed.
            Processed: 15/37, Gaps: 0. The ME-model is feasible if TS_thmpp_c
is closed.
            Processed: 16/37, Gaps: 1. The ME-model is not feasible if
TS_adocbl_c is closed.
            Processed: 17/37, Gaps: 1. The ME-model is feasible if TS_i17tca1_c
is closed.
            Processed: 18/37, Gaps: 1. The ME-model is feasible if TS_sheme_c
is closed.
            Processed: 19/37, Gaps: 1. The ME-model is feasible if TS_pg180_c
is closed.
            Processed: 20/37, Gaps: 1. The ME-model is feasible if
TS_ai17tca1_c is closed.
            Processed: 21/37, Gaps: 1. The ME-model is feasible if TS_ptrc_c is
closed.
            Processed: 22/37, Gaps: 1. The ME-model is feasible if TS_mqn8_c is
closed.
            Processed: 23/37, Gaps: 1. The ME-model is feasible if TS_sttca1_c
is closed.
            Processed: 24/37, Gaps: 1. The ME-model is feasible if
TS_sttcaglc_c is closed.
            Processed: 25/37, Gaps: 1. The ME-model is feasible if TS_udcpdp_c
is closed.
            Processed: 26/37, Gaps: 1. The ME-model is feasible if TS_clpni17_c
is closed.
            Processed: 27/37, Gaps: 1. The ME-model is feasible if
TS_clpnai17_c is closed.
            Processed: 28/37, Gaps: 1. The ME-model is feasible if
TS_sttcaala__D_c is closed.
            Processed: 29/37, Gaps: 1. The ME-model is feasible if TS_cobalt2_c
is closed.
            Processed: 30/37, Gaps: 1. The ME-model is feasible if TS_spmd_c is
closed.
            Processed: 31/37, Gaps: 1. The ME-model is feasible if TS_amet_c is
closed.
            Processed: 32/37, Gaps: 1. The ME-model is feasible if
TS_ai17tcaacgam_c is closed.
            Processed: 33/37, Gaps: 1. The ME-model is feasible if
TS_ai17tcaglc_c is closed.
            Processed: 34/37, Gaps: 1. The ME-model is feasible if TS_pgi17_c
is closed.
            Processed: 35/37, Gaps: 1. The ME-model is feasible if
TS_ai17tcaala__D_c is closed.
```

```
             Processed: 36/37, Gaps: 1. The ME-model is feasible if TS_PGP_c is
closed.
             Processed: 37/37, Gaps: 1. The ME-model is feasible if TS_glycyl_c
is closed.
~ Troubleshooter added the following sinks: TS_adocbl_c.
~ Final step. Fully optimizing with precision 1e-6 and save solution into the
coralME model…
  Gapfilled coralME model is feasible with growth rate 0.038990 (M-model:
55.651038).
coralME model was saved in the models/AGORA-Abiotrophia_defectiva_ATCC_49176-ME
directory as MEModel-step3-Abiotrophia_defectiva_ATCC_49176-ME-TS.pkl
```
