## Supplementary File S2 for "Expanded ME-model analysis capabilities enable genetic recoding, proteostasis, RNA decay, and cofactor essentiality in bacterial models"

November 3, 2026

[1]:
```
import itables
itables.init_notebook_mode(all_interactive=True)
```

```
<IPython.core.display.HTML object>
```

```
<IPython.core.display.HTML object>
```

[2]:
```
import os
os.environ["TQDM_DISABLE"] = "1"
from tqdm import tqdm
```

[3]:
```
# Install dependencies
# !{sys.executable} -m pip install -r requirements.txt
```

[4]:
```
import sys
sys.path.insert(0, './Supplementary-File-S8')
import coralme
coralme.__file__.split('Supplementary-File-S8')[1]
```

[4]:
```
'/coralme/__init__.py'
```

[5]:
```
import coralme
# WARNING: Run again after copy-pasting files from manual-curation file
def generate_files(organism):
    try:
        builder = coralme.builder.main.MEBuilder(me_model_id = organism,␣
↪config_files=['1.input+manual.json', '2.organism+manual.json'])
        builder.configuration['df_gene_cplxs_mods_rxns'] = 'GRO-Ecoli-ME-{:
↪s}-noBBH/automated-org-with-refs-tRNAs.xlsx'.format(organism)
        # Original GenBank was modified to have WT Ecoli locus tags
        builder.configuration['genbank-path'] = 'blastn-and-blastp-bbh-outputs/
↪ASM584_to_{:s}/combined-{:s}-pEVOL-pBpF.gb'
        builder.configuration['genbank-path'] = builder.
↪configuration['genbank-path'].format(organism, organism)

        if organism == 'ASM584v2':
            builder.configuration['genbank-path'] = 'dna-sequences/ASM584v2␣
↪(wildtype)/genomic.gbff'
```

```
        builder.configuration['START_tRNA'] = ['b2814', 'b2815', 'b2816',␣
↪'b3171']
        # set this to True the first time, then copy-paste information from␣
↪reference Ecoli ME-model
        builder.configuration['dev_reference'] = True
        builder.configuration['run_bbh_blast'] = False
        builder.configuration['ncRNAs'] = { "ffs" : "b0455", "rnpB" :␣
↪"b3123" }

    elif organism == 'rE.coli-57':
        # rE.coli-57 IDs after PGAP
        builder.configuration['START_tRNA'] = ['pgaptmp_002345',␣
↪'pgaptmp_002346', 'pgaptmp_002347', 'pgaptmp_002670']
        # set this to True the first time, then copy-paste information from␣
↪reference Ecoli ME-model
        builder.configuration['dev_reference'] = False
        builder.configuration['run_bbh_blast'] = False
        builder.configuration['ncRNAs'] = { "ffs" : "pgaptmp_000364",␣
↪"rnpB" : "pgaptmp_002621" }

    elif organism == 'C321.deltaA':  # ASM47403v1
        builder.configuration['START_tRNA'] = ['N840_2869', 'N840_2870',␣
↪'N840_2871', 'N840_3223']
        # set this to True the first time, then copy-paste information from␣
↪reference Ecoli ME-model
        builder.configuration['dev_reference'] = False
        builder.configuration['run_bbh_blast'] = False
        builder.configuration['ncRNAs'] = { "ffs" : "N840_0460", "rnpB" :␣
↪"N840_3173" }

    elif organism == 'rEc.b.dC.12':
        builder.configuration['include_pseudo_genes'] = True
        # rEc.b.dC.12 IDs after PGAP
        builder.configuration['START_tRNA'] = ['pgaptmp_002925',␣
↪'pgaptmp_002926', 'pgaptmp_002927', 'pgaptmp_003283']
        # set this to True the first time, then copy-paste information from␣
↪reference Ecoli ME-model
        builder.configuration['dev_reference'] = False
        builder.configuration['run_bbh_blast'] = False
        builder.configuration['ncRNAs'] = { "ffs" : "pgaptmp_000469",␣
↪"rnpB" : "pgaptmp_003234" }

    elif organism == 'rEc.y.dC.46':
        builder.configuration['include_pseudo_genes'] = True
        # rEc.y.dC.46 IDs after PGAP
```

```
            builder.configuration['START_tRNA'] = ['pgaptmp_002924',␣
↪'pgaptmp_002925', 'pgaptmp_002926', 'pgaptmp_003281']
            # set this to True the first time, then copy-paste information from␣
↪reference Ecoli ME-model
            builder.configuration['dev_reference'] = False
            builder.configuration['run_bbh_blast'] = False
            builder.configuration['ncRNAs'] = { "ffs" : "pgaptmp_000469",␣
↪"rnpB" : "pgaptmp_003232" }

        elif organism == 'ALE-C321.deltaA':
            builder.configuration['START_tRNA'] = ['CXP41_RS15410',␣
↪'CXP41_RS15415', 'CXP41_RS15420', 'CXP41_RS17310']
            # set this to True the first time, then copy-paste information from␣
↪reference Ecoli ME-model
            builder.configuration['dev_reference'] = False
            builder.configuration['run_bbh_blast'] = False
            builder.configuration['ncRNAs'] = { "ffs" : "CXP41_RS02480", "rnpB"␣
↪: "CXP41_RS17060" }

        elif organism == 'Ochre':
            builder.configuration['START_tRNA'] = ['ACI6BN_20510',␣
↪'ACI6BN_18735']
            # set this to True the first time, then copy-paste information from␣
↪reference Ecoli ME-model
            builder.configuration['dev_reference'] = False
            builder.configuration['run_bbh_blast'] = False
            builder.configuration['ncRNAs'] = { "ffs" : "ACI6BN_10435", "rnpB" :
↪ "ACI6BN_18980" }

        else:
            raise ValueError('No Model ID selected.')

        builder.configuration['m-model-path'] = "inputs/iJO1366b-coralme.json"
        builder.configuration['ME-Model-ID'] = 'GRO-Ecoli-ME-{:s}-noBBH'.
↪format(organism)
        builder.configuration['log_directory'] = 'GRO-Ecoli-ME-{:s}-noBBH'.
↪format(organism)
        builder.configuration['out_directory'] = 'GRO-Ecoli-ME-{:s}-noBBH'.
↪format(organism)

        # do no use transcription units coordinates from reference Ecoli␣
↪ME-model
        builder.configuration['df_TranscriptionalUnits'] = ''

        builder.configuration['add_lipoproteins'] = True
        # do not use biocyc data from reference Ecoli ME-model
```

```
        builder.configuration['biocyc.genes'] = ''
        builder.configuration['biocyc.prots'] = ''
        builder.configuration['biocyc.TUs'] = ''
        builder.configuration['biocyc.RNAs'] = ''
        builder.configuration['biocyc.seqs'] = ''

        builder.generate_files(overwrite = True)

        # this trna_to_codon.txt requires translation to new RNA IDs
        with open('trna_to_codon.txt', 'r') as infile:
            source = pandas.read_csv(infile, sep = '\t', header = None).
↪set_index(0).to_dict()[1]
        # source

        with open('GRO-Ecoli-ME-{:s}-noBBH/automated-org-with-refs-tRNAs.xlsx'.
↪format(organism), 'rb') as infile:
            target = pandas.read_excel(infile)
        target['tRNA-codon association'] = target.apply(lambda x: source.
↪get(x['Gene Locus ID'], None), axis = 1)

        # also, delete PrfA associated data
        if organism in [ 'rEc.b.dC.12', 'rEc.y.dC.46' ]:
            # b1211 works because we're using the same building_data as the␣
↪reference model
            idx = target[target['Gene Locus ID'] == 'b1211'].index
            target = target.drop(idx)

        # save to file
        output = 'GRO-Ecoli-ME-{:s}-noBBH/automated-org-with-refs-tRNAs.xlsx'.
↪format(organism)
        coralme.builder.preprocess_inputs._save_to_excel(target, output,␣
↪builder.me_model.reconstruction_time)
        return builder, target
    except Exception:
        import traceback
        traceback.print_exc()

    return builder
```

[6]:
```
# # WARNING: if re-running build_me_model, copy-paste genome sequence into␣
↪building_data
# for organism in [ 'rEc.y.dC.46' ]:
#     res = generate_files(organism)
```

[7]:
```
# WARNING: if re-running build_me_model, copy-paste genome sequence into␣
↪building_data
```

```
import multiprocessing
models = [ 'ASM584v2', 'rE.coli-57', 'C321.deltaA', 'ALE-C321.deltaA', 'rEc.b.
  ↪dC.12', 'rEc.y.dC.46', 'Ochre' ]
with multiprocessing.Pool(max(len(models), multiprocessing.cpu_count()-1)) as␣
  ↪pool:
    res = pool.map(generate_files, models)
```

```
Initiating file processing…Initiating file processing…

~ Processing files for GRO-Ecoli-ME-rE.coli-57-noBBH…
~ Processing files for GRO-Ecoli-ME-Ochre-noBBH…
Initiating file processing…
~ Processing files for GRO-Ecoli-ME-rEc.b.dC.12-noBBH…
Initiating file processing…
~ Processing files for GRO-Ecoli-ME-ASM584v2-noBBH…
Initiating file processing…
~ Processing files for GRO-Ecoli-ME-rEc.y.dC.46-noBBH…
Initiating file processing…Initiating file processing…

~ Processing files for GRO-Ecoli-ME-C321.deltaA-noBBH…~ Processing files for
GRO-Ecoli-ME-ALE-C321.deltaA-noBBH…

Reading GRO-Ecoli-ME-rE.coli-57-noBBH done.
Reading GRO-Ecoli-ME-Ochre-noBBH done.
Reading GRO-Ecoli-ME-C321.deltaA-noBBH done.
Writing the Organism-Specific Matrix to GRO-Ecoli-ME-rE.coli-57-noBBH/automated-
org-with-refs-tRNAs.xlsx…
Reading GRO-Ecoli-ME-ASM584v2-noBBH done.
~ Processing files for iJL1678b…
Writing the Organism-Specific Matrix to GRO-Ecoli-ME-Ochre-noBBH/automated-org-
with-refs-tRNAs.xlsx…
Reading GRO-Ecoli-ME-ALE-C321.deltaA-noBBH done.
Reading iJL1678b done.
Reading GRO-Ecoli-ME-rEc.y.dC.46-noBBH done.
Writing the Organism-Specific Matrix to GRO-Ecoli-ME-C321.deltaA-
noBBH/automated-org-with-refs-tRNAs.xlsx…
Reading GRO-Ecoli-ME-rEc.b.dC.12-noBBH done.
Writing the Organism-Specific Matrix to GRO-Ecoli-ME-
rEc.y.dC.46-noBBH/automated-org-with-refs-tRNAs.xlsx…
Writing the Organism-Specific Matrix to GRO-Ecoli-ME-ALE-C321.deltaA-
noBBH/automated-org-with-refs-tRNAs.xlsx…
Writing the Organism-Specific Matrix to GRO-Ecoli-ME-
rEc.b.dC.12-noBBH/automated-org-with-refs-tRNAs.xlsx…
Writing the Organism-Specific Matrix to GRO-Ecoli-ME-ASM584v2-noBBH/automated-
org-with-refs-tRNAs.xlsx…
Organism-Specific Matrix saved to GRO-Ecoli-ME-rE.coli-57-noBBH/automated-org-
with-refs-tRNAs.xlsx file.
```

```
File processing done.
Organism-Specific Matrix saved to GRO-Ecoli-ME-Ochre-noBBH/automated-org-with-
refs-tRNAs.xlsx file.
File processing done.
Organism-Specific Matrix saved to GRO-Ecoli-ME-C321.deltaA-noBBH/automated-org-
with-refs-tRNAs.xlsx file.
File processing done.
Organism-Specific Matrix saved to GRO-Ecoli-ME-rEc.b.dC.12-noBBH/automated-org-
with-refs-tRNAs.xlsx file.
Organism-Specific Matrix saved to GRO-Ecoli-ME-rEc.y.dC.46-noBBH/automated-org-
with-refs-tRNAs.xlsx file.
File processing done.
File processing done.
Organism-Specific Matrix saved to GRO-Ecoli-ME-ALE-C321.deltaA-noBBH/automated-
org-with-refs-tRNAs.xlsx file.
File processing done.
Organism-Specific Matrix saved to GRO-Ecoli-ME-ASM584v2-noBBH/automated-org-
with-refs-tRNAs.xlsx file.
File processing done.
```

[8]:
```
builder, target = res[-3]
builder.me_model.id
```

[8]:
```
'rEc.b.dC.12'
```

[9]:
```
# check renamed locus tags
target[target['Gene Locus ID'] == 'b0893'].dropna(how = 'all', axis = 1)
```

[9]:
```
     Gene Locus ID           Definition Gene Names Feature Type    Complex ID
Cofactors in Modified Complex Generic Complex ID
5148         b0893  serine--tRNA ligase       serS          CDS  Ser_RS_dim:2
NaN          cys_ligase
5149         b0893  serine--tRNA ligase       serS          CDS  Ser_RS_dim:2
mg2(1)                  NaN
```

[10]:
```
# check presence of bipA transport reactions
tmp = target[target['M-model Reaction ID'].notna() & target['M-model Reaction␣
↪ID'].str.contains('BIPA')]
tmp.dropna(how = 'all', axis = 1).drop_duplicates()
```

[10]:
```
    Gene Locus ID
Definition Gene Names Feature Type       Complex ID M-model Reaction ID  \
718         b3454  high-affinity branched-chain amino acid ABC transporter ATP-
binding protein LivF       livF           CDS    ABC-15-CPLX:1
BIPAabcpp
719         b3455  high-affinity branched-chain amino acid ABC transporter ATP-
binding protein LivG       livG           CDS    ABC-15-CPLX:1
BIPAabcpp
```

```
720         b3456                           branched chain amino acid ABC
transporter permease LivM       livM          CDS    ABC-15-CPLX:1
BIPAabcpp
721         b3457              high-affinity branched-chain amino acid ABC
transporter permease LivH       livH          CDS    ABC-15-CPLX:1
BIPAabcpp
722         b3460           branched chain amino acid ABC transporter substrate-
binding protein LivJ       livJ          CDS    ABC-15-CPLX:1
BIPAabcpp
723         b4208
D-serine/D-alanine/glycine transporter       cycA          CDS   CYCA-MONOMER:1
BIPAt2pp
724         b0241
phosphoporin PhoE       phoE          CDS     CPLX0-7530:3            BIPAtex
725         b0929
porin OmpF       ompF          CDS     CPLX0-7534:3            BIPAtex
726         b1377
porin OmpN       ompN          CDS   G6700-MONOMER:1            BIPAtex
727         b2215
porin OmpC       ompC          CDS     CPLX0-7533:3            BIPAtex

                                                          Reaction Name
Reversibility
718                    L-biphenylalanine transport via ABC system (periplasm)
True
719                    L-biphenylalanine transport via ABC system (periplasm)
True
720                    L-biphenylalanine transport via ABC system (periplasm)
True
721                    L-biphenylalanine transport via ABC system (periplasm)
True
722                    L-biphenylalanine transport via ABC system (periplasm)
True
723             L-biphenylalanine transport in via proton symport (periplasm)
False
724  L-biphenylalanine transport via diffusion (extracellular to periplasm)
True
725  L-biphenylalanine transport via diffusion (extracellular to periplasm)
True
726  L-biphenylalanine transport via diffusion (extracellular to periplasm)
True
727  L-biphenylalanine transport via diffusion (extracellular to periplasm)
True
```

[11]:
```
# check presence of tRNA(bipA) synthetase
tmp = target[target['Complex ID'].notna() & target['Complex ID'].str.
 ↪contains('bipA_tRNA_synth_copy1-MONOMER')]
```

```
tmp.dropna(how = 'all', axis = 1)
```

[11]:
```
                Gene Locus ID Feature Type                       Complex ID
9163  bipA_tRNA_synth_copy1          CDS  bipA_tRNA_synth_copy1-MONOMER:1
```

[12]:
```
# check problematic complexes
cplxs = 'CPLX0-7719|CPLX0-7761|DIHYDROXYACIDDEHYDRAT-CPLX|SULFITE-REDUCT-CPLX'
target[target['Complex ID'].str.contains(cplxs, na = False)]
```

[12]:
```
     Gene Locus ID
Definition Gene Names  Old Locus Tag  BioCyc  Reference BBH  Reference
Definition Feature Type  Complex Name  \
355          b0118    bifunctional aconitate hydratase 2/2-methylisocitrate
dehydratase       acnB            NaN     NaN            NaN
NaN          CDS           NaN
357          b0118    bifunctional aconitate hydratase 2/2-methylisocitrate
dehydratase       acnB            NaN     NaN            NaN
NaN          CDS           NaN
1131         b3771                                          dihydroxy-acid
dehydratase       ilvD            NaN     NaN            NaN
NaN          CDS           NaN
1132         b3771                                          dihydroxy-acid
dehydratase       ilvD            NaN     NaN            NaN
NaN          CDS           NaN
1398         b2763          assimilatory sulfite reductase (NADPH) hemoprotein
subunit        cysI            NaN     NaN            NaN                   NaN
CDS           NaN
1399         b2764  NADPH-dependent assimilatory sulfite reductase flavoprotein
subunit        cysJ            NaN     NaN            NaN                   NaN
CDS           NaN
1521         b2763          assimilatory sulfite reductase (NADPH) hemoprotein
subunit        cysI            NaN     NaN            NaN                   NaN
CDS           NaN
1522         b2764  NADPH-dependent assimilatory sulfite reductase flavoprotein
subunit        cysJ            NaN     NaN            NaN                   NaN
CDS           NaN
1529         b2763          assimilatory sulfite reductase (NADPH) hemoprotein
subunit        cysI            NaN     NaN            NaN                   NaN
CDS           NaN
1530         b2764  NADPH-dependent assimilatory sulfite reductase flavoprotein
subunit        cysJ            NaN     NaN            NaN                   NaN
CDS           NaN
2663         b0118    bifunctional aconitate hydratase 2/2-methylisocitrate
dehydratase       acnB            NaN     NaN            NaN
NaN          CDS           NaN
3524         b0750                                             quinolinate
synthase NadA       nadA            NaN     NaN            NaN
```

NaN CDS NaN
3813 b2763 assimilatory sulfite reductase (NADPH) hemoprotein
subunit cysI NaN NaN NaN NaN
CDS NaN
3814 b2764 NADPH-dependent assimilatory sulfite reductase flavoprotein
subunit cysJ NaN NaN NaN NaN
CDS NaN
4352 b0118 bifunctional aconitate hydratase 2/2-methylisocitrate
dehydratase acnB NaN NaN NaN
NaN CDS NaN
4353 b0118 bifunctional aconitate hydratase 2/2-methylisocitrate
dehydratase acnB NaN NaN NaN
NaN CDS NaN
4990 b0750 quinolinate
synthase NadA nadA NaN NaN NaN
NaN CDS NaN
4991 b0750 quinolinate
synthase NadA nadA NaN NaN NaN
NaN CDS NaN
7173 b2763 assimilatory sulfite reductase (NADPH) hemoprotein
subunit cysI NaN NaN NaN NaN
CDS NaN
7174 b2763 assimilatory sulfite reductase (NADPH) hemoprotein
subunit cysI NaN NaN NaN NaN
CDS NaN
7175 b2764 NADPH-dependent assimilatory sulfite reductase flavoprotein
subunit cysJ NaN NaN NaN NaN
CDS NaN
7176 b2764 NADPH-dependent assimilatory sulfite reductase flavoprotein
subunit cysJ NaN NaN NaN NaN
CDS NaN
8308 b3771 dihydroxy-acid
dehydratase ilvD NaN NaN NaN
NaN CDS NaN
8309 b3771 dihydroxy-acid
dehydratase ilvD NaN NaN NaN
NaN CDS NaN

Complex ID Cofactors in Modified Complex
Generic Complex ID MetaComplex ID ME-model SubReaction M-model Reaction ID \
355 CPLX0-7761:2 4fe4s(1)
NaN NaN NaN ACONTa
357 CPLX0-7761:2 4fe4s(1)
NaN NaN NaN ACONTb
1131 DIHYDROXYACIDDEHYDRAT-CPLX:2 4fe4s(1)
NaN NaN NaN DHAD1
1132 DIHYDROXYACIDDEHYDRAT-CPLX:2 4fe4s(1)

```
NaN            NaN                   NaN               DHAD2
1398          SULFITE-REDUCT-CPLX:4  4fe4s(1) AND sheme(1) AND fmn(1) AND fad(1)
NaN            NaN                   NaN              FADRx2
1399          SULFITE-REDUCT-CPLX:8  4fe4s(1) AND sheme(1) AND fmn(1) AND fad(1)
NaN            NaN                   NaN              FADRx2
1521          SULFITE-REDUCT-CPLX:4  4fe4s(1) AND sheme(1) AND fmn(1) AND fad(1)
NaN            NaN                   NaN                FLVR
1522          SULFITE-REDUCT-CPLX:8  4fe4s(1) AND sheme(1) AND fmn(1) AND fad(1)
NaN            NaN                   NaN                FLVR
1529          SULFITE-REDUCT-CPLX:4  4fe4s(1) AND sheme(1) AND fmn(1) AND fad(1)
NaN            NaN                   NaN              FMNRx2
1530          SULFITE-REDUCT-CPLX:8  4fe4s(1) AND sheme(1) AND fmn(1) AND fad(1)
NaN            NaN                   NaN              FMNRx2
2663                   CPLX0-7761:2                                    4fe4s(1)
NaN            NaN                   NaN             MICITDr
3524                   CPLX0-7719:2                                    4fe4s(1)
NaN            NaN                   NaN               QULNS
3813          SULFITE-REDUCT-CPLX:4  4fe4s(1) AND sheme(1) AND fmn(1) AND fad(1)
NaN            NaN                   NaN               SULRi
3814          SULFITE-REDUCT-CPLX:8  4fe4s(1) AND sheme(1) AND fmn(1) AND fad(1)
NaN            NaN                   NaN               SULRi
4352                   CPLX0-7761:2                                         NaN
NaN            NaN                   NaN                 NaN
4353                   CPLX0-7761:2                                    4fe4s(1)
NaN            NaN                   NaN                 NaN
4990                   CPLX0-7719:2                                         NaN
NaN            NaN                   NaN                 NaN
4991                   CPLX0-7719:2                                    4fe4s(1)
NaN            NaN                   NaN                 NaN
7173          SULFITE-REDUCT-CPLX:4                                         NaN
NaN            NaN                   NaN                 NaN
7174          SULFITE-REDUCT-CPLX:4  4fe4s(1) AND sheme(1) AND fmn(1) AND fad(1)
NaN            NaN                   NaN                 NaN
7175          SULFITE-REDUCT-CPLX:8                                         NaN
NaN            NaN                   NaN                 NaN
7176          SULFITE-REDUCT-CPLX:8  4fe4s(1) AND sheme(1) AND fmn(1) AND fad(1)
NaN            NaN                   NaN                 NaN
8308  DIHYDROXYACIDDEHYDRAT-CPLX:2                                          NaN
NaN            NaN                   NaN                 NaN
8309  DIHYDROXYACIDDEHYDRAT-CPLX:2                                     4fe4s(1)
NaN            NaN                   NaN                 NaN

                                                       Reaction Name
Reversibility  GroEL_dependent_folding  DnaK_dependent_folding
N_terminal_methionine_cleavage  RNA mods/enzyme  \
355                aconitase (half-reaction A, Citrate hydro-lyase)
True                       NaN                     NaN
```

NaN NaN
357 aconitase (half-reaction B, Isocitrate hydro-lyase)
True NaN NaN
NaN NaN
1131 dihydroxy-acid dehydratase (2,3-dihydroxy-3-methylbutanoate)
False NaN NaN
NaN NaN
1132 Dihydroxy-acid dehydratase (2,3-dihydroxy-3-methylpentanoate)
False NaN NaN
NaN NaN
1398 FAD reductase
False NaN NaN
NaN NaN
1399 FAD reductase
False NaN NaN
NaN NaN
1521 flavin reductase
False NaN NaN
NaN NaN
1522 flavin reductase
False NaN NaN
NaN NaN
1529 FMN reductase
False NaN NaN
NaN NaN
1530 FMN reductase
False NaN NaN
NaN NaN
2663 2-methylisocitrate dehydratase
True NaN NaN
NaN NaN
3524 quinolinate synthase
False NaN NaN
NaN NaN
3813 sulfite reductase (NADPH2)
False NaN NaN
NaN NaN
3814 sulfite reductase (NADPH2)
False NaN NaN
NaN NaN
4352 NaN
NaN NaN NaN
NaN NaN
4353 NaN
NaN NaN NaN
NaN NaN
4990 NaN

```
NaN                      NaN                      NaN
NaN                NaN
4991                                                             NaN
NaN                      NaN                      NaN
NaN                NaN
7173                                                             NaN
NaN                      NaN                      NaN
NaN                NaN
7174                                                             NaN
NaN                      NaN                      NaN
NaN                NaN
7175                                                             NaN
NaN                      NaN                      NaN
NaN                NaN
7176                                                             NaN
NaN                      NaN                      NaN
NaN                NaN
8308                                                             NaN
NaN                      NaN                      NaN
NaN                NaN
8309                                                             NaN
NaN                      NaN                      NaN
NaN                NaN

     tRNA-codon association  RNA stability  Complex Location  Subunit Location
Translocation Pathway
355                    None            NaN               NaN               NaN
NaN
357                    None            NaN               NaN               NaN
NaN
1131                   None            NaN               NaN               NaN
NaN
1132                   None            NaN               NaN               NaN
NaN
1398                   None            NaN               NaN               NaN
NaN
1399                   None            NaN               NaN               NaN
NaN
1521                   None            NaN               NaN               NaN
NaN
1522                   None            NaN               NaN               NaN
NaN
1529                   None            NaN               NaN               NaN
NaN
1530                   None            NaN               NaN               NaN
NaN
2663                   None            NaN               NaN               NaN
```

```
NaN
3524                    None          NaN              NaN              NaN
NaN
3813                    None          NaN              NaN              NaN
NaN
3814                    None          NaN              NaN              NaN
NaN
4352                    None          NaN              NaN              NaN
NaN
4353                    None          NaN              NaN              NaN
NaN
4990                    None          NaN              NaN              NaN
NaN
4991                    None          NaN              NaN              NaN
NaN
7173                    None          NaN              NaN              NaN
NaN
7174                    None          NaN              NaN              NaN
NaN
7175                    None          NaN              NaN              NaN
NaN
7176                    None          NaN              NaN              NaN
NaN
8308                    None          NaN              NaN              NaN
NaN
8309                    None          NaN              NaN              NaN
NaN
```

[13]:
```
# check problematic complexes
cplxs = 'CPLX0-7617|CPLX0-7824|IscA_tetra'
target[target['Complex ID'].str.contains(cplxs, na = False)]
```

[13]:
```
     Gene Locus ID                                 Definition Gene Names  Old
Locus Tag  BioCyc  Reference BBH  Reference Definition Feature Type  Complex
Name    Complex ID  \
4401          b0156  iron-sulfur cluster insertion protein ErpA       erpA
NaN     NaN             NaN                     NaN          CDS          NaN
CPLX0-7617:2
4402          b0156  iron-sulfur cluster insertion protein ErpA       erpA
NaN     NaN             NaN                     NaN          CDS          NaN
CPLX0-7617:2
4403          b0156  iron-sulfur cluster insertion protein ErpA       erpA
NaN     NaN             NaN                     NaN          CDS          NaN
CPLX0-7617:2
5973          b1684         Fe-S cluster assembly scaffold SufA       sufA
NaN     NaN             NaN                     NaN          CDS          NaN
CPLX0-7824:2
```

5974 b1684 Fe-S cluster assembly scaffold SufA sufA
NaN NaN NaN NaN CDS NaN
CPLX0-7824:2
5975 b1684 Fe-S cluster assembly scaffold SufA sufA
NaN NaN NaN NaN CDS NaN
CPLX0-7824:2
6917 b2528 iron-sulfur cluster assembly protein IscA iscA
NaN NaN NaN NaN CDS NaN
IscA_tetra:4
6918 b2528 iron-sulfur cluster assembly protein IscA iscA
NaN NaN NaN NaN CDS NaN
IscA_tetra:4
6919 b2528 iron-sulfur cluster assembly protein IscA iscA
NaN NaN NaN NaN CDS NaN
IscA_tetra:4

Cofactors in Modified Complex Generic Complex ID MetaComplex ID ME-model SubReaction M-model Reaction ID Reaction Name Reversibility
GroEL_dependent_folding DnaK_dependent_folding \
4401 NaN fes_transfers_complex NaN
NaN NaN NaN NaN NaN
NaN
4402 2fe2s(1) NaN NaN
NaN NaN NaN NaN NaN
NaN
4403 4fe4s(1) NaN NaN
NaN NaN NaN NaN NaN
NaN
5973 NaN fes_transfers_complex NaN
NaN NaN NaN NaN NaN
NaN
5974 2fe2s(1) NaN NaN
NaN NaN NaN NaN NaN
NaN
5975 4fe4s(1) NaN NaN
NaN NaN NaN NaN NaN
NaN
6917 NaN fes_transfers_complex NaN
NaN NaN NaN NaN NaN
NaN
6918 2fe2s(1) NaN NaN
NaN NaN NaN NaN NaN
NaN
6919 4fe4s(1) NaN NaN
NaN NaN NaN NaN NaN
NaN

| | N_terminal_methionine_cleavage | RNA mods/enzyme | tRNA-codon association | RNA stability | Complex Location | Subunit Location | Translocation | Pathway |
|---|---|---|---|---|---|---|---|---|
| 4401 | | NaN | NaN | None | NaN | NaN | NaN | NaN |
| 4402 | | NaN | NaN | None | NaN | NaN | NaN | NaN |
| 4403 | | NaN | NaN | None | NaN | NaN | NaN | NaN |
| 5973 | | NaN | NaN | None | NaN | NaN | NaN | NaN |
| 5974 | | NaN | NaN | None | NaN | NaN | NaN | NaN |
| 5975 | | NaN | NaN | None | NaN | NaN | NaN | NaN |
| 6917 | | NaN | NaN | None | NaN | NaN | NaN | NaN |
| 6918 | | NaN | NaN | None | NaN | NaN | NaN | NaN |
| 6919 | | NaN | NaN | None | NaN | NaN | NaN | NaN |
