## Supplementary File S3 for "Expanded ME-model analysis capabilities enable genetic recoding, proteostasis, RNA decay, and cofactor essentiality in bacterial models"

November 3, 2026

[1]:
```
import itables
itables.init_notebook_mode(all_interactive=True)
```

```
<IPython.core.display.HTML object>
```

```
<IPython.core.display.HTML object>
```

[2]:
```
import os
os.environ["TQDM_DISABLE"] = "1"
from tqdm import tqdm
```

[3]:
```
# Install dependencies
# !{sys.executable} -m pip install -r requirements.txt
```

[4]:
```
import sys
sys.path.insert(0, './Supplementary-File-S8')
import coralme
coralme.__file__.split('Supplementary-File-S8')[1]
```

[4]:
```
'/coralme/__init__.py'
```

Revert coralME genome file

[5]:
```
%%bash
for organism in rE.coli-57 C321.deltaA rEc.b.dC.12 rEc.y.dC.46 ALE-C321.deltaA␣
↪Ochre; do
    cp "blastn-and-blastp-bbh-outputs/ASM584_to_${organism}/
↪combined-${organism}-pEVOL-pBpF.gb" \
        "GRO-Ecoli-ME-${organism}-noBBH/building_data/genome_modified.gb"
done
```

[6]:
```
from coralme.builder.main import MEBuilder
def build_me_model(organism):
    builder = MEBuilder(config_files=['GRO-Ecoli-ME-{:s}-noBBH/coralme-config.
↪yaml'.format(organism)])
    builder.configuration['df_gene_cplxs_mods_rxns'] = 'GRO-Ecoli-ME-{:s}-noBBH/
↪automated-org-with-refs-tRNAs.xlsx'.format(organism)

    if organism in ['rEc.b.dC.12', 'rEc.y.dC.46']:
```

```
        builder.configuration['knockouts'] = [ 'b1211' ] # Delta PrfA (Release␣
↪Factor 1)
        builder.configuration['genetic_recoding'] = { 'TAG' : { 'bipa__L_c' :␣
↪'trna-bipA' }} # 4-Biphenylyl-L-alanine

    builder.configuration['run_bbh_blast'] = False
    builder.configuration['include_pseudo_genes'] = True # needed for internal␣
↪codon stops in rEc genomes
    builder.configuration['add_rrna_modifications'] = True
    builder.configuration['feature_types'] = ['CDS', 'rRNA', 'tRNA', 'ncRNA']

    if organism == 'rEc.b.dC.12':
        builder.configuration['knockouts'] += [ 'pgaptmp_001771',␣
↪'pgaptmp_003770' ]
    elif organism == 'rEc.y.dC.46':
        builder.configuration['knockouts'] += [ 'pgaptmp_001771',␣
↪'pgaptmp_003768' ]

    try:
        builder.build_me_model(overwrite = False)
    except Exception:
        import traceback
        traceback.print_exc()
    return builder.me_model
```

[7]:

```
import multiprocessing
with multiprocessing.Pool(multiprocessing.cpu_count()-1) as pool:
    res = pool.map(build_me_model, [ 'ASM584v2', 'rE.coli-57', 'C321.deltaA',␣
↪'ALE-C321.deltaA', 'rEc.b.dC.12', 'rEc.y.dC.46', 'Ochre' ])
```

```
Initiating ME-model reconstruction…
Initiating ME-model reconstruction…
Initiating ME-model reconstruction…
Initiating ME-model reconstruction…
Initiating ME-model reconstruction…
Initiating ME-model reconstruction…
Initiating ME-model reconstruction…
ME-model was saved in the GRO-Ecoli-ME-Ochre-noBBH directory as MEModel-
step1-GRO-Ecoli-ME-Ochre-noBBH.pkl
ME-model was saved in the GRO-Ecoli-ME-rE.coli-57-noBBH directory as MEModel-
step1-GRO-Ecoli-ME-rE.coli-57-noBBH.pkl
ME-model was saved in the GRO-Ecoli-ME-ALE-C321.deltaA-noBBH directory as
MEModel-step1-GRO-Ecoli-ME-ALE-C321.deltaA-noBBH.pkl
ME-model was saved in the GRO-Ecoli-ME-ASM584v2-noBBH directory as MEModel-
step1-GRO-Ecoli-ME-ASM584v2-noBBH.pkl
ME-model was saved in the GRO-Ecoli-ME-C321.deltaA-noBBH directory as MEModel-
step1-GRO-Ecoli-ME-C321.deltaA-noBBH.pkl
```

```
ME-model was saved in the GRO-Ecoli-ME-rEc.y.dC.46-noBBH directory as MEModel-
step1-GRO-Ecoli-ME-rEc.y.dC.46-noBBH.pkl
ME-model was saved in the GRO-Ecoli-ME-rEc.b.dC.12-noBBH directory as MEModel-
step1-GRO-Ecoli-ME-rEc.b.dC.12-noBBH.pkl
ME-model was saved in the GRO-Ecoli-ME-rE.coli-57-noBBH directory as MEModel-
step2-GRO-Ecoli-ME-rE.coli-57-noBBH.pkl
ME-model reconstruction is done.
Number of metabolites in the ME-model is 6080 (+266.27%, from 1660)
Number of reactions in the ME-model is 13245 (+457.21%, from 2377)
Number of genes in the ME-model is 1505 (+19.07%, from 1264)
Number of missing genes from reconstruction cannot be determined.
ME-model was saved in the GRO-Ecoli-ME-Ochre-noBBH directory as MEModel-
step2-GRO-Ecoli-ME-Ochre-noBBH.pkl
ME-model reconstruction is done.
Number of metabolites in the ME-model is 6161 (+270.48%, from 1663)
Number of reactions in the ME-model is 13849 (+481.16%, from 2383)
Number of genes in the ME-model is 1528 (+20.89%, from 1264)
Number of missing genes from reconstruction cannot be determined.
ME-model was saved in the GRO-Ecoli-ME-ALE-C321.deltaA-noBBH directory as
MEModel-step2-GRO-Ecoli-ME-ALE-C321.deltaA-noBBH.pkl
ME-model reconstruction is done.
Number of metabolites in the ME-model is 6245 (+276.20%, from 1660)
Number of reactions in the ME-model is 14037 (+490.53%, from 2377)
Number of genes in the ME-model is 1566 (+23.89%, from 1264)
Number of missing genes from reconstruction cannot be determined.
ME-model was saved in the GRO-Ecoli-ME-C321.deltaA-noBBH directory as MEModel-
step2-GRO-Ecoli-ME-C321.deltaA-noBBH.pkl
ME-model reconstruction is done.
Number of metabolites in the ME-model is 6216 (+274.46%, from 1660)
Number of reactions in the ME-model is 13992 (+488.64%, from 2377)
Number of genes in the ME-model is 1555 (+23.02%, from 1264)
Number of missing genes from reconstruction cannot be determined.
ME-model was saved in the GRO-Ecoli-ME-rEc.y.dC.46-noBBH directory as MEModel-
step2-GRO-Ecoli-ME-rEc.y.dC.46-noBBH.pkl
ME-model reconstruction is done.
Number of metabolites in the ME-model is 6255 (+276.13%, from 1663)
Number of reactions in the ME-model is 14067 (+490.31%, from 2383)
Number of genes in the ME-model is 1569 (+24.13%, from 1264)
Number of missing genes from reconstruction cannot be determined.
ME-model was saved in the GRO-Ecoli-ME-rEc.b.dC.12-noBBH directory as MEModel-
step2-GRO-Ecoli-ME-rEc.b.dC.12-noBBH.pkl
ME-model reconstruction is done.
Number of metabolites in the ME-model is 6261 (+276.49%, from 1663)
Number of reactions in the ME-model is 14073 (+490.56%, from 2383)
Number of genes in the ME-model is 1571 (+24.29%, from 1264)
Number of missing genes from reconstruction cannot be determined.
ME-model was saved in the GRO-Ecoli-ME-ASM584v2-noBBH directory as MEModel-
step2-GRO-Ecoli-ME-ASM584v2-noBBH.pkl
```

```
ME-model reconstruction is done.
Number of metabolites in the ME-model is 6989 (+321.02%, from 1660)
Number of reactions in the ME-model is 14974 (+529.95%, from 2377)
Number of genes in the ME-model is 1663 (+31.57%, from 1264)
Number of missing genes from reconstruction cannot be determined.
```

[8]:
```
model = res[-2]
print(model.pseudo_genes)

for data in model.translation_data:
    if data.notes:
        print(data.id, data.notes)
```

```
['RNA_b0090', 'RNA_b0893', 'RNA_pgaptmp_003451']
b0090 ['Codon at position 242 encodes a recoded stop codon.']
b0893 ['Codon at position 212 encodes a recoded stop codon.']
b1474 ['Codon at position 195 encodes Selenocysteine.']
pgaptmp_003451 ["Last codon 'UGG' (position 155) does not encode a stop codon."]
b3894 ['Codon at position 195 encodes Selenocysteine.']
b4079 ['Codon at position 139 encodes Selenocysteine.']
```

[9]:
```
model
```

[9]:
```
<MEModel GRO-Ecoli-ME-rEc.y.dC.46-noBBH at 0x700862e3f7d0>
```

[10]:
```
model.query('charging_tRNA_trna-bipA_UAG')[0].reaction
```

[10]:
```
'[0.000116266666666667*mu + 4.55184e-5] RNA_trna-bipA +
[4.27350427350427e-6*mu*(0.000116266666666667*mu + 1.0000455184)]
bipA_tRNA_synth_copy1-MONOMER + [0.000116266666666667*mu + 4.55184e-5] bipa__L_c
--> 1.0 generic_tRNA_UAG_bipa__L_c'
```

[11]:
```
model.query('translation_b0893')[0]
```

[11]:
```
<TranslationReaction translation_b0893 at 0x7006f2833080>
```

[12]:
```
model.reactions.EX_btn_e.bounds = (-1000., +1000.)
model.remove_troubleshooted_reactions()
import coralme
coralme.builder.main.METroubleshooter.troubleshoot(model, check = ['E-matrix'])
```

```
WARNING:root:Metabolite types valid values are ME-Deadends, Biomass, Cofactors,
Amino-acids, All-Deadends, Metabolite, GenericComponent, ProcessedProtein,
TranslatedGene, TranscribedGene, Complex. The predefined order of metabolites
will be tested.


The MINOS and quad MINOS solvers are a courtesy of Prof Michael A. Saunders.
Please cite Ma, D., Yang, L., Fleming, R. et al. Reliable and efficient solution
of genome-scale models of Metabolism and macromolecular Expression. Sci Rep 7,
```

```
40863 (2017). https://doi.org/10.1038/srep40863

~ Troubleshooting started…
  Checking if the coralME model can simulate growth without gapfilling
reactions…
  Original coralME model is feasible with a tested growth rate of 0.001000 1/h
~ Final step. Fully optimizing with precision 1e-6 and save solution into the
coralME model…
  Gapfilled coralME model is feasible with growth rate 0.899447 (M-model:
0.000000).
coralME model was saved in the ./ directory as MEModel-step3-GRO-Ecoli-ME-
rEc.y.dC.46-noBBH-TS.pkl
```

[13]:
```python
model.query('b1637')
```

```
/opt/python3-venv/p312-test_coralME/lib/python3.12/site-
packages/IPython/core/formatters.py:370: FormatterWarning: text/html formatter
returned invalid type <class 'coralme.core.extended_classes.MappableList'>
(expected <class 'str'>) for object: MappableList([<TranscribedGene RNA_b1637 at
0x7006e17b18b0>, <TranslatedGene protein_b1637 at 0x7006e17b2d80>,
<TranslationReaction translation_b1637 at 0x7006e17b1460>,
<TranscriptionReaction transcription_TU_b1637 at 0x7006e17b34d0>, <MEReaction
DM_RNA_b1637 at 0x7006cfc23500>, <TranslationData b1637 at 0x7006e17b2f00>,
<TranscriptionData TU_b1637 at 0x7006e17b3e90>])
  warnings.warn(
```

[13]:
```
MappableList([<TranscribedGene RNA_b1637 at 0x7006e17b18b0>,
 <TranslatedGene protein_b1637 at 0x7006e17b2d80>,
 <TranslationReaction translation_b1637 at 0x7006e17b1460>,
 <TranscriptionReaction transcription_TU_b1637 at 0x7006e17b34d0>,
 <MEReaction DM_RNA_b1637 at 0x7006cfc23500>,
 <TranslationData b1637 at 0x7006e17b2f00>,
 <TranscriptionData TU_b1637 at 0x7006e17b3e90>])
```

[14]:
```python
model.query('charging_tRNA_trna-bipA_UAG')[0].metabolites
```

[14]:
```
{<GenerictRNA generic_tRNA_UAG_bipa__L_c at 0x7006f66337a0>: 1.0,
 <TranscribedGene RNA_trna-bipA at 0x700700c04bc0>: -0.000116266666666667*mu -
4.55184e-5,
 <Metabolite bipa__L_c at 0x7006f6630bc0>: -0.000116266666666667*mu -
4.55184e-5,
 <Complex bipA_tRNA_synth_copy1-MONOMER at 0x7006cfe77740>:
-4.27350427350427e-6*mu*(0.000116266666666667*mu + 1.0000455184)}
```

[15]:
```python
model.global_info['knockouts']
```

[15]:
```
['b1211', 'pgaptmp_001771', 'pgaptmp_003768']
```

[16]:
```
model.query('Tyr_RS_dim')[0].reactions
```

[16]:
```
frozenset({<tRNAChargingReaction charging_tRNA_pgaptmp_001290_UAC at
0x7006cfead880>,
           <tRNAChargingReaction charging_tRNA_pgaptmp_001290_UAU at
0x7006cfead550>,
           <tRNAChargingReaction charging_tRNA_pgaptmp_001292_UAC at
0x7006cfeadd90>,
           <tRNAChargingReaction charging_tRNA_pgaptmp_001292_UAU at
0x7006cfeada90>,
           <tRNAChargingReaction charging_tRNA_pgaptmp_004106_UAC at
0x7006cfe38470>,
           <tRNAChargingReaction charging_tRNA_pgaptmp_004106_UAU at
0x7006cfe38170>,
           <ComplexFormation formation_Tyr_RS_dim at 0x7006cf50f4a0>})
```

[17]:
```
model.feasibility(0.001)
```

[17]:
```
True
```

[18]:
```
model.query('charging_tRNA_trna-bipA_UAG')[0].flux
```

[18]: $3.933755134114987 \times 10^{-12} \ \frac{\text{millimole}}{(\text{gram}\cdot\text{hour})}$

[19]:
```
model.feasibility(0.0001)
```

[19]:
```
True
```

[20]:
```
model.query('charging_tRNA_trna-bipA_UAG')[0].flux
```

[20]: $3.987050740258725 \times 10^{-14} \ \frac{\text{millimole}}{(\text{gram}\cdot\text{hour})}$

[21]:
```
model.doubling_time
```

[21]: 6931.471805599453 hour
