## Supplementary File S4 for "Expanded ME-model analysis capabilities enable genetic recoding, proteostasis, RNA decay, and cofactor essentiality in bacterial models"

November 3, 2026

[1]:
```
import itables
itables.init_notebook_mode(all_interactive=True)
```

```
<IPython.core.display.HTML object>
```

```
<IPython.core.display.HTML object>
```

[2]:
```
import os
os.environ["TQDM_DISABLE"] = "1"
from tqdm import tqdm
```

[3]:
```
# Install dependencies
# !{sys.executable} -m pip install -r requirements.txt
```

[4]:
```
import sys
sys.path.insert(0, './Supplementary-File-S8')
import coralme
coralme.__file__.split('Supplementary-File-S8')[1]
```

[4]:
```
'/coralme/__init__.py'
```

[5]:
```
models = [
    'GRO-Ecoli-ME-ASM584v2-noBBH',
    'GRO-Ecoli-ME-rE.coli-57-noBBH',
    'GRO-Ecoli-ME-Ochre-noBBH',
    'GRO-Ecoli-ME-C321.deltaA-noBBH',
    'GRO-Ecoli-ME-ALE-C321.deltaA-noBBH',
    'GRO-Ecoli-ME-rEc.b.dC.12-noBBH',
    'GRO-Ecoli-ME-rEc.y.dC.46-noBBH',
]
models
```

[5]:
```
['GRO-Ecoli-ME-ASM584v2-noBBH',
 'GRO-Ecoli-ME-rE.coli-57-noBBH',
 'GRO-Ecoli-ME-Ochre-noBBH',
 'GRO-Ecoli-ME-C321.deltaA-noBBH',
 'GRO-Ecoli-ME-ALE-C321.deltaA-noBBH',
 'GRO-Ecoli-ME-rEc.b.dC.12-noBBH',
 'GRO-Ecoli-ME-rEc.y.dC.46-noBBH']
```

[6]:
```
import coralme
for org in models:
    me = coralme.io.pickle.load_pickle_me_model(f'{org}/MEModel-step2-{org}.
 ↪pkl')
    print(len(me.reactions))
    print(len(me.metabolites))
    print(len(me.all_genes))
    print()
```

```
14974
6989
1663

13245
6080
1505

13849
6161
1528

13992
6216
1555

14037
6245
1566

14073
6261
1571

14067
6255
1569
```

[7]:
```
import coralme
for org in models:
    me = coralme.io.pickle.load_pickle_me_model(f'{org}/MEModel-step2-{org}.
 ↪pkl')
    me.reactions.EX_btn_e.bounds = (-1000., +1000.)
    me._all_genes = [] # reset
    print(me.pseudo_genes)

    for data in me.translation_data:
```

```python
        if data.notes:
            print(data.id, data.notes)

    # print(me.reactions.formation_ribosome.reaction)
    me.optimize()
    print(me.growth_rate)
    print()
```

```
[]
b1474 ['Codon at position 195 encodes Selenocysteine.']
b3894 ['Codon at position 195 encodes Selenocysteine.']
b4079 ['Codon at position 139 encodes Selenocysteine.']
The MINOS and quad MINOS solvers are a courtesy of Prof Michael A. Saunders.
Please cite Ma, D., Yang, L., Fleming, R. et al. Reliable and efficient solution
of genome-scale models of Metabolism and macromolecular Expression. Sci Rep 7,
40863 (2017). https://doi.org/10.1038/srep40863

Iteration        Solution to check        Solver Status
---------        ------------------       -------------
        1        1.4050280687025918       Not feasible
        2        0.7025140343512959       Optimal
        3        1.0537710515269438       Not feasible
        4        0.8781425429391199       Not feasible
        5        0.7903282886452079       Not feasible
        6        0.7464211614982519       Optimal
        7        0.7683747250717299       Optimal
        8        0.7793515068584689       Not feasible
        9        0.7738631159650994       Not feasible
       10        0.7711189205184146       Not feasible
       11        0.7697468227950722       Not feasible
       12        0.7690607739334010       Not feasible
       13        0.7687177495025654       Optimal
       14        0.7688892617179832       Not feasible
       15        0.7688035056102743       Optimal
       16        0.7688463836641288       Optimal
       17        0.7688678226910560       Optimal
       18        0.7688785422045197       Not feasible
       19        0.7688731824477879       Optimal
       20        0.7688758623261538       Not feasible
       21        0.7688745223869708       Not feasible
       22        0.7688738524173794       Optimal
0.7688738524173794 / hour

[]
b1474 ['Codon at position 195 encodes Selenocysteine.']
b3894 ['Codon at position 195 encodes Selenocysteine.']
b4079 ['Codon at position 139 encodes Selenocysteine.']
The MINOS and quad MINOS solvers are a courtesy of Prof Michael A. Saunders.
```

```
Please cite Ma, D., Yang, L., Fleming, R. et al. Reliable and efficient solution
of genome-scale models of Metabolism and macromolecular Expression. Sci Rep 7,
40863 (2017). https://doi.org/10.1038/srep40863

Iteration       Solution to check       Solver Status
---------       ------------------      -------------
        1       1.4050280687025918      Not feasible
        2       0.7025140343512959      Optimal
        3       1.0537710515269438      Not feasible
        4       0.8781425429391199      Not feasible
        5       0.7903282886452079      Optimal
        6       0.8342354157921639      Not feasible
        7       0.8122818522186859      Not feasible
        8       0.8013050704319469      Not feasible
        9       0.7958166795385774      Optimal
       10       0.7985608749852622      Optimal
       11       0.7999329727086045      Optimal
       12       0.8006190215702758      Optimal
       13       0.8009620460011113      Not feasible
       14       0.8007905337856935      Not feasible
       15       0.8007047776779846      Not feasible
       16       0.8006618996241301      Optimal
       17       0.8006833386510575      Not feasible
       18       0.8006726191375938      Optimal
       19       0.8006779788943257      Not feasible
       20       0.8006752990159598      Not feasible
       21       0.8006739590767769      Optimal
       22       0.8006746290463683      Optimal
0.8006746290463683 / hour

[]
b1474 ['Codon at position 195 encodes Selenocysteine.']
b4079 ['Codon at position 139 encodes Selenocysteine.']
b3894 ['Codon at position 195 encodes Selenocysteine.']
The MINOS and quad MINOS solvers are a courtesy of Prof Michael A. Saunders.
Please cite Ma, D., Yang, L., Fleming, R. et al. Reliable and efficient solution
of genome-scale models of Metabolism and macromolecular Expression. Sci Rep 7,
40863 (2017). https://doi.org/10.1038/srep40863

Iteration       Solution to check       Solver Status
---------       ------------------      -------------
        1       1.4050280687025918      Not feasible
        2       0.7025140343512959      Optimal
        3       1.0537710515269438      Not feasible
        4       0.8781425429391199      Optimal
        5       0.9659567972330318      Not feasible
        6       0.9220496700860759      Not feasible
        7       0.9000961065125979      Optimal
```

```
        8       0.9110728882993369      Not feasible
        9       0.9055844974059674      Not feasible
       10       0.9028403019592826      Not feasible
       11       0.9014682042359402      Optimal
       12       0.9021542530976114      Not feasible
       13       0.9018112286667759      Not feasible
       14       0.9016397164513581      Not feasible
       15       0.9015539603436491      Optimal
       16       0.9015968383975036      Optimal
       17       0.9016182774244308      Optimal
       18       0.9016289969378944      Not feasible
       19       0.9016236371811626      Not feasible
       20       0.9016209573027967      Not feasible
       21       0.9016196173636137      Optimal
       22       0.9016202873332052      Not feasible
0.9016196173636137 / hour

[]
The MINOS and quad MINOS solvers are a courtesy of Prof Michael A. Saunders.
Please cite Ma, D., Yang, L., Fleming, R. et al. Reliable and efficient solution
of genome-scale models of Metabolism and macromolecular Expression. Sci Rep 7,
40863 (2017). https://doi.org/10.1038/srep40863

Iteration       Solution to check       Solver Status
---------       ------------------      -------------
        1       1.4050280687025918      Not feasible
        2       0.7025140343512959      Optimal
        3       1.0537710515269438      Not feasible
        4       0.8781425429391199      Not feasible
        5       0.7903282886452079      Optimal
        6       0.8342354157921639      Not feasible
        7       0.8122818522186859      Optimal
        8       0.8232586340054249      Optimal
        9       0.8287470248987944      Optimal
       10       0.8314912203454792      Optimal
       11       0.8328633180688215      Not feasible
       12       0.8321772692071503      Not feasible
       13       0.8318342447763147      Optimal
       14       0.8320057569917325      Optimal
       15       0.8320915130994414      Optimal
       16       0.8321343911532959      Not feasible
       17       0.8321129521263686      Optimal
       18       0.8321236716398323      Not feasible
       19       0.8321183118831004      Not feasible
       20       0.8321156320047345      Not feasible
       21       0.8321142920655515      Optimal
       22       0.8321149620351430      Optimal
0.832114962035143 / hour
```

```
[]
b1474 ['Codon at position 195 encodes Selenocysteine.']
b3894 ['Codon at position 195 encodes Selenocysteine.']
b4079 ['Codon at position 139 encodes Selenocysteine.']
The MINOS and quad MINOS solvers are a courtesy of Prof Michael A. Saunders.
Please cite Ma, D., Yang, L., Fleming, R. et al. Reliable and efficient solution
of genome-scale models of Metabolism and macromolecular Expression. Sci Rep 7,
40863 (2017). https://doi.org/10.1038/srep40863

Iteration       Solution to check       Solver Status
---------       ------------------      -------------
        1       1.4050280687025918      Not feasible
        2       0.7025140343512959      Optimal
        3       1.0537710515269438      Not feasible
        4       0.8781425429391199      Not feasible
        5       0.7903282886452079      Optimal
        6       0.8342354157921639      Not feasible
        7       0.8122818522186859      Optimal
        8       0.8232586340054249      Not feasible
        9       0.8177702431120554      Not feasible
       10       0.8150260476653706      Not feasible
       11       0.8136539499420282      Not feasible
       12       0.8129679010803570      Not feasible
       13       0.8126248766495214      Optimal
       14       0.8127963888649392      Optimal
       15       0.8128821449726481      Not feasible
       16       0.8128392669187936      Not feasible
       17       0.8128178278918664      Not feasible
       18       0.8128071083784028      Optimal
       19       0.8128124681351345      Not feasible
       20       0.8128097882567686      Optimal
       21       0.8128111281959516      Not feasible
       22       0.8128104582263601      Optimal
0.8128104582263601 / hour

['RNA_b3035', 'RNA_pgaptmp_003453']
b1474 ['Codon at position 195 encodes Selenocysteine.']
b3035 ['Codon at position 81 encodes a recoded stop codon.']
pgaptmp_003453 ["Last codon 'UGG' (position 155) does not encode a stop codon."]
b3894 ['Codon at position 195 encodes Selenocysteine.']
b4079 ['Codon at position 139 encodes Selenocysteine.']
The MINOS and quad MINOS solvers are a courtesy of Prof Michael A. Saunders.
Please cite Ma, D., Yang, L., Fleming, R. et al. Reliable and efficient solution
of genome-scale models of Metabolism and macromolecular Expression. Sci Rep 7,
40863 (2017). https://doi.org/10.1038/srep40863

Iteration       Solution to check       Solver Status
```

```
---------       ------------------      -------------
        1       1.4050280687025918      Not feasible
        2       0.7025140343512959      Optimal
        3       1.0537710515269438      Not feasible
        4       0.8781425429391199      Optimal
        5       0.9659567972330318      Not feasible
        6       0.9220496700860759      Not feasible
        7       0.9000961065125979      Not feasible
        8       0.8891193247258589      Optimal
        9       0.8946077156192284      Optimal
       10       0.8973519110659132      Optimal
       11       0.8987240087892555      Optimal
       12       0.8994100576509267      Optimal
       13       0.8997530820817623      Optimal
       14       0.8999245942971801      Optimal
       15       0.9000103504048890      Not feasible
       16       0.8999674723510345      Optimal
       17       0.8999889113779618      Optimal
       18       0.8999996308914253      Optimal
       19       0.9000049906481571      Not feasible
       20       0.9000023107697912      Optimal
       21       0.9000036507089741      Not feasible
       22       0.9000029807393827      Not feasible
0.9000023107697912 / hour

['RNA_b0090', 'RNA_b0893', 'RNA_pgaptmp_003451']
b0090 ['Codon at position 242 encodes a recoded stop codon.']
b0893 ['Codon at position 212 encodes a recoded stop codon.']
b1474 ['Codon at position 195 encodes Selenocysteine.']
pgaptmp_003451 ["Last codon 'UGG' (position 155) does not encode a stop codon."]
b3894 ['Codon at position 195 encodes Selenocysteine.']
b4079 ['Codon at position 139 encodes Selenocysteine.']
The MINOS and quad MINOS solvers are a courtesy of Prof Michael A. Saunders.
Please cite Ma, D., Yang, L., Fleming, R. et al. Reliable and efficient solution
of genome-scale models of Metabolism and macromolecular Expression. Sci Rep 7,
40863 (2017). https://doi.org/10.1038/srep40863

Iteration       Solution to check       Solver Status
---------       ------------------      -------------
        1       1.4050280687025918      Not feasible
        2       0.7025140343512959      Optimal
        3       1.0537710515269438      Not feasible
        4       0.8781425429391199      Optimal
        5       0.9659567972330318      Not feasible
        6       0.9220496700860759      Not feasible
        7       0.9000961065125979      Not feasible
        8       0.8891193247258589      Optimal
        9       0.8946077156192284      Optimal
```

```
        10      0.8973519110659132      Optimal
        11      0.8987240087892555      Optimal
        12      0.8994100576509267      Optimal
        13      0.8997530820817623      Not feasible
        14      0.8995815698663445      Not feasible
        15      0.8994958137586356      Not feasible
        16      0.8994529357047811      Not feasible
        17      0.8994314966778539      Optimal
        18      0.8994422161913176      Optimal
        19      0.8994475759480494      Not feasible
        20      0.8994448960696835      Optimal
        21      0.899446236008866       Optimal
        22      0.8994469059784579      Optimal
0.8994469059784579 / hour
```

[8]:

```
import coralme
for org in [
    'GRO-Ecoli-ME-rEc.b.dC.12-noBBH',
    'GRO-Ecoli-ME-rEc.y.dC.46-noBBH']:

    me = coralme.io.pickle.load_pickle_me_model(f'{org}/MEModel-step2-{org}.
 ↪pkl')
    print(len(me.all_genes))
    me.reactions.EX_btn_e.bounds = (-1000., +1000.)
    me._all_genes = [] # reset
    print(me.pseudo_genes)

    for data in me.translation_data:
        if data.notes:
            print(data.id, data.notes)

    # print(me.reactions.formation_ribosome.reaction)
    me.reactions.EX_bipa__L_e.bounds = (0., 0.)
    display(me.reactions.EX_bipa__L_e)
    print(me.feasibility())
    print()
```

```
1571
['RNA_b3035', 'RNA_pgaptmp_003453']
b1474 ['Codon at position 195 encodes Selenocysteine.']
b3035 ['Codon at position 81 encodes a recoded stop codon.']
pgaptmp_003453 ["Last codon 'UGG' (position 155) does not encode a stop codon."]
b3894 ['Codon at position 195 encodes Selenocysteine.']
b4079 ['Codon at position 139 encodes Selenocysteine.']

<BoundaryReaction EX_bipa__L_e at 0x786ac5f4a180>

True
```

```
1569
['RNA_b0090', 'RNA_b0893', 'RNA_pgaptmp_003451']
b0090 ['Codon at position 242 encodes a recoded stop codon.']
b0893 ['Codon at position 212 encodes a recoded stop codon.']
b1474 ['Codon at position 195 encodes Selenocysteine.']
pgaptmp_003451 ["Last codon 'UGG' (position 155) does not encode a stop codon."]
b3894 ['Codon at position 195 encodes Selenocysteine.']
b4079 ['Codon at position 139 encodes Selenocysteine.']
```

```
<BoundaryReaction EX_bipa__L_e at 0x786ac61a8050>
```

```
False
```

[9]:

```
import coralme
for org in models:
    me = coralme.io.pickle.load_pickle_me_model(f'{org}/MEModel-step2-{org}.
 ↪pkl')
    me.reactions.EX_btn_e.bounds = (-1000., +1000.)
    me._all_genes = [] # reset

    ref = pandas.read_csv('genes-reference.txt', header = None, index_col = 0).
 ↪index
    lst = set(ref).difference([ x.id[4:] for x in me.all_genes ])
    print('Missing genes from ref:', len(lst), sorted(lst))
    lst = set([ x.id[4:] for x in me.all_genes ]).difference(ref)
    print('Missing genes in ref:', len(lst), sorted(lst))
    print()
    # 'b2610' in set(ref).difference([ x.id[4:] for x in me.all_genes ]),\ # ffh
    # 'b3123' in set(ref).difference([ x.id[4:] for x in me.all_genes ]),\ #␣
 ↪rnpB ncRNA
    # 'b0455' in set(ref).difference([ x.id[4:] for x in me.all_genes ]) # ffs␣
 ↪ncRNA
```

```
Missing genes from ref: 45 ['b0188', 'b0195', 'b0405', 'b0406', 'b0636',
'b0661', 'b0969', 'b1133', 'b1344', 'b1807', 'b1870', 'b1871', 'b1922', 'b2092',
'b2318', 'b2324', 'b2474', 'b2532', 'b2559', 'b2573', 'b2575', 'b2583', 'b2607',
'b2741', 'b2745', 'b2791', 'b2960', 'b3064', 'b3166', 'b3202', 'b3282', 'b3343',
'b3344', 'b3345', 'b3461', 'b3470', 'b3651', 'b3706', 'b3741', 'b3783', 'b3965',
'b4166', 'b4168', 'b4171', 'b4293']
Missing genes in ref: 19 ['b0296', 'b0437', 'b0438', 'b0439', 'b0835', 'b0852',
'b0882', 'b0905', 'b1128', 'b2330', 'b3178', 'b3259', 'b3492', 'b3499', 'b3931',
'b3932', 'b4174', 'b4175', 'b4506']

Missing genes from ref: 307 ['b0014', 'b0015', 'b0019', 'b0027', 'b0051',
'b0053', 'b0058', 'b0098', 'b0150', 'b0151', 'b0152', 'b0153', 'b0168', 'b0170',
'b0172', 'b0177', 'b0188', 'b0195', 'b0201', 'b0202', 'b0203', 'b0204', 'b0205',
'b0206', 'b0216', 'b0244', 'b0260', 'b0261', 'b0273', 'b0306', 'b0307', 'b0308',
```

'b0365', 'b0366', 'b0367', 'b0368', 'b0405', 'b0406', 'b0408', 'b0409', 'b0455',
'b0536', 'b0578', 'b0627', 'b0636', 'b0657', 'b0661', 'b0664', 'b0665', 'b0666',
'b0668', 'b0670', 'b0672', 'b0673', 'b0743', 'b0744', 'b0745', 'b0746', 'b0747',
'b0748', 'b0749', 'b0807', 'b0859', 'b0883', 'b0884', 'b0891', 'b0948', 'b0967',
'b0969', 'b0971', 'b1032', 'b1086', 'b1116', 'b1117', 'b1118', 'b1133', 'b1135',
'b1209', 'b1230', 'b1231', 'b1269', 'b1325', 'b1326', 'b1329', 'b1344', 'b1363',
'b1377', 'b1378', 'b1380', 'b1385', 'b1386', 'b1387', 'b1388', 'b1389', 'b1390',
'b1391', 'b1392', 'b1393', 'b1394', 'b1395', 'b1397', 'b1398', 'b1539', 'b1665',
'b1666', 'b1718', 'b1801', 'b1807', 'b1822', 'b1835', 'b1870', 'b1871', 'b1909',
'b1910', 'b1911', 'b1919', 'b1922', 'b1927', 'b1975', 'b1977', 'b1984', 'b1986',
'b1989', 'b2032', 'b2033', 'b2034', 'b2035', 'b2036', 'b2037', 'b2038', 'b2039',
'b2040', 'b2041', 'b2045', 'b2048', 'b2049', 'b2051', 'b2052', 'b2053', 'b2091',
'b2092', 'b2093', 'b2094', 'b2095', 'b2096', 'b2183', 'b2189', 'b2318', 'b2324',
'b2348', 'b2392', 'b2393', 'b2396', 'b2397', 'b2401', 'b2402', 'b2403', 'b2404',
'b2474', 'b2477', 'b2512', 'b2517', 'b2532', 'b2559', 'b2573', 'b2575', 'b2583',
'b2588', 'b2589', 'b2590', 'b2591', 'b2594', 'b2595', 'b2607', 'b2610', 'b2614',
'b2617', 'b2652', 'b2691', 'b2692', 'b2693', 'b2694', 'b2695', 'b2741', 'b2745',
'b2785', 'b2791', 'b2806', 'b2814', 'b2815', 'b2816', 'b2828', 'b2864', 'b2946',
'b2960', 'b2967', 'b2975', 'b2976', 'b2979', 'b2987', 'b3064', 'b3069', 'b3084',
'b3123', 'b3166', 'b3168', 'b3171', 'b3174', 'b3175', 'b3179', 'b3202', 'b3272',
'b3273', 'b3274', 'b3275', 'b3276', 'b3277', 'b3278', 'b3282', 'b3287', 'b3288',
'b3289', 'b3300', 'b3340', 'b3341', 'b3343', 'b3344', 'b3345', 'b3461', 'b3464',
'b3465', 'b3470', 'b3497', 'b3545', 'b3590', 'b3591', 'b3609', 'b3626', 'b3643',
'b3651', 'b3658', 'b3705', 'b3706', 'b3740', 'b3741', 'b3756', 'b3757', 'b3758',
'b3759', 'b3760', 'b3761', 'b3783', 'b3796', 'b3797', 'b3798', 'b3799', 'b3836',
'b3838', 'b3839', 'b3851', 'b3852', 'b3853', 'b3854', 'b3855', 'b3965', 'b3968',
'b3969', 'b3970', 'b3971', 'b3976', 'b3977', 'b3978', 'b3979', 'b3981', 'b4007',
'b4008', 'b4009', 'b4010', 'b4022', 'b4134', 'b4142', 'b4143', 'b4163', 'b4164',
'b4165', 'b4166', 'b4168', 'b4171', 'b4180', 'b4270', 'b4287', 'b4288', 'b4289',
'b4290', 'b4291', 'b4293', 'b4301', 'b4311', 'b4356', 'b4358', 'b4368', 'b4369',
'b4370', 'b4371', 'b4375', 'b4467', 'b4468']
Missing genes in ref: 123 ['CmR', 'b0437', 'b0438', 'b0439', 'b0835', 'b0852',
'b0882', 'b0905', 'b1066', 'b1128', 'b2330', 'b3178', 'b3259', 'b3931', 'b3932',
'b4174', 'b4175', 'pgaptmp_000193', 'pgaptmp_000194', 'pgaptmp_000195',
'pgaptmp_000196', 'pgaptmp_000197', 'pgaptmp_000198', 'pgaptmp_000208',
'pgaptmp_000238', 'pgaptmp_000439', 'pgaptmp_000532', 'pgaptmp_000533',
'pgaptmp_000534', 'pgaptmp_000535', 'pgaptmp_000536', 'pgaptmp_000537',
'pgaptmp_000538', 'pgaptmp_000605', 'pgaptmp_000606', 'pgaptmp_000607',
'pgaptmp_000608', 'pgaptmp_000609', 'pgaptmp_000610', 'pgaptmp_000611',
'pgaptmp_000753', 'pgaptmp_000841', 'pgaptmp_000886', 'pgaptmp_001041',
'pgaptmp_001043', 'pgaptmp_001360', 'pgaptmp_001361', 'pgaptmp_001596',
'pgaptmp_001597', 'pgaptmp_001598', 'pgaptmp_001627', 'pgaptmp_001629',
'pgaptmp_001635', 'pgaptmp_001637', 'pgaptmp_001640', 'pgaptmp_001781',
'pgaptmp_001940', 'pgaptmp_001968', 'pgaptmp_001969', 'pgaptmp_001973',
'pgaptmp_001974', 'pgaptmp_001975', 'pgaptmp_001976', 'pgaptmp_002160',
'pgaptmp_002161', 'pgaptmp_002162', 'pgaptmp_002163', 'pgaptmp_002221',
'pgaptmp_002222', 'pgaptmp_002223', 'pgaptmp_002224', 'pgaptmp_002225',
'pgaptmp_002345', 'pgaptmp_002346', 'pgaptmp_002347', 'pgaptmp_002389',

```
'pgaptmp_002494', 'pgaptmp_002569', 'pgaptmp_002670', 'pgaptmp_002673',
'pgaptmp_002764', 'pgaptmp_002765', 'pgaptmp_002766', 'pgaptmp_002767',
'pgaptmp_002768', 'pgaptmp_002769', 'pgaptmp_002770', 'pgaptmp_003015',
'pgaptmp_003231', 'pgaptmp_003232', 'pgaptmp_003233', 'pgaptmp_003234',
'pgaptmp_003235', 'pgaptmp_003236', 'pgaptmp_003270', 'pgaptmp_003271',
'pgaptmp_003272', 'pgaptmp_003273', 'pgaptmp_003325', 'pgaptmp_003326',
'pgaptmp_003327', 'pgaptmp_003328', 'pgaptmp_003329', 'pgaptmp_003442',
'pgaptmp_003443', 'pgaptmp_003444', 'pgaptmp_003445', 'pgaptmp_003449',
'pgaptmp_003450', 'pgaptmp_003451', 'pgaptmp_003452', 'pgaptmp_003481',
'pgaptmp_003482', 'pgaptmp_003483', 'pgaptmp_003484', 'pgaptmp_003612',
'pgaptmp_003643', 'pgaptmp_003644', 'pgaptmp_003645', 'pgaptmp_003749',
'pgaptmp_003764', 'pgaptmp_003765', 'pgaptmp_003766']

Missing genes from ref: 281 ['b0007', 'b0014', 'b0015', 'b0019', 'b0027',
'b0051', 'b0053', 'b0058', 'b0098', 'b0168', 'b0170', 'b0172', 'b0177', 'b0188',
'b0195', 'b0201', 'b0202', 'b0203', 'b0204', 'b0205', 'b0206', 'b0216', 'b0244',
'b0260', 'b0261', 'b0273', 'b0349', 'b0405', 'b0406', 'b0408', 'b0409', 'b0452',
'b0455', 'b0536', 'b0613', 'b0627', 'b0636', 'b0657', 'b0661', 'b0664', 'b0665',
'b0666', 'b0668', 'b0670', 'b0672', 'b0673', 'b0743', 'b0744', 'b0745', 'b0746',
'b0747', 'b0748', 'b0749', 'b0807', 'b0829', 'b0857', 'b0859', 'b0883', 'b0884',
'b0891', 'b0948', 'b0967', 'b0969', 'b0971', 'b1032', 'b1086', 'b1116', 'b1117',
'b1118', 'b1133', 'b1135', 'b1193', 'b1200', 'b1209', 'b1211', 'b1223', 'b1230',
'b1231', 'b1269', 'b1325', 'b1344', 'b1363', 'b1385', 'b1444', 'b1484', 'b1588',
'b1602', 'b1611', 'b1621', 'b1665', 'b1666', 'b1680', 'b1683', 'b1718', 'b1801',
'b1807', 'b1822', 'b1835', 'b1870', 'b1871', 'b1909', 'b1910', 'b1911', 'b1922',
'b1975', 'b1977', 'b1984', 'b1986', 'b1989', 'b1993', 'b2048', 'b2092', 'b2132',
'b2149', 'b2169', 'b2183', 'b2189', 'b2202', 'b2224', 'b2258', 'b2310', 'b2318',
'b2324', 'b2341', 'b2348', 'b2396', 'b2397', 'b2401', 'b2402', 'b2403', 'b2404',
'b2474', 'b2477', 'b2480', 'b2482', 'b2492', 'b2512', 'b2517', 'b2532', 'b2552',
'b2559', 'b2573', 'b2575', 'b2583', 'b2588', 'b2589', 'b2590', 'b2591', 'b2594',
'b2595', 'b2607', 'b2610', 'b2614', 'b2617', 'b2652', 'b2691', 'b2692', 'b2693',
'b2694', 'b2695', 'b2703', 'b2741', 'b2745', 'b2785', 'b2791', 'b2796', 'b2806',
'b2814', 'b2815', 'b2816', 'b2828', 'b2864', 'b2866', 'b2919', 'b2946', 'b2960',
'b2967', 'b3064', 'b3069', 'b3084', 'b3089', 'b3123', 'b3166', 'b3168', 'b3171',
'b3174', 'b3175', 'b3179', 'b3182', 'b3202', 'b3272', 'b3273', 'b3274', 'b3275',
'b3276', 'b3277', 'b3278', 'b3282', 'b3287', 'b3288', 'b3289', 'b3300', 'b3340',
'b3341', 'b3343', 'b3344', 'b3345', 'b3350', 'b3461', 'b3464', 'b3465', 'b3470',
'b3497', 'b3545', 'b3546', 'b3590', 'b3591', 'b3609', 'b3623', 'b3643', 'b3651',
'b3658', 'b3705', 'b3706', 'b3740', 'b3741', 'b3756', 'b3757', 'b3758', 'b3759',
'b3760', 'b3761', 'b3783', 'b3785', 'b3796', 'b3797', 'b3798', 'b3799', 'b3836',
'b3838', 'b3839', 'b3851', 'b3852', 'b3853', 'b3854', 'b3855', 'b3965', 'b3968',
'b3969', 'b3970', 'b3971', 'b3976', 'b3977', 'b3978', 'b3979', 'b3981', 'b4007',
'b4008', 'b4009', 'b4010', 'b4022', 'b4134', 'b4142', 'b4143', 'b4163', 'b4164',
'b4165', 'b4166', 'b4168', 'b4171', 'b4180', 'b4219', 'b4270', 'b4287', 'b4293',
'b4301', 'b4368', 'b4369', 'b4370', 'b4371', 'b4375']
Missing genes in ref: 120 ['ACI6BN_00005', 'ACI6BN_00010', 'ACI6BN_00015',
'ACI6BN_00020', 'ACI6BN_00955', 'ACI6BN_00960', 'ACI6BN_00965', 'ACI6BN_00970',
'ACI6BN_00990', 'ACI6BN_00995', 'ACI6BN_01160', 'ACI6BN_01955', 'ACI6BN_02990',
```

'ACI6BN_03005', 'ACI6BN_03015', 'ACI6BN_03045', 'ACI6BN_03055', 'ACI6BN_03355', 'ACI6BN_03360', 'ACI6BN_03365', 'ACI6BN_04610', 'ACI6BN_04615', 'ACI6BN_06655', 'ACI6BN_06665', 'ACI6BN_07595', 'ACI6BN_07900', 'ACI6BN_08350', 'ACI6BN_09045', 'ACI6BN_09050', 'ACI6BN_09055', 'ACI6BN_09060', 'ACI6BN_09065', 'ACI6BN_09395', 'ACI6BN_09400', 'ACI6BN_09405', 'ACI6BN_09410', 'ACI6BN_09415', 'ACI6BN_09420', 'ACI6BN_09425', 'ACI6BN_10020', 'ACI6BN_11320', 'ACI6BN_11470', 'ACI6BN_11520', 'ACI6BN_11525', 'ACI6BN_11530', 'ACI6BN_11535', 'ACI6BN_11540', 'ACI6BN_11545', 'ACI6BN_12700', 'ACI6BN_13155', 'ACI6BN_13670', 'ACI6BN_13675', 'ACI6BN_13680', 'ACI6BN_13835', 'ACI6BN_14485', 'ACI6BN_14490', 'ACI6BN_14495', 'ACI6BN_14500', 'ACI6BN_14645', 'ACI6BN_14650', 'ACI6BN_14655', 'ACI6BN_14660', 'ACI6BN_14680', 'ACI6BN_14685', 'ACI6BN_14690', 'ACI6BN_14695', 'ACI6BN_15265', 'ACI6BN_15270', 'ACI6BN_15275', 'ACI6BN_15280', 'ACI6BN_15285', 'ACI6BN_15540', 'ACI6BN_15545', 'ACI6BN_15550', 'ACI6BN_15555', 'ACI6BN_15725', 'ACI6BN_15730', 'ACI6BN_15735', 'ACI6BN_15740', 'ACI6BN_15745', 'ACI6BN_15750', 'ACI6BN_16880', 'ACI6BN_18230', 'ACI6BN_18235', 'ACI6BN_18240', 'ACI6BN_18245', 'ACI6BN_18250', 'ACI6BN_18255', 'ACI6BN_18260', 'ACI6BN_18720', 'ACI6BN_18735', 'ACI6BN_19250', 'ACI6BN_19745', 'ACI6BN_20275', 'ACI6BN_20510', 'ACI6BN_21110', 'ACI6BN_21115', 'ACI6BN_21120', 'ACI6BN_21125', 'ACI6BN_21130', 'ACI6BN_21295', 'CmR', 'b0296', 'b0437', 'b0438', 'b0439', 'b0835', 'b0852', 'b0882', 'b0905', 'b1066', 'b1128', 'b2330', 'b3178', 'b3259', 'b3931', 'b3932', 'b4174', 'b4175', 'b4506']

Missing genes from ref: 256 ['b0007', 'b0014', 'b0015', 'b0027', 'b0051', 'b0053', 'b0058', 'b0098', 'b0168', 'b0170', 'b0172', 'b0177', 'b0188', 'b0195', 'b0201', 'b0202', 'b0203', 'b0204', 'b0205', 'b0206', 'b0216', 'b0244', 'b0344', 'b0349', 'b0405', 'b0406', 'b0408', 'b0409', 'b0455', 'b0536', 'b0627', 'b0636', 'b0657', 'b0661', 'b0664', 'b0665', 'b0666', 'b0668', 'b0670', 'b0672', 'b0673', 'b0743', 'b0744', 'b0745', 'b0746', 'b0747', 'b0748', 'b0749', 'b0756', 'b0757', 'b0774', 'b0775', 'b0807', 'b0859', 'b0883', 'b0884', 'b0891', 'b0948', 'b0967', 'b0969', 'b0971', 'b1032', 'b1086', 'b1116', 'b1117', 'b1118', 'b1133', 'b1135', 'b1200', 'b1209', 'b1211', 'b1230', 'b1231', 'b1269', 'b1325', 'b1344', 'b1474', 'b1611', 'b1621', 'b1665', 'b1666', 'b1680', 'b1683', 'b1718', 'b1807', 'b1822', 'b1835', 'b1870', 'b1871', 'b1909', 'b1910', 'b1911', 'b1922', 'b1975', 'b1977', 'b1984', 'b1986', 'b1989', 'b2092', 'b2183', 'b2189', 'b2255', 'b2256', 'b2318', 'b2324', 'b2348', 'b2396', 'b2397', 'b2401', 'b2402', 'b2403', 'b2404', 'b2474', 'b2477', 'b2480', 'b2492', 'b2512', 'b2517', 'b2532', 'b2559', 'b2573', 'b2575', 'b2583', 'b2588', 'b2589', 'b2590', 'b2591', 'b2594', 'b2595', 'b2607', 'b2610', 'b2614', 'b2617', 'b2652', 'b2691', 'b2692', 'b2693', 'b2694', 'b2695', 'b2701', 'b2741', 'b2745', 'b2785', 'b2791', 'b2806', 'b2814', 'b2815', 'b2816', 'b2828', 'b2864', 'b2946', 'b2960', 'b2967', 'b3035', 'b3064', 'b3069', 'b3084', 'b3089', 'b3123', 'b3166', 'b3168', 'b3171', 'b3174', 'b3175', 'b3179', 'b3182', 'b3202', 'b3272', 'b3273', 'b3274', 'b3275', 'b3276', 'b3277', 'b3278', 'b3282', 'b3287', 'b3288', 'b3289', 'b3300', 'b3340', 'b3343', 'b3344', 'b3345', 'b3460', 'b3461', 'b3464', 'b3465', 'b3470', 'b3497', 'b3545', 'b3546', 'b3564', 'b3590', 'b3591', 'b3609', 'b3651', 'b3658', 'b3705', 'b3706', 'b3740', 'b3741', 'b3756', 'b3757', 'b3758', 'b3759', 'b3760', 'b3761', 'b3783', 'b3785', 'b3796', 'b3797', 'b3798', 'b3799', 'b3836', 'b3838', 'b3839', 'b3851', 'b3852', 'b3853', 'b3854', 'b3855', 'b3894', 'b3965', 'b3968', 'b3969', 'b3970', 'b3971', 'b3976', 'b3977', 'b3978', 'b3979', 'b3981', 'b4007', 'b4008', 'b4009', 'b4010', 'b4022', 'b4079', 'b4134',

'b4142', 'b4143', 'b4163', 'b4164', 'b4165', 'b4166', 'b4168', 'b4171', 'b4180', 'b4270', 'b4287', 'b4293', 'b4368', 'b4369', 'b4370', 'b4371', 'b4375']
Missing genes in ref: 122 ['CmR', 'N840_0199', 'N840_0200', 'N840_0201', 'N840_0202', 'N840_0203', 'N840_0204', 'N840_0214', 'N840_0244', 'N840_0541', 'N840_0677', 'N840_0678', 'N840_0679', 'N840_0680', 'N840_0681', 'N840_0682', 'N840_0683', 'N840_0756', 'N840_0757', 'N840_0758', 'N840_0759', 'N840_0760', 'N840_0761', 'N840_0762', 'N840_0892', 'N840_0980', 'N840_1041', 'N840_1247', 'N840_1248', 'N840_1699', 'N840_1700', 'N840_1949', 'N840_1950', 'N840_1951', 'N840_2012', 'N840_2014', 'N840_2020', 'N840_2022', 'N840_2025', 'N840_2236', 'N840_2396', 'N840_2447', 'N840_2448', 'N840_2452', 'N840_2453', 'N840_2454', 'N840_2455', 'N840_2647', 'N840_2648', 'N840_2649', 'N840_2650', 'N840_2709', 'N840_2743', 'N840_2744', 'N840_2745', 'N840_2746', 'N840_2747', 'N840_2869', 'N840_2870', 'N840_2871', 'N840_2919', 'N840_3022', 'N840_3122', 'N840_3223', 'N840_3226', 'N840_3325', 'N840_3326', 'N840_3327', 'N840_3328', 'N840_3329', 'N840_3330', 'N840_3331', 'N840_3603', 'N840_3829', 'N840_3830', 'N840_3831', 'N840_3832', 'N840_3833', 'N840_3834', 'N840_3867', 'N840_3868', 'N840_3869', 'N840_3870', 'N840_3921', 'N840_3922', 'N840_3923', 'N840_3924', 'N840_3925', 'N840_4039', 'N840_4040', 'N840_4041', 'N840_4042', 'N840_4046', 'N840_4047', 'N840_4048', 'N840_4049', 'N840_4079', 'N840_4080', 'N840_4081', 'N840_4082', 'N840_4212', 'N840_4243', 'N840_4244', 'N840_4245', 'N840_4348', 'N840_4451', 'N840_4452', 'N840_4453', 'b0296', 'b0437', 'b0438', 'b0439', 'b0835', 'b0852', 'b0882', 'b0905', 'b3178', 'b3259', 'b3931', 'b3932', 'b4174', 'b4175']

Missing genes from ref: 249 ['b0007', 'b0014', 'b0015', 'b0027', 'b0051', 'b0053', 'b0058', 'b0098', 'b0168', 'b0170', 'b0172', 'b0177', 'b0188', 'b0195', 'b0201', 'b0202', 'b0203', 'b0204', 'b0205', 'b0206', 'b0216', 'b0244', 'b0349', 'b0405', 'b0406', 'b0408', 'b0409', 'b0455', 'b0536', 'b0627', 'b0636', 'b0657', 'b0661', 'b0664', 'b0665', 'b0666', 'b0668', 'b0670', 'b0672', 'b0673', 'b0743', 'b0744', 'b0745', 'b0746', 'b0747', 'b0748', 'b0749', 'b0774', 'b0775', 'b0807', 'b0859', 'b0883', 'b0884', 'b0891', 'b0948', 'b0967', 'b0969', 'b0971', 'b1032', 'b1086', 'b1116', 'b1117', 'b1118', 'b1133', 'b1135', 'b1200', 'b1209', 'b1211', 'b1230', 'b1231', 'b1269', 'b1325', 'b1344', 'b1600', 'b1611', 'b1621', 'b1665', 'b1666', 'b1680', 'b1683', 'b1718', 'b1801', 'b1807', 'b1822', 'b1835', 'b1849', 'b1870', 'b1871', 'b1909', 'b1910', 'b1911', 'b1922', 'b1975', 'b1977', 'b1984', 'b1986', 'b1989', 'b2092', 'b2183', 'b2189', 'b2318', 'b2324', 'b2348', 'b2396', 'b2397', 'b2401', 'b2402', 'b2403', 'b2404', 'b2474', 'b2477', 'b2480', 'b2512', 'b2517', 'b2532', 'b2559', 'b2573', 'b2575', 'b2583', 'b2588', 'b2589', 'b2590', 'b2591', 'b2594', 'b2595', 'b2607', 'b2610', 'b2614', 'b2617', 'b2652', 'b2691', 'b2692', 'b2693', 'b2694', 'b2695', 'b2741', 'b2745', 'b2785', 'b2791', 'b2806', 'b2814', 'b2815', 'b2816', 'b2828', 'b2864', 'b2946', 'b2960', 'b2967', 'b3035', 'b3064', 'b3069', 'b3084', 'b3123', 'b3166', 'b3168', 'b3171', 'b3174', 'b3175', 'b3179', 'b3182', 'b3202', 'b3272', 'b3273', 'b3274', 'b3275', 'b3276', 'b3277', 'b3278', 'b3282', 'b3287', 'b3288', 'b3289', 'b3300', 'b3340', 'b3341', 'b3343', 'b3344', 'b3345', 'b3461', 'b3464', 'b3465', 'b3470', 'b3497', 'b3545', 'b3546', 'b3564', 'b3590', 'b3591', 'b3609', 'b3643', 'b3651', 'b3658', 'b3705', 'b3706', 'b3740', 'b3741', 'b3756', 'b3757', 'b3758', 'b3759', 'b3760', 'b3761', 'b3783', 'b3785', 'b3796', 'b3797', 'b3798', 'b3799', 'b3836', 'b3838', 'b3839', 'b3851', 'b3852', 'b3853', 'b3854', 'b3855', 'b3965', 'b3968', 'b3969', 'b3970', 'b3971',

```
'b3976', 'b3977', 'b3978', 'b3979', 'b3981', 'b4007', 'b4008', 'b4009', 'b4010',
'b4022', 'b4134', 'b4142', 'b4143', 'b4163', 'b4164', 'b4165', 'b4166', 'b4168',
'b4171', 'b4180', 'b4270', 'b4287', 'b4293', 'b4368', 'b4369', 'b4370', 'b4371',
'b4375']
Missing genes in ref: 126 ['CXP41_RS01050', 'CXP41_RS01055', 'CXP41_RS01060',
'CXP41_RS01065', 'CXP41_RS01070', 'CXP41_RS01075', 'CXP41_RS01130',
'CXP41_RS01280', 'CXP41_RS02915', 'CXP41_RS03665', 'CXP41_RS03670',
'CXP41_RS03675', 'CXP41_RS03680', 'CXP41_RS03685', 'CXP41_RS03690',
'CXP41_RS03695', 'CXP41_RS04085', 'CXP41_RS04090', 'CXP41_RS04095',
'CXP41_RS04100', 'CXP41_RS04105', 'CXP41_RS04110', 'CXP41_RS04115',
'CXP41_RS04925', 'CXP41_RS05385', 'CXP41_RS05730', 'CXP41_RS06845',
'CXP41_RS06855', 'CXP41_RS09265', 'CXP41_RS09270', 'CXP41_RS10570',
'CXP41_RS10575', 'CXP41_RS10580', 'CXP41_RS10910', 'CXP41_RS10920',
'CXP41_RS10960', 'CXP41_RS10970', 'CXP41_RS10985', 'CXP41_RS12120',
'CXP41_RS12980', 'CXP41_RS13230', 'CXP41_RS13235', 'CXP41_RS13255',
'CXP41_RS13260', 'CXP41_RS13265', 'CXP41_RS13270', 'CXP41_RS14280',
'CXP41_RS14285', 'CXP41_RS14290', 'CXP41_RS14295', 'CXP41_RS14600',
'CXP41_RS14780', 'CXP41_RS14785', 'CXP41_RS14790', 'CXP41_RS14795',
'CXP41_RS14800', 'CXP41_RS15410', 'CXP41_RS15415', 'CXP41_RS15420',
'CXP41_RS15665', 'CXP41_RS16215', 'CXP41_RS16775', 'CXP41_RS17310',
'CXP41_RS17325', 'CXP41_RS17865', 'CXP41_RS17870', 'CXP41_RS17875',
'CXP41_RS17880', 'CXP41_RS17885', 'CXP41_RS17890', 'CXP41_RS17895',
'CXP41_RS19350', 'CXP41_RS20515', 'CXP41_RS20520', 'CXP41_RS20525',
'CXP41_RS20530', 'CXP41_RS20535', 'CXP41_RS20540', 'CXP41_RS20725',
'CXP41_RS20730', 'CXP41_RS20735', 'CXP41_RS20740', 'CXP41_RS20995',
'CXP41_RS21000', 'CXP41_RS21005', 'CXP41_RS21010', 'CXP41_RS21015',
'CXP41_RS21600', 'CXP41_RS21605', 'CXP41_RS21610', 'CXP41_RS21615',
'CXP41_RS21645', 'CXP41_RS21650', 'CXP41_RS21655', 'CXP41_RS21660',
'CXP41_RS21820', 'CXP41_RS21825', 'CXP41_RS21830', 'CXP41_RS21835',
'CXP41_RS22555', 'CXP41_RS22710', 'CXP41_RS22715', 'CXP41_RS22720',
'CXP41_RS23270', 'CXP41_RS23865', 'CXP41_RS23870', 'CXP41_RS23875', 'CmR',
'b0296', 'b0437', 'b0438', 'b0439', 'b0835', 'b0852', 'b0882', 'b0905', 'b1066',
'b1128', 'b2330', 'b3178', 'b3259', 'b3931', 'b3932', 'b4174', 'b4175', 'b4506']

Missing genes from ref: 248 ['b0007', 'b0014', 'b0015', 'b0027', 'b0051',
'b0053', 'b0058', 'b0098', 'b0168', 'b0170', 'b0172', 'b0177', 'b0188', 'b0195',
'b0201', 'b0202', 'b0203', 'b0204', 'b0205', 'b0206', 'b0216', 'b0244', 'b0344',
'b0349', 'b0405', 'b0406', 'b0408', 'b0409', 'b0455', 'b0536', 'b0627', 'b0636',
'b0657', 'b0661', 'b0664', 'b0665', 'b0666', 'b0668', 'b0670', 'b0672', 'b0673',
'b0743', 'b0744', 'b0745', 'b0746', 'b0747', 'b0748', 'b0749', 'b0774', 'b0775',
'b0807', 'b0859', 'b0883', 'b0884', 'b0891', 'b0948', 'b0967', 'b0969', 'b0971',
'b1032', 'b1086', 'b1116', 'b1117', 'b1118', 'b1133', 'b1135', 'b1200', 'b1209',
'b1211', 'b1230', 'b1231', 'b1269', 'b1325', 'b1344', 'b1611', 'b1621', 'b1665',
'b1666', 'b1680', 'b1683', 'b1718', 'b1801', 'b1807', 'b1822', 'b1835', 'b1870',
'b1871', 'b1909', 'b1910', 'b1911', 'b1922', 'b1975', 'b1977', 'b1984', 'b1986',
'b1989', 'b2092', 'b2183', 'b2189', 'b2318', 'b2324', 'b2348', 'b2396', 'b2397',
'b2401', 'b2402', 'b2403', 'b2404', 'b2474', 'b2477', 'b2480', 'b2492', 'b2512',
'b2517', 'b2532', 'b2559', 'b2573', 'b2575', 'b2583', 'b2588', 'b2589', 'b2590',
```

'b2591', 'b2594', 'b2595', 'b2607', 'b2610', 'b2614', 'b2617', 'b2652', 'b2691', 'b2692', 'b2693', 'b2694', 'b2695', 'b2741', 'b2745', 'b2785', 'b2791', 'b2806', 'b2814', 'b2815', 'b2816', 'b2828', 'b2864', 'b2946', 'b2960', 'b2967', 'b3064', 'b3069', 'b3084', 'b3089', 'b3123', 'b3166', 'b3168', 'b3171', 'b3174', 'b3175', 'b3179', 'b3182', 'b3202', 'b3272', 'b3273', 'b3274', 'b3275', 'b3276', 'b3277', 'b3278', 'b3282', 'b3287', 'b3288', 'b3289', 'b3300', 'b3340', 'b3341', 'b3343', 'b3344', 'b3345', 'b3461', 'b3464', 'b3465', 'b3470', 'b3497', 'b3545', 'b3546', 'b3590', 'b3591', 'b3609', 'b3643', 'b3651', 'b3658', 'b3705', 'b3706', 'b3740', 'b3741', 'b3756', 'b3757', 'b3758', 'b3759', 'b3760', 'b3761', 'b3783', 'b3785', 'b3796', 'b3797', 'b3798', 'b3799', 'b3836', 'b3838', 'b3839', 'b3851', 'b3852', 'b3853', 'b3854', 'b3855', 'b3965', 'b3968', 'b3969', 'b3970', 'b3971', 'b3976', 'b3977', 'b3978', 'b3979', 'b3981', 'b4007', 'b4008', 'b4009', 'b4010', 'b4022', 'b4134', 'b4142', 'b4143', 'b4163', 'b4164', 'b4165', 'b4166', 'b4168', 'b4171', 'b4180', 'b4270', 'b4287', 'b4293', 'b4368', 'b4369', 'b4370', 'b4371', 'b4375']
Missing genes in ref: 130 ['CmR', 'b0296', 'b0437', 'b0438', 'b0439', 'b0835', 'b0852', 'b0882', 'b0905', 'b1066', 'b1128', 'b2330', 'b3178', 'b3259', 'b3931', 'b3932', 'b4174', 'b4175', 'b4506', 'bipA_tRNA_synth_copy1', 'pgaptmp_000204', 'pgaptmp_000205', 'pgaptmp_000206', 'pgaptmp_000207', 'pgaptmp_000208', 'pgaptmp_000209', 'pgaptmp_000219', 'pgaptmp_000249', 'pgaptmp_000550', 'pgaptmp_000685', 'pgaptmp_000686', 'pgaptmp_000687', 'pgaptmp_000688', 'pgaptmp_000689', 'pgaptmp_000690', 'pgaptmp_000691', 'pgaptmp_000766', 'pgaptmp_000767', 'pgaptmp_000768', 'pgaptmp_000769', 'pgaptmp_000770', 'pgaptmp_000771', 'pgaptmp_000772', 'pgaptmp_000932', 'pgaptmp_001020', 'pgaptmp_001085', 'pgaptmp_001291', 'pgaptmp_001293', 'pgaptmp_001755', 'pgaptmp_001756', 'pgaptmp_002005', 'pgaptmp_002006', 'pgaptmp_002007', 'pgaptmp_002067', 'pgaptmp_002069', 'pgaptmp_002075', 'pgaptmp_002077', 'pgaptmp_002080', 'pgaptmp_002288', 'pgaptmp_002417', 'pgaptmp_002452', 'pgaptmp_002502', 'pgaptmp_002503', 'pgaptmp_002507', 'pgaptmp_002508', 'pgaptmp_002509', 'pgaptmp_002510', 'pgaptmp_002705', 'pgaptmp_002706', 'pgaptmp_002707', 'pgaptmp_002708', 'pgaptmp_002766', 'pgaptmp_002799', 'pgaptmp_002800', 'pgaptmp_002801', 'pgaptmp_002802', 'pgaptmp_002803', 'pgaptmp_002925', 'pgaptmp_002926', 'pgaptmp_002927', 'pgaptmp_002976', 'pgaptmp_003083', 'pgaptmp_003182', 'pgaptmp_003283', 'pgaptmp_003286', 'pgaptmp_003384', 'pgaptmp_003385', 'pgaptmp_003386', 'pgaptmp_003387', 'pgaptmp_003388', 'pgaptmp_003389', 'pgaptmp_003390', 'pgaptmp_003453', 'pgaptmp_003664', 'pgaptmp_003888', 'pgaptmp_003889', 'pgaptmp_003890', 'pgaptmp_003891', 'pgaptmp_003892', 'pgaptmp_003893', 'pgaptmp_003927', 'pgaptmp_003928', 'pgaptmp_003929', 'pgaptmp_003930', 'pgaptmp_003982', 'pgaptmp_003983', 'pgaptmp_003984', 'pgaptmp_003985', 'pgaptmp_003986', 'pgaptmp_004100', 'pgaptmp_004101', 'pgaptmp_004102', 'pgaptmp_004103', 'pgaptmp_004107', 'pgaptmp_004108', 'pgaptmp_004109', 'pgaptmp_004110', 'pgaptmp_004139', 'pgaptmp_004140', 'pgaptmp_004141', 'pgaptmp_004142', 'pgaptmp_004270', 'pgaptmp_004301', 'pgaptmp_004302', 'pgaptmp_004303', 'pgaptmp_004407', 'pgaptmp_004514', 'pgaptmp_004515', 'pgaptmp_004516', 'trna-bipA']

Missing genes from ref: 250 ['b0007', 'b0014', 'b0015', 'b0027', 'b0051', 'b0053', 'b0058', 'b0098', 'b0168', 'b0170', 'b0172', 'b0177', 'b0188', 'b0195',

'b0201', 'b0202', 'b0203', 'b0204', 'b0205', 'b0206', 'b0216', 'b0244', 'b0344', 'b0349', 'b0405', 'b0406', 'b0408', 'b0409', 'b0455', 'b0536', 'b0627', 'b0636', 'b0657', 'b0661', 'b0664', 'b0665', 'b0666', 'b0668', 'b0670', 'b0672', 'b0673', 'b0743', 'b0744', 'b0745', 'b0746', 'b0747', 'b0748', 'b0749', 'b0774', 'b0775', 'b0807', 'b0859', 'b0883', 'b0884', 'b0891', 'b0948', 'b0967', 'b0969', 'b0971', 'b1032', 'b1086', 'b1116', 'b1117', 'b1118', 'b1133', 'b1135', 'b1200', 'b1209', 'b1211', 'b1230', 'b1231', 'b1269', 'b1325', 'b1344', 'b1385', 'b1611', 'b1621', 'b1665', 'b1666', 'b1680', 'b1683', 'b1718', 'b1801', 'b1807', 'b1822', 'b1835', 'b1870', 'b1871', 'b1909', 'b1910', 'b1911', 'b1922', 'b1975', 'b1977', 'b1984', 'b1986', 'b1989', 'b2092', 'b2183', 'b2189', 'b2318', 'b2324', 'b2348', 'b2396', 'b2397', 'b2401', 'b2402', 'b2403', 'b2404', 'b2474', 'b2477', 'b2480', 'b2512', 'b2517', 'b2532', 'b2559', 'b2573', 'b2575', 'b2583', 'b2588', 'b2589', 'b2590', 'b2591', 'b2594', 'b2595', 'b2607', 'b2610', 'b2614', 'b2617', 'b2652', 'b2691', 'b2692', 'b2693', 'b2694', 'b2695', 'b2741', 'b2745', 'b2785', 'b2791', 'b2806', 'b2814', 'b2815', 'b2816', 'b2828', 'b2864', 'b2866', 'b2946', 'b2960', 'b2967', 'b3064', 'b3069', 'b3084', 'b3089', 'b3123', 'b3166', 'b3168', 'b3171', 'b3174', 'b3175', 'b3179', 'b3182', 'b3202', 'b3272', 'b3273', 'b3274', 'b3275', 'b3276', 'b3277', 'b3278', 'b3282', 'b3287', 'b3288', 'b3289', 'b3300', 'b3340', 'b3341', 'b3343', 'b3344', 'b3345', 'b3461', 'b3464', 'b3465', 'b3470', 'b3497', 'b3545', 'b3546', 'b3564', 'b3590', 'b3591', 'b3609', 'b3643', 'b3651', 'b3658', 'b3705', 'b3706', 'b3740', 'b3741', 'b3756', 'b3757', 'b3758', 'b3759', 'b3760', 'b3761', 'b3783', 'b3785', 'b3796', 'b3797', 'b3798', 'b3799', 'b3836', 'b3838', 'b3839', 'b3851', 'b3852', 'b3853', 'b3854', 'b3855', 'b3965', 'b3968', 'b3969', 'b3970', 'b3971', 'b3976', 'b3977', 'b3978', 'b3979', 'b3981', 'b4007', 'b4008', 'b4009', 'b4010', 'b4022', 'b4134', 'b4142', 'b4143', 'b4163', 'b4164', 'b4165', 'b4166', 'b4168', 'b4171', 'b4180', 'b4270', 'b4287', 'b4293', 'b4368', 'b4369', 'b4370', 'b4371', 'b4375']
Missing genes in ref: 130 ['CmR', 'b0296', 'b0437', 'b0438', 'b0439', 'b0835', 'b0852', 'b0882', 'b0905', 'b1066', 'b1128', 'b2330', 'b3178', 'b3259', 'b3931', 'b3932', 'b4174', 'b4175', 'b4506', 'bipA_tRNA_synth_copy1', 'pgaptmp_000204', 'pgaptmp_000205', 'pgaptmp_000206', 'pgaptmp_000207', 'pgaptmp_000208', 'pgaptmp_000209', 'pgaptmp_000219', 'pgaptmp_000249', 'pgaptmp_000550', 'pgaptmp_000685', 'pgaptmp_000686', 'pgaptmp_000687', 'pgaptmp_000688', 'pgaptmp_000689', 'pgaptmp_000690', 'pgaptmp_000691', 'pgaptmp_000766', 'pgaptmp_000767', 'pgaptmp_000768', 'pgaptmp_000769', 'pgaptmp_000770', 'pgaptmp_000771', 'pgaptmp_000772', 'pgaptmp_000932', 'pgaptmp_001020', 'pgaptmp_001084', 'pgaptmp_001290', 'pgaptmp_001292', 'pgaptmp_001755', 'pgaptmp_001756', 'pgaptmp_002005', 'pgaptmp_002006', 'pgaptmp_002007', 'pgaptmp_002067', 'pgaptmp_002069', 'pgaptmp_002075', 'pgaptmp_002077', 'pgaptmp_002080', 'pgaptmp_002287', 'pgaptmp_002416', 'pgaptmp_002451', 'pgaptmp_002501', 'pgaptmp_002502', 'pgaptmp_002506', 'pgaptmp_002507', 'pgaptmp_002508', 'pgaptmp_002509', 'pgaptmp_002704', 'pgaptmp_002705', 'pgaptmp_002706', 'pgaptmp_002707', 'pgaptmp_002765', 'pgaptmp_002798', 'pgaptmp_002799', 'pgaptmp_002800', 'pgaptmp_002801', 'pgaptmp_002802', 'pgaptmp_002924', 'pgaptmp_002925', 'pgaptmp_002926', 'pgaptmp_002974', 'pgaptmp_003081', 'pgaptmp_003180', 'pgaptmp_003281', 'pgaptmp_003284', 'pgaptmp_003382', 'pgaptmp_003383', 'pgaptmp_003384', 'pgaptmp_003385', 'pgaptmp_003386', 'pgaptmp_003387', 'pgaptmp_003388', 'pgaptmp_003451',

'pgaptmp_003662', 'pgaptmp_003886', 'pgaptmp_003887', 'pgaptmp_003888',
'pgaptmp_003889', 'pgaptmp_003890', 'pgaptmp_003891', 'pgaptmp_003925',
'pgaptmp_003926', 'pgaptmp_003927', 'pgaptmp_003928', 'pgaptmp_003980',
'pgaptmp_003981', 'pgaptmp_003982', 'pgaptmp_003983', 'pgaptmp_003984',
'pgaptmp_004098', 'pgaptmp_004099', 'pgaptmp_004100', 'pgaptmp_004101',
'pgaptmp_004105', 'pgaptmp_004106', 'pgaptmp_004107', 'pgaptmp_004108',
'pgaptmp_004137', 'pgaptmp_004138', 'pgaptmp_004139', 'pgaptmp_004140',
'pgaptmp_004268', 'pgaptmp_004299', 'pgaptmp_004300', 'pgaptmp_004301',
'pgaptmp_004405', 'pgaptmp_004512', 'pgaptmp_004513', 'pgaptmp_004514', 'trna-bipA']
