## Supplementary File S5 for "Expanded ME-model analysis capabilities enable genetic recoding, proteostasis, RNA decay, and cofactor essentiality in bacterial models"

November 3, 2026

[1]:
```
import itables
itables.init_notebook_mode(all_interactive=True)
```

```
<IPython.core.display.HTML object>
```

```
<IPython.core.display.HTML object>
```

[2]:
```
import sys
sys.path.insert(0, './Supplementary-File-S8')
import coralme
coralme.__file__.split('Supplementary-File-S8')[1]
```

[2]:
```
'/coralme/__init__.py'
```

[3]:
```
import coralme
model = coralme.io.pickle.load_pickle_me_model('iMmy281-ME.pkl')
model.optimize()
```

```
Iteration       Solution to check       Solver Status
---------       ------------------      -------------
        1       1.4050280687025918      Not feasible
        2       0.7025140343512959      Not feasible
        3       0.3512570171756479      Not feasible
        4       0.1756285085878240      Not feasible
        5       0.0878142542939120      Optimal
        6       0.1317213814408680      Not feasible
        7       0.1097678178673900      Optimal
        8       0.1207445996541290      Not feasible
        9       0.1152562087607595      Not feasible
       10       0.1125120133140747      Optimal
       11       0.1138841110374171      Optimal
       12       0.1145701598990883      Not feasible
       13       0.1142271354682527      Not feasible
       14       0.1140556232528349      Not feasible
       15       0.1139698671451260      Optimal
       16       0.1140127451989804      Optimal
       17       0.1140341842259077      Not feasible
       18       0.1140234647124441      Optimal
       19       0.1140288244691759      Optimal
```

```
20        0.1140315043475418      Optimal
21        0.1140328442867247      Optimal
22        0.1140335142563162      Not feasible
```

[3]:
```
True
```

[4]:
```
model
```

[4]:
```
<MEModel mycoplasmaME at 0x702813d1b9e0>
```

### 0.0.1 Single-cofactor essentiality analysis

[5]:
```
# 4.5 minutes
res1 = coralme.util.essentiality.single_cofactor_essentiality_analysis(model)
```

```
100%|      | 18/18 [04:38<00:00, 15.45s/it]
```

[6]:
```
for value in res1.values():
    value.pop('rxns', None)

pandas.DataFrame(res1).T
```

[6]:
```
                                status
ca2_c         Can grow without cofactor
cd2_c         Can grow without cofactor
cobalt2_c     Can grow without cofactor
cs_c          Can grow without cofactor
fad_c       Can't grow without cofactor
fe2_c       Can't grow without cofactor
k_c         Can't grow without cofactor
lipoyl_c      Can grow without cofactor
mg2_c       Can't grow without cofactor
mn2_c       Can't grow without cofactor
na1_c         Can grow without cofactor
nad_c       Can't grow without cofactor
nh4_c         Can grow without cofactor
ni2_c         Can grow without cofactor
pi_c        Can't grow without cofactor
thmpp_c     Can't grow without cofactor
tl_c          Can grow without cofactor
zn2_c       Can't grow without cofactor
```

[7]:
```
# essential metabolites as enzyme cofactor
rxns, status = coralme.util.essentiality.test_cofactor_essentiality(model,
 ↪model.metabolites.fad_c)
status
```

[7]:
```
"Can't grow without cofactor"
```

[8]:
```
# non-essential metabolites as enzyme cofactor
rxns, status = coralme.util.essentiality.test_cofactor_essentiality(model,␣
↪model.metabolites.ca2_c)
status
```

[8]:
```
'Can grow without cofactor'
```

[9]:
```
import warnings
from IPython.core.formatters import FormatterWarning
warnings.filterwarnings('ignore', category = FormatterWarning)

# get reactions of a metabolite as enzyme cofactor
model.metabolites.lipoyl_c.as_enzyme_cofactor()
```

[9]:
```
MappableList([<ComplexData PDH_1_CPLX_mod_thmpp(1)_mod_lipoyl(1)_mod_fad(1) at
0x70274a9ebef0>,
 <ComplexData PDH_1_CPLX_mod_thmpp(1)_mod_lipoyl(1)_mod_fad(1)_lipoyl_scavenging
at 0x70274a7bfb90>,
 <ComplexData PDH_2_CPLX_mod_thmpp(1)_mod_lipoyl(1)_mod_fad(1) at
0x70274a9eba70>,
 <ComplexData PDH_2_CPLX_mod_thmpp(1)_mod_lipoyl(1)_mod_fad(1)_lipoyl_scavenging
at 0x70274a7bfc80>])
```

### 0.0.2 Single-gene essentiality analysis

[10]:
```
# ~45 seconds
res_m = coralme.util.essentiality.single_gene_essentiality_analysis(model.gem)
```

```
100%|      | 149/149 [02:46<00:00,  1.11s/it]
```

[11]:
```
# ~1.0 hours
res_me = coralme.util.essentiality.single_gene_essentiality_analysis(model,␣
↪debug = False)
# debug will save a pickle file for every single gene knockout model
```

```
100%|      | 281/281 [1:14:16<00:00, 15.86s/it]
```

[12]:
```
# single gene
test = coralme.util.essentiality.perform_gene_knockouts(model,␣
↪genes=['JCVISYN3A_0432'])
test.optimize() # use test.feasibility() to check a certain growth rate,␣
↪default 0.01
```

```
Iteration       Solution to check       Solver Status
---------       ------------------      -------------
        1       1.4050280687025918      Not feasible
        2       0.7025140343512959      Not feasible
        3       0.3512570171756479      Not feasible
        4       0.1756285085878240      Not feasible
```

```
         5       0.0878142542939120      Not feasible
         6       0.0439071271469560      Not feasible
         7       0.0219535635734780      Not feasible
         8       0.0109767817867390      Not feasible
         9       0.0054883908933695      Not feasible
        10       0.0027441954466847      Not feasible
        11       0.0013720977233424      Not feasible
        12       0.0006860488616712      Not feasible
        13       0.0003430244308356      Not feasible
        14       0.0001715122154178      Not feasible
        15       0.0000857561077089      Not feasible
        16       0.0000428780538544      Not feasible
        17       0.0000214390269272      Not feasible
        18       0.0000107195134636      Not feasible
        19       0.0000053597567318      Not feasible
        20       0.0000026798783659      Not feasible
        21       0.0000013399391830      Not feasible
        22       0.0000006699695915      Not feasible
```

[12]: False

[13]:
```
# multiple genes
test = coralme.util.essentiality.perform_gene_knockouts(model,␣
 ↪genes=['JCVISYN3A_0227', 'JCVISYN3A_0228'])
test.optimize() # use test.feasibility() to check a certain growth rate,␣
 ↪default 0.01
```

```
Iteration        Solution to check       Solver Status
---------        ------------------      -------------
        1        1.4050280687025918      Not feasible
        2        0.7025140343512959      Not feasible
        3        0.3512570171756479      Not feasible
        4        0.1756285085878240      Not feasible
        5        0.0878142542939120      Optimal
        6        0.1317213814408680      Not feasible
        7        0.1097678178673900      Optimal
        8        0.1207445996541290      Not feasible
        9        0.1152562087607595      Not feasible
       10        0.1125120133140747      Not feasible
       11        0.1111399155907324      Not feasible
       12        0.1104538667290612      Optimal
       13        0.1107968911598968      Not feasible
       14        0.1106253789444790      Optimal
       15        0.1107111350521879      Not feasible
       16        0.1106682569983334      Optimal
       17        0.1106896960252606      Optimal
       18        0.1107004155387243      Optimal
       19        0.1107057752954561      Optimal
```

```
        20       0.1107084551738220      Optimal
        21       0.1107097951130049      Optimal
        22       0.1107104650825964      Optimal
```

[13]: True

[14]:
```
dict_res_me = { k.id.removeprefix('RNA_'):v for k,v in res_me }
dict_res_m = { k.id:v for k,v in res_m }

df = pandas.concat([pandas.Series(dict_res_me), pandas.Series(dict_res_m)],␣
 ↪axis = 1)
df.columns = ['me', 'm']

# save predictions
df.sort_values("m").to_csv("gene_essentility_analysis.csv")

# plot results
from upsetplot import UpSet, from_indicators
warnings.filterwarnings('ignore', category = FutureWarning, module =␣
 ↪'upsetplot')

df = pandas.read_csv('gene_essentility_analysis.csv', index_col = 0)

data = pandas.DataFrame({
    'ME-model essential genes': df['me'].eq(True),
    'ME-model non-essential genes': df['me'].eq(False),
    'M-model essential genes': df['m'].eq(True),
    'M-model non-essential genes': df['m'].eq(False)
}, index = df.index)

upset_data = from_indicators(data.columns, data)

upset = UpSet(upset_data, subset_size = 'count', show_counts = True, sort_by =␣
 ↪'cardinality')

upset.plot()
plt.savefig('essentiality_upset.svg', format = 'svg', bbox_inches = 'tight')
plt.show()
```

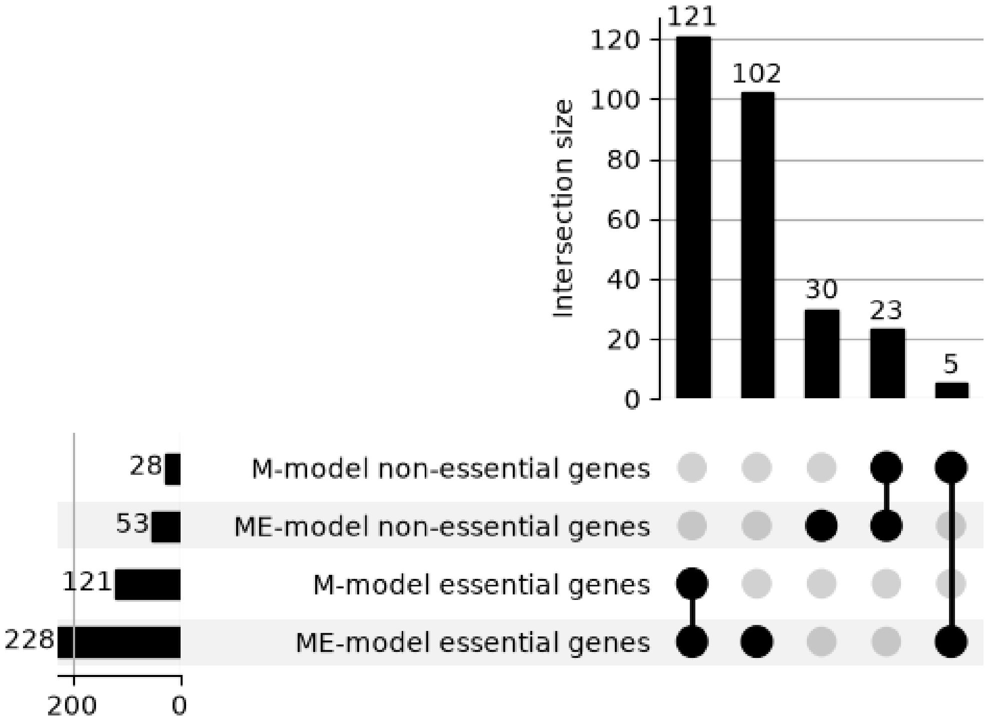
Intersection size
121
102
30
23
5
0
20
40
60
80
100
120
28
53
121
228
M-model non-essential genes
ME-model non-essential genes
M-model essential genes
ME-model essential genes
200
0
