## Supplementary File S6 for "Expanded ME-model analysis capabilities enable genetic recoding, proteostasis, RNA decay, and cofactor essentiality in bacterial models"

November 3, 2026

### 1 Examples: `single_cofactor_deletion` in CORALME

This notebook shows three common use cases for the cofactor deletion function:

1. Test a **single cofactor**.
2. Test a **user-defined list of cofactors**.
3. Test **all reaction-associated cofactors** in the model.

The API follows the style of COBRApy `single_gene_deletion` and returns a `pandas.DataFrame` with:

- `ids`
- `growth`
- `status`

> **Important:** when several cofactors are supplied in `cofactor_list`, each cofactor is deleted **independently** from the wild-type model. This is not a simultaneous multi-cofactor knockout.

#### 1.1 1. Environment and project paths

[1]:
```
import sys
sys.path.insert(0, './Supplementary-File-S8')
import coralme
coralme.__file__.split('Supplementary-File-S8')[1]
```

[1]: '/coralme/__init__.py'

[2]:
```
import sys
from pathlib import Path

import cobra
import coralme
import pandas as pd

print("Python :", sys.version.split()[0])
print("COBRApy:", cobra.__version__)
print("CORALME:", coralme.__version__)
print("CORALME path:", coralme.__file__)
```

```
Python : 3.12.14
COBRApy: 0.29.0
CORALME: 0+untagged.18.gd7f6581.dirty
CORALME path: /home/glucksfall/backup/Filen.io/supp-coralme-v1.3/Supplementary-
File-S8/coralme/__init__.py
```

[3]:
```
PROJECT = Path("./")
SRC = PROJECT

EXCEL_FILE = PROJECT / "iML1515_cofactor_rules.xlsx"

if str(SRC) not in sys.path:
    sys.path.insert(0, str(SRC))

# from cofactor_deletion
from coralme.util.essentiality    import (
    single_cofactor_deletion,
    constrained_reactions,
)

print("Excel file:", EXCEL_FILE)
print("Exists    :", EXCEL_FILE.exists())
```

```
Excel file: iML1515_cofactor_rules.xlsx
Exists    : True
```

##### 1.2 2. Load the cofactor-aware model

[4]:
```
cobra_model = coralme.util.excel2model.FromExcel(EXCEL_FILE)

model = coralme.core.model.MEModel.from_cobra(
    cobra_model
)

print("Model type :", type(model))
print("Reactions  :", len(model.reactions))
print("Metabolites:", len(model.metabolites))
```

```
INFO: genes are added from the 'reactions' spreadsheet, and gene annotations
from the 'genes' spreadsheet.
Model type : <class 'coralme.core.model.MEModel'>
Reactions  : 2775
Metabolites: 1877
```

##### 1.3 3. Example 1 — test one cofactor

A single cofactor can be supplied either as a string or as a one-element list. For clarity, the examples below use a list.

[5]:
```python
one_cofactor = ["mg2"]

result_one = single_cofactor_deletion(
    model,
    cofactor_list=one_cofactor,
    method="fba",
    processes=1,
    verbose=False,
)

result_one
```

[5]:

| | ids | growth | status |
|---|---|---|---|
| 0 | {mg2} | 9.573933e-32 | optimal |

[6]:
```python
# Optional: show how many reactions are disabled by this deletion.

cofactor = one_cofactor[0]

n_constrained = len(
    constrained_reactions(
        model,
        {cofactor},
    )
)

print("Cofactor              :", cofactor)
print("Constrained reactions :", n_constrained)
```

```
Cofactor              : mg2
Constrained reactions : 248
```

##### 1.4 4. Example 2 — test a user-defined list

Each entry in `cofactor_list` is tested as a separate single deletion from the wild-type model.

[7]:
```python
selected_cofactors = [
    "mg2",
    "fe2",
    "zn2",
    "pheme",
    "2fe2s",
]

result_selected = single_cofactor_deletion(
    model,
    cofactor_list=selected_cofactors,
    method="fba",
```

```
    processes=1,
    verbose=False,
)

result_selected
```

[7]:

| | ids | growth | status |
|---|---|---|---|
| 0 | {mg2} | 9.573933e-32 | optimal |
| 1 | {fe2} | 8.769973e-01 | optimal |
| 2 | {zn2} | 1.313886e-32 | optimal |
| 3 | {pheme} | 6.263490e-01 | optimal |
| 4 | {2fe2s} | 7.104562e-01 | optimal |

[8]:

```
# Add a compact summary with the number of constrained reactions.

selected_summary = result_selected.copy()

selected_summary["Cofactor"] = selected_summary["ids"].apply(
    lambda ids: next(iter(ids))
)

selected_summary["ConstrainedReactions"] = (
    selected_summary["Cofactor"].apply(
        lambda cofactor: len(
            constrained_reactions(
                model,
                {cofactor},
            )
        )
    )
)

selected_summary = selected_summary[
    [
        "Cofactor",
        "growth",
        "status",
        "ConstrainedReactions",
    ]
]

selected_summary
```

[8]:

| | Cofactor | growth | status | ConstrainedReactions |
|---|---|---|---|---|
| 0 | mg2 | 9.573933e-32 | optimal | 248 |
| 1 | fe2 | 8.769973e-01 | optimal | 7 |
| 2 | zn2 | 1.313886e-32 | optimal | 36 |

```
3    pheme  6.263490e-01  optimal                        7
4    2fe2s  7.104562e-01  optimal                        9
```

##### 1.5 5. Example 3 — test all associated cofactors

If `cofactor_list` is omitted, `single_cofactor_deletion` automatically tests all cofactors that occur in at least one reaction cofactor rule.

For this iML1515-derived model, that means **47 associated cofactors**.

[9]:
```python
result_all = single_cofactor_deletion(
    model,
    method="fba",
    processes=1,
    verbose=False,
)

print("Number of single-cofactor deletions:", len(result_all))

result_all
```

```
Number of single-cofactor deletions: 47
```

[9]:

| | ids | growth | status |
|---|---|---|---|
| 0 | {2fe2s} | 7.104562e-01 | optimal |
| 1 | {3fe4s} | 8.418159e-01 | optimal |
| 2 | {4fe4s} | -9.788960e-32 | optimal |
| 3 | {NiFe_cofactor} | 8.769973e-01 | optimal |
| 4 | {adocbl} | 8.769973e-01 | optimal |
| 5 | {bmocogdp} | 8.769973e-01 | optimal |
| 6 | {btn} | 3.690056e-52 | optimal |
| 7 | {ca2} | 8.769973e-01 | optimal |
| 8 | {cbl1} | 8.769973e-01 | optimal |
| 9 | {cd2} | 8.769973e-01 | optimal |
| 10 | {cl} | 8.769973e-01 | optimal |
| 11 | {cobalt2} | 8.648569e-54 | optimal |
| 12 | {cs} | 8.769973e-01 | optimal |
| 13 | {cu} | 8.769973e-01 | optimal |
| 14 | {cu2} | 7.776144e-01 | optimal |
| 15 | {dpm} | -1.476022e-51 | optimal |
| 16 | {fad} | 0.000000e+00 | optimal |
| 17 | {fadh2} | 8.769973e-01 | optimal |
| 18 | {fe2} | 8.769973e-01 | optimal |
| 19 | {fe3} | 8.769973e-01 | optimal |
| 20 | {fmn} | 6.387131e-28 | optimal |
| 21 | {fmnh2} | 8.769973e-01 | optimal |
| 22 | {glycyl} | 8.769973e-01 | optimal |
| 23 | {hco3} | 8.769973e-01 | optimal |

```
24           {hemeO}  7.776144e-01  optimal
25           {hemed}  8.769973e-01  optimal
26               {k} -2.865640e-30  optimal
27              {li}  8.769973e-01  optimal
28          {lipoyl}  8.408972e-01  optimal
29          {lys__L}  8.769973e-01  optimal
30             {mg2}  9.573933e-32  optimal
31             {mn2}  0.000000e+00  optimal
32             {na1}  8.769973e-01  optimal
33             {nad}  1.153142e-53  optimal
34            {nadh}  0.000000e+00  optimal
35            {nadp}  8.769973e-01  optimal
36           {nadph}  0.000000e+00  optimal
37             {nh4}  8.769973e-01  optimal
38             {ni2}  8.769973e-01  optimal
39           {pheme}  6.263490e-01  optimal
40              {pi}  8.769973e-01  optimal
41             {pqq}  8.769973e-01  optimal
42          {pydx5p} -1.441428e-54  optimal
43           {sheme}  8.769973e-01  optimal
44           {thmpp} -2.066431e-50  optimal
45              {tl}  8.769973e-01  optimal
46             {zn2}  1.313886e-32  optimal
```

[10]:

```
all_summary = result_all.copy()

all_summary["Cofactor"] = all_summary["ids"].apply(
    lambda ids: next(iter(ids))
)

all_summary["ConstrainedReactions"] = (
    all_summary["Cofactor"].apply(
        lambda cofactor: len(
            constrained_reactions(
                model,
                {cofactor},
            )
        )
    )
)

all_summary["HasEffect"] = (
    all_summary["ConstrainedReactions"] > 0
)

all_summary = all_summary[
    [
```

```
        "Cofactor",
        "growth",
        "status",
        "HasEffect",
        "ConstrainedReactions",
    ]
].sort_values("Cofactor").reset_index(drop=True)

all_summary
```

[10]:

| | Cofactor | growth | status | HasEffect | ConstrainedReactions |
|---|---|---|---|---|---|
| 0 | 2fe2s | 7.104562e-01 | optimal | True | 9 |
| 1 | 3fe4s | 8.418159e-01 | optimal | True | 5 |
| 2 | 4fe4s | -9.788960e-32 | optimal | True | 22 |
| 3 | NiFe_cofactor | 8.769973e-01 | optimal | True | 1 |
| 4 | adocbl | 8.769973e-01 | optimal | True | 2 |
| 5 | bmocogdp | 8.769973e-01 | optimal | True | 15 |
| 6 | btn | 3.690056e-52 | optimal | True | 1 |
| 7 | ca2 | 8.769973e-01 | optimal | True | 28 |
| 8 | cbl1 | 8.769973e-01 | optimal | False | 0 |
| 9 | cd2 | 8.769973e-01 | optimal | True | 7 |
| 10 | cl | 8.769973e-01 | optimal | True | 1 |
| 11 | cobalt2 | 8.648569e-54 | optimal | True | 21 |
| 12 | cs | 8.769973e-01 | optimal | False | 0 |
| 13 | cu | 8.769973e-01 | optimal | True | 3 |
| 14 | cu2 | 7.776144e-01 | optimal | True | 2 |
| 15 | dpm | -1.476022e-51 | optimal | True | 1 |
| 16 | fad | 0.000000e+00 | optimal | True | 40 |
| 17 | fadh2 | 8.769973e-01 | optimal | False | 0 |
| 18 | fe2 | 8.769973e-01 | optimal | True | 7 |
| 19 | fe3 | 8.769973e-01 | optimal | False | 0 |
| 20 | fmn | 6.387131e-28 | optimal | True | 15 |
| 21 | fmnh2 | 8.769973e-01 | optimal | False | 0 |
| 22 | glycyl | 8.769973e-01 | optimal | True | 2 |
| 23 | hco3 | 8.769973e-01 | optimal | True | 5 |
| 24 | hemeO | 7.776144e-01 | optimal | True | 1 |
| 25 | hemed | 8.769973e-01 | optimal | True | 2 |
| 26 | k | -2.865640e-30 | optimal | True | 16 |
| 27 | li | 8.769973e-01 | optimal | False | 0 |
| 28 | lipoyl | 8.408972e-01 | optimal | True | 3 |
| 29 | lys__L | 8.769973e-01 | optimal | True | 1 |
| 30 | mg2 | 9.573933e-32 | optimal | True | 248 |
| 31 | mn2 | 0.000000e+00 | optimal | True | 24 |
| 32 | na1 | 8.769973e-01 | optimal | False | 0 |
| 33 | nad | 1.153142e-53 | optimal | True | 4 |
| 34 | nadh | 0.000000e+00 | optimal | True | 2 |
| 35 | nadp | 8.769973e-01 | optimal | True | 2 |

| | | | | | |
|---|---|---|---|---|---|
| 36 | nadph | 0.000000e+00 | optimal | True | 2 |
| 37 | nh4 | 8.769973e-01 | optimal | True | 1 |
| 38 | ni2 | 8.769973e-01 | optimal | True | 4 |
| 39 | pheme | 6.263490e-01 | optimal | True | 7 |
| 40 | pi | 8.769973e-01 | optimal | True | 1 |
| 41 | pqq | 8.769973e-01 | optimal | True | 1 |
| 42 | pydx5p | -1.441428e-54 | optimal | True | 45 |
| 43 | sheme | 8.769973e-01 | optimal | True | 4 |
| 44 | thmpp | -2.066431e-50 | optimal | True | 9 |
| 45 | tl | 8.769973e-01 | optimal | False | 0 |
| 46 | zn2 | 1.313886e-32 | optimal | True | 36 |

##### 1.6 6. Inspect specific biological outcomes

[11]:
```
# Example: cofactors predicted to be lethal or nearly lethal.

LETHAL_TOL = 1e-9

lethal = all_summary[
    all_summary["growth"].abs() <= LETHAL_TOL
]

lethal
```

[11]:

| | Cofactor | growth | status | HasEffect | ConstrainedReactions |
|---|---|---|---|---|---|
| 2 | 4fe4s | -9.788960e-32 | optimal | True | 22 |
| 6 | btn | 3.690056e-52 | optimal | True | 1 |
| 11 | cobalt2 | 8.648569e-54 | optimal | True | 21 |
| 15 | dpm | -1.476022e-51 | optimal | True | 1 |
| 16 | fad | 0.000000e+00 | optimal | True | 40 |
| 20 | fmn | 6.387131e-28 | optimal | True | 15 |
| 26 | k | -2.865640e-30 | optimal | True | 16 |
| 30 | mg2 | 9.573933e-32 | optimal | True | 248 |
| 31 | mn2 | 0.000000e+00 | optimal | True | 24 |
| 33 | nad | 1.153142e-53 | optimal | True | 4 |
| 34 | nadh | 0.000000e+00 | optimal | True | 2 |
| 36 | nadph | 0.000000e+00 | optimal | True | 2 |
| 42 | pydx5p | -1.441428e-54 | optimal | True | 45 |
| 44 | thmpp | -2.066431e-50 | optimal | True | 9 |
| 46 | zn2 | 1.313886e-32 | optimal | True | 36 |

[12]:
```
# Example: cofactors that constrain reactions but do not reduce WT growth.

WT_GROWTH = result_all["growth"].max()

neutral_but_effective = all_summary[
    (all_summary["HasEffect"])
```

```
    & ((all_summary["growth"] - WT_GROWTH).abs() <= 1e-6)
]

neutral_but_effective
```

[12]:

| | Cofactor | growth | status | HasEffect | ConstrainedReactions |
|---|---|---|---|---|---|
| 3 | NiFe_cofactor | 0.876997 | optimal | True | 1 |
| 4 | adocbl | 0.876997 | optimal | True | 2 |
| 5 | bmocogdp | 0.876997 | optimal | True | 15 |
| 7 | ca2 | 0.876997 | optimal | True | 28 |
| 9 | cd2 | 0.876997 | optimal | True | 7 |
| 10 | cl | 0.876997 | optimal | True | 1 |
| 13 | cu | 0.876997 | optimal | True | 3 |
| 18 | fe2 | 0.876997 | optimal | True | 7 |
| 22 | glycyl | 0.876997 | optimal | True | 2 |
| 23 | hco3 | 0.876997 | optimal | True | 5 |
| 25 | hemed | 0.876997 | optimal | True | 2 |
| 29 | lys__L | 0.876997 | optimal | True | 1 |
| 35 | nadp | 0.876997 | optimal | True | 2 |
| 37 | nh4 | 0.876997 | optimal | True | 1 |
| 38 | ni2 | 0.876997 | optimal | True | 4 |
| 40 | pi | 0.876997 | optimal | True | 1 |
| 41 | pqq | 0.876997 | optimal | True | 1 |
| 43 | sheme | 0.876997 | optimal | True | 4 |

#### 1.7 7. Save results (optional)

[13]:

```
OUTPUT_DIR = PROJECT / "results" / "single_cofactor_examples"
OUTPUT_DIR.mkdir(parents=True, exist_ok=True)

selected_file = OUTPUT_DIR / "selected_cofactor_deletions.tsv"
all_file = OUTPUT_DIR / "all_single_cofactor_deletions.tsv"

selected_summary.to_csv(
    selected_file,
    sep="\t",
    index=False,
)

all_summary.to_csv(
    all_file,
    sep="\t",
    index=False,
)

print("Saved:")
print(" ", selected_file)
```

```
print(" ", all_file)
```

```
Saved:
  results/single_cofactor_examples/selected_cofactor_deletions.tsv
  results/single_cofactor_examples/all_single_cofactor_deletions.tsv
```
